# Flavin cycling under prebiotic conditions: bidirectional electron transfer and versatility in nickel and iron containing environments

**DOI:** 10.64898/2026.07.08.736930

**Authors:** Oskari Lehtinen, Delfina P. Henriques Pereira, Medhin T. Yasin, Nicole Paczia, Martina Preiner

## Abstract

Flavins are organic redox cofactors central to metabolism and uniquely capable of acting as extracellular electron shuttles. For life to have emerged, it must have disengaged itself from its stationary geochemical environment, a step requiring mobile redox-active components. The role of flavins at life’s origin has been debated for decades, centered on their capacity for both one- and two-electron chemistry, distinguishing them from nicotinamides and iron-sulfur clusters. Here we chart the abiotic reduction of flavin mononucleotide (FMN), flavin adenine dinucleotide (FAD), and riboflavin under hydrothermal conditions (40 °C, 1 bar N_2_ or 5 bar H_2_, pH 6, 8, and 10) by nickel (Ni) and iron (Fe). Flavins show greater environmental versatility than hydride carriers such as NAD and can harvest electrons from metals that would otherwise reduce water’s protons to H_2_. Reduction is favoured under acidic conditions, while increasing molecular charge at higher pH impedes electron transfer. Ni acts as a hydrogenation catalyst, reducing deprotonated flavins via hydride transfer, suggesting mineral composition could have influenced geochemical selection of early electron carriers. Reduced FMNH_2_ and FADH_2_ were tested as electron shuttles toward Fe^3+^-containing minerals, revealing that FMNH_2_ enables faster mineral dissolution than FADH_2_. We further demonstrate complete redox cycling of FMN through Ni-assisted H_2_ reduction and subsequent oxidation by magnetite (Fe_3_O_4_) under inert atmosphere, releasing Fe^2+^. This study highlights the versatility, stability and redox chemical capabilities of flavins in prebiotic context.

**Significance statement:** Here we show how flavins, among the most evolutionarily conserved organic cofactors in biology, interact with metal surfaces under prebiotic hydrothermal conditions, and how geochemical parameters such as metal abundance and pH influence their ability to accept electrons from the environment. Flavins could serve as electron shuttles in geochemical systems that might otherwise lose their reducing power to the volatile hydrogen gas. More than that, flavins cycle through reduction and oxidation under conditions relevant to mineral-containing hydrothermal sites on the early Earth, linking the motile electron-carrying component of prebiotic chemical networks with immobile mineral environments.

## Introduction

Flavins and derivatives thereof have been in focus of pre-enzymatic hypotheses of life’s emergence for decades^[1–8]^ as connectors between RNA world hypotheses and protometabolic processes (“nucleotide-derived”), either as parts of protometabolic electron transport chains or as photocatalysts^[7,9]^. Phylogenetic reconstructions confirm the presence of flavins (FAD, FMN, Riboflavin, **Fig. 1 A**) in the last universal common ancestor (LUCA) ^[10– 12]^. Also prebiotic syntheses of flavins were shown via a variety of routes^[13,14]^.

**Figure 1:**
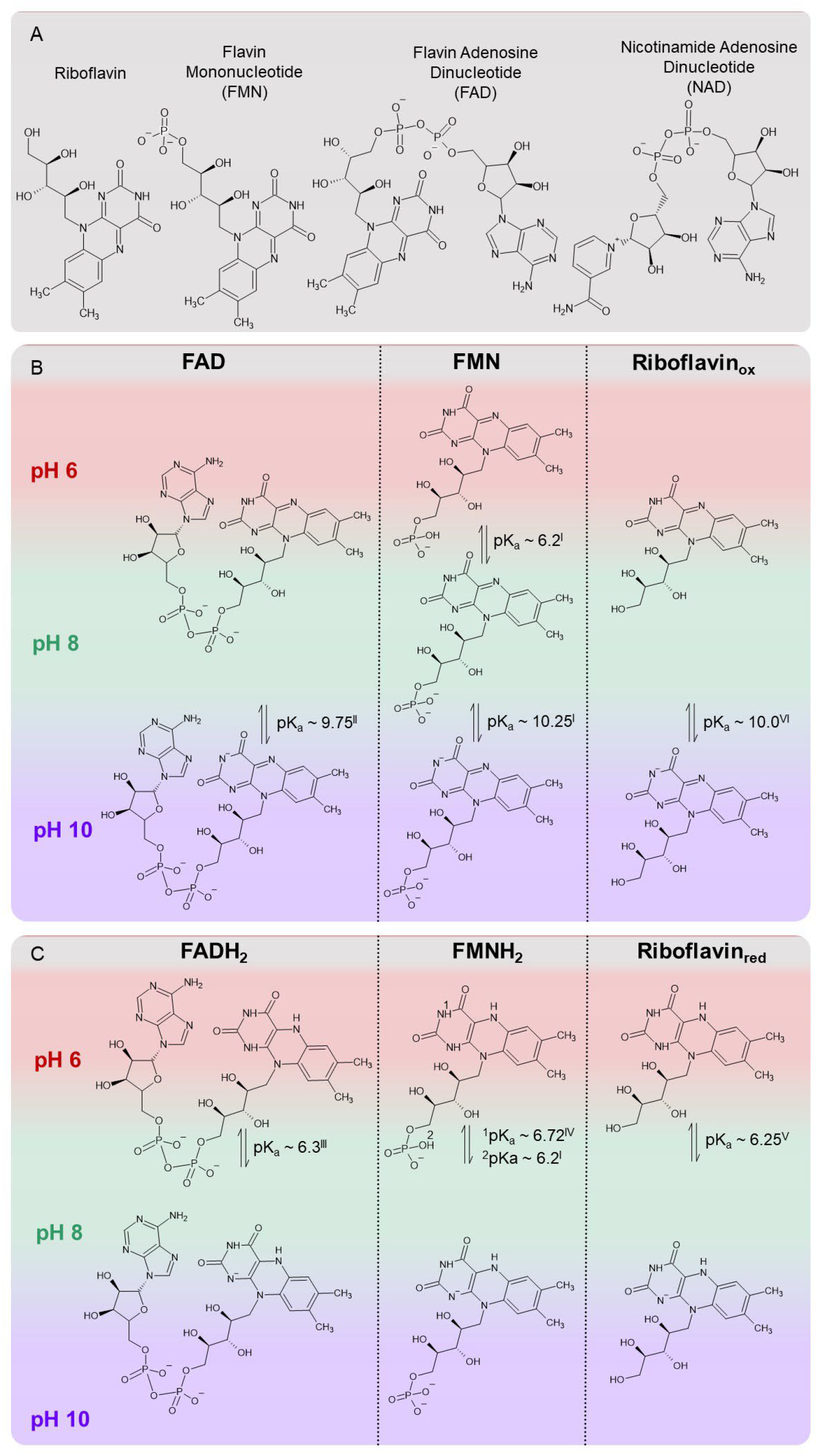
The flavins used in this study and their protonation states. A) Overview of the most common organic redox cofactors discussed in this study. While flavins enable one and two electron transfers, deazaflavins and nicotinamides are hydride carries. B) pKa values of cofactors investigated in this study in their oxidized state under the tested conditions. C) pKa values of cofactors investigated in this study in their reduced state under the tested conditions. pKa values: ^I^ from ref.^[32]^; ^II^ from ref.^[33]^; ^III^ from ref.^[34]^, ^IV^ from ref.^[35]^, ^V^ from ref.^[36]^, ^VI^ from ref.^[37]^.

In extant biology, flavins take over a myriad of tasks in cells: metabolic redox reactions, DNA repair, extracellular electron shuttling, bioluminescence, blue light sensing, cofactor synthesis and more^[15–17]^. When it comes to metabolic redox reactions, flavins are capable of performing two-electron reactions, such as nicotinamide cofactors (NAD, NADP, nicotine amide mononucleotide), but flavins can also do single (radical) electron transfer like iron sulphur clusters ^[18]^. That ability makes flavins essential to a variety of biochemical reactions, the most ground-breaking discovery being that of flavin-based electron bifurcation^[19]^. During this process, hydride electron pairs (as e.g. H_2_) are split into an electron with a higher redox potential (e.g. NADH) and another with lower (e.g. Ferredoxin) than that of the original electron pair^[20]^. This mechanism is critical for organisms living at the “thermodynamic limit of life” (feedstock from CO_2_ reduction with hydrogen), as is the case of methanogens, acetogens, and therefore possibly LUCA^[21–23]^. Prebiotically, however, the role of bifurcation has been questioned^[24]^, although radical semiquinones can be stabilized abiotically^[25,26]^. It was also recently shown that an excess of Fe^0^ and H_2_ can reduce low-potential ferredoxin directly, indicating that electron bifurcation would not be necessary to reduce ferredoxin in a very reducing prebiotic environment, such as water-rock interaction (serpentinizing) systems^[27]^. The role flavins could have had in such non-enzymatic bifurcation processes still needs to be investigated.

In a microbial context, flavins also function as electron shuttles, to transport electrons to extracellular Fe^3+[28]^. As far as this process is understood, it serves both to find terminal electron acceptors and to help transform insoluble Fe^3+^ to soluble Fe^2+^, which subsequently can either be reoxidized by iron oxidizers or assimilated^[29]^. The opposite direction of electron shuttling – taking electrons from either Fe^2+^ or Fe^0^ via flavins – has also been proposed^[30]^, but is still debated ^[31]^.

Apart from the already mentioned roles of flavins, these and other cofactors can be tested as parts of non-enzymatic protometabolic networks. In such, they could have enabled transport of electrons, transport of metal ions and possibly even drive synthesis of other central biomolecules due to its unique redox chemistry^[38,39]^.

Recently, FAD has been used successfully within a non-enzymatic electron transport chain from a ketoacid via NAD and FAD to a final electron acceptor^[4]^. In this study, we test flavins’ ability to shuttle electrons from H_2_ and transition metals such as nickel (Ni) and iron (Fe) beyond the context of biological systems and in perspective of the origin of life. These two metals are both found in H_2_-rich geochemical environments such as serpentinzing systems, and are also the central metals in life’s most ancient hydrogenase, and thus central to make the electrons of H_2_ accessible to early autotrophic metabolism^[40]^. Especially Ni has lately been in focus as a generally bridging transition metal between geo- and biochemistry^[41]^. Furthermore, the influence of pH on reduction yields is investigated. From electrochemical studies with flavins it is known that their protonation and deprotonation patterns (pK_a_) directly affect their reducibility (**Fig. 1 B and C**)^[32]^.

We simultaneously monitor metal ion formation during the reactions and their effect on the overall stability of different flavins. H_2_ formation from the metals with and without cofactors shows that flavins take up electrons that would otherwise reduce the protons of water. Lastly, we compare the ability of both FADH_2_ and FMNH_2_ to reduce and dissolve Fe^3+^ from iron oxides to evaluate the overall non-enzymatic abilities of both cofactors.

## Results and discussion

FAD and FMN’s reduction were tested under both H_2_ and inert atmosphere with a variety of metals and minerals (4 h, 40°C; ***SI Appendix*, Tab. S1**). The results of these exploratory reactions show that magnetite (Fe_3_O_4_) and nickel oxide (NiO) were not able to promote reduction to FADH_2_ under the tested conditions (***SI Appendix*, Fig. S5**). Native metals such as nickel (Ni^0^) and iron (Fe^0^) can assist in the reduction of both flavins and so does pyrite (FeS) with far lower yields **(*SI Appendix*, Fig. S5 C)**. Based on this, a more detailed screening with Ni and Fe nanopowder (Ni: 100 nm, Sigma; Fe: 60–70 nm, Nanografi; metal to cofactor ratio 4.5:1) at three different pH values (6, 8 and 10) under either H_2_ (5 bar) or N_2_ (1 bar) at 40 °C was performed. The aim was to investigate different electron transfer mechanisms, based on our previous work with another redox cofactor, nicotinamide adenosine dinucleotide (NAD)^[42,43]^. The starting concentration of oxidized flavins was 4 mM (4 μmol in 1 mL buffer), the metal powders were 1 mg (18 μmol of Ni or Fe). The concentration of H_2_ resulting from 5 bar pressure under the given reaction conditions is about 3.5 mM, calculated previously using Henry’s law^[10]^. This value is comparable with H_2_ concentrations found in extant serpentinizing systems^[44,45]^, but can also be transferred to more general reducing geochemical settings. The calculated redox potentials for 3.5 mM H_2_ under the different pH values are: E(pH 6) = –395 mV; E(pH 8) = –519 mV; E(pH 10) = –643 mV (***SI Data* 1**). We determined the concentrations of reduced and oxidized flavins at two time points (0.25 h and 2 h) via UV-Vis spectroscopy and measured pHs after reactions to ensure their stability during the reaction (**Fig. 2 A and *SI Appendix*, Table S21 and Fig. S3**). back into solution (***SI Appendix*, Figs. S19–21 and Table S34**). All reactions were performed in triplicate. C) Metal ion concentrations after 2h reactions measured via ICP-MS (***SI Appendix*, Table S24**). In Ni samples, concentrations of Ni^2+^ are depicted, in Fe samples, the concentrations of both Fe^2+^ and Fe^3+^. Additional ferrozine tests showed that the majority of dissolved ion ions is in the Fe^2+^ state (***SI Appendix*, Table S27 and Fig. S9**). 2 h controls with flavin-free buffer over the two metals show either similar concentrations as with FAD or lower (***SI Appendix*, Table S24**). Especially in the case of Ni, a higher concentration of ions correlates with a higher amount of flavin precipitation and loss. D) Proposed mechanistic difference between Ni and Fe promoted reduction at pH 10. The flavin being deprotonated makes it harder to accept electrons directly. However, direct hydride transfer seems possible with Ni. E) The concentrations of reduced riboflavin in solution after a 2 h reaction with Ni and Fe under 5 bar of H_2_ gas or 1 bar of N_2_ gas, at three different pHs – quantified via UV-Vis as described in SI methods and ***SI Appendix*, Figs. S2, Tabs. S2 and 15–20, Eqs. S1–5**. Dotted lines indicate the 100% yield mark at a concentration of 100 μM, the starting concentration of the oxidised flavins. All reactions were performed in triplicate.

**Figure 2:**
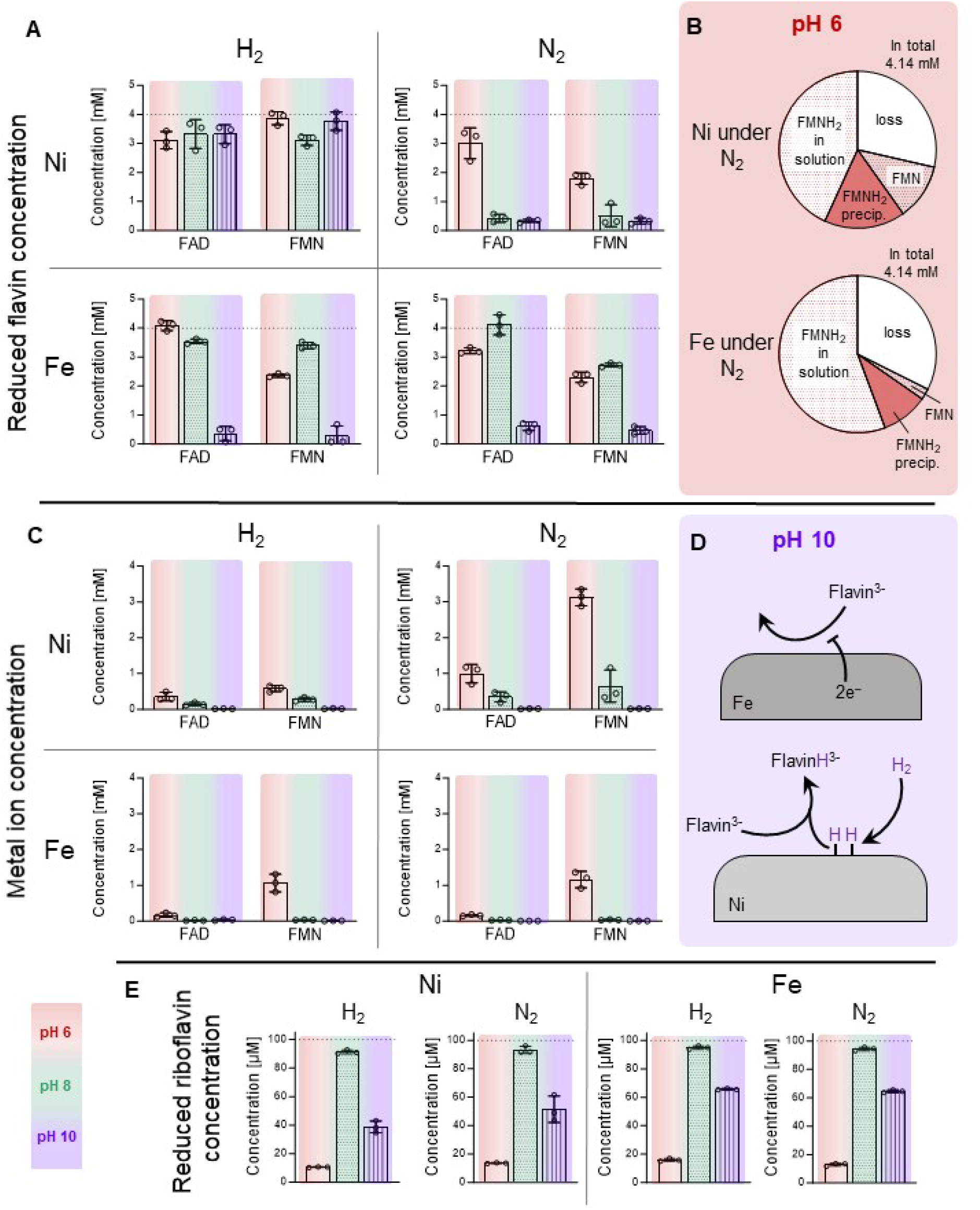
Metal-assisted flavin reduction under different pHs. A) The concentrations of reduced flavins in solution (FADH_2_ and FMNH_2_) after a 2 h reaction with Ni or Fe under 5 bar of H_2_ gas or 1 bar of N_2_ gas, at three different pHs – quantified via UV-Vis as described in SI methods and ***SI Appendix*, Figs S2, Table S2–14, Eqs. S1–5**. Dotted lines indicate the 100% yield mark at a concentration of 4 mM, the starting concentration of the oxidized flavins. An additional time point (0.25 h) was taken, showing that a lot of the reduction processes happen fast (***SI Appendix*, Table S21 and Fig. S3**). All reactions were performed in triplicate. B) Overview of flavin losses during reaction for pH 6 with Ni and Fe. In the case of Ni, FMN measureable precipitates during the reaction and work up, but can be shaken

The reduction results show a clear pattern: at acidic pH, both FAD and FMN are readily reduced, regardless of the gas phase (**Fig. 2 A**). At pH 8, Ni does not significantly reduce either flavin without H_2_ atmosphere, while Fe reactions yield more than 3 mM of reduced flavin under both N_2_ and H_2_. Under pH 10, however, only the combination of Ni with H_2_ gas achieves almost 100% yield. Controls without Ni or Fe (***SI Appendix*, Figs. S22 and S23**) clearly show that metals are necessary for the reduction to happen.

This is in line with existing literature on the electrochemical reduction of flavins and stems from the pK_a_ values of both flavins^[32]^. First, the flavins’ phosphate groups and then the isoalloaxine ring deprotonate with increasing pH (**Fig. 1 B**), which increases their negative charge and consequently decreases the flavins’ redox potential, i.e. they are harder to reduce and easier to oxidize. (***SI Appendix*, Fig. S6**). We confirmed this trend also by performing cyclic voltammetry in our according buffers (***SI Appendix*, Fig. S7**).

So if high pH hinders flavin reduction, why does the combination of Ni with H_2_ work? A direct transfer of electrons from the metals is, in this case, unlikely, given that the isoalloaxine ring is already negatively charged and the reduction also requires two protons, which are few at pH 10. Ni, however, is a known hydrogenation catalyst ^[46]^, i.e. it can cleave H_2_ on its surface and transfer the resulting hydride, in this case onto FAD and FMN (**Fig. 2 D**), providing electrons and protons at the same time. This shows that environmental conditions such as pH, metal abundance, and distribution, could have a major effect on the functionality of flavins in a prebiotic scenario. In other words, under H_2_-rich conditions, Ni could assist flavin reduction under both acidic, neutral and alkaline conditions. This pH flexibility and stability are remarkable, especially compared to NAD that quickly becomes unstable under acidic or alkaline conditions^[42,47]^.

We also tested the unphosphorylated riboflavin under the same experimental conditions, for 2 h reaction time (**Fig. 2 E**). Since riboflavin is far less soluble than its phosphorylated derivatives, the starting concentration of oxidized riboflavin was 100 μM, resulting in a far higher metal to coenzyme ratio than for FAD and FMN experiments (180:1). Under all conditions, riboflavin gets reduced readily, even at pH 10, which differs from FAD and FMN with the exception of Ni under H_2_ gas. One explanation is that riboflavin is far less negatively charged than either FAD or FMN under pH 8 or pH 10 – so direct electron transfer from metal to coenzyme is more facile. Another one is that the amount of coenzyme reduced in this reaction is a lot lower (100 μM) than in the FMN and FAD experiments. Even though the efficiency of reducing FAD and FMN at pH 10 is significantly smaller, compared to other pHs, more than 100 μM are reduced.

We monitored the changes of the used metal powders via Micro-X-ray fluoroscence (μXRF) and visual observations such as the formation of blue vivianite: an iron phosphate complex (***SI Appendix*, Fig. S17 and Table S33)**. At pH 6 and pH 8 an association with phosphate and the produced metal ions is likely, while at pH10 the formation of hydroxides likely outcompetes the formation of metal carbonates^[48–50]^.

### Counting the losses

After determining the concentration of reduced flavin, reoxidation of the samples was performed to account for losses. For most conditions, this aligns with the concentration of a metal-free flavin control (4 mM) (***SI Appendix*, Figs. S22 and S23, and Table S21**). At pH 6, especially for reactions of FMN under N_2_, a lower concentration of flavins is detected after re-oxidation than in the control (**Fig. 2 B**). At the same time FMN samples show a visible red pellet in Ni and Fe experiments after the 4 °C centrifugation (***SI Appendix*, Fig. S19)**, with the first precipitation starting in the reaction vial as soon as the stirring stops. Anaerobic resuspension of the pellet shows that between 0.4 and 0.7 mM of FMNH_2_ (10–17.5 % of the starting FMN concentration) fell out of solution that way (***SI Appendix*, Fig. S21, Table S34**).

The ion concentrations (**Fig. 2 C**) after each reaction were measured via Inductively Coupled Plasma Mass Spectrometry (ICP-MS) and compared to flavin-free controls to reveal several trends. First, more ions go into solution when cofactors are present. Second, the FMN samples show higher ion concentrations than the FAD samples, although the reduction yields are comparable and thus the resulting oxidation of the metals should be similar. FMN forms metal-complexes faster than FAD^[51]^. This is due to the structural differences between FAD and FMN. In FAD, the AMP moiety and the flavin moiety can interact, e.g. by forming intramolecular hydrogen bonding, and hinder stable metal complexation^[33,52–54]^. The ions concentrations after FMN reduction and cold centrifugation indicate that its interaction with metal ions outcompetes the formation of less soluble metal hydroxides or phosphates^[55,56]^. Third, the FMN reactions producing the most dissolved ions (under N_2_), suffer the most flavin precipitation and overall loss of signal after re-oxidation (***SI Appendix*, Figures S4, and Table S21 and S34**).

As literature describes the formation of Fe^2+^, Fe^3+^ and Ni^2+^ complexes with FMN^[52]^, self-association of FMN^[57]^, and photolysis of flavin-metal complexes^[58]^, we did not find a simple explanation for reduced FMN precipitating out of solution at acidic pH after reactions with metals. A well-documented observation, however, seems to be the formation of semiquinone-metal complexes with a red colouration^[59]^, which in the case of FMN can lead to precipitation^[26]^. Considering that under this study’s conditions (pH 6), semiquinone species could form when oxidized and reduced flavin exists at equal concentrations (50:50)^[25]^, it seems feasible that metal complexes can form and fall out of solution. Disproportionation of the semiquinone back to oxidized and reduced flavin could also explain the FMNH_2_ recovered from the pellet (**Fig. 2 B**).

We performed control experiments at pH 6 with FMNH_2_ and Ni^2+^, Fe^3+^, or Fe^2+^ salts that show that Ni^2+^ and Fe^3+^ indeed irreversibly decrease the UV-Vis absorption of the flavin (***SI Appendix*, Figs. S13 and S14**), and a red-brown pellet is formed with Ni^2+^ (***SI Appendix*, Figs. S15**). Starting with oxidized flavin, this effect cannot be observed, the UV-Vis spectrum remains the same as without Ni^2+^ (***SI Appendix*, Fig. S12 and S16**). When FMNH_2_ is mixed with Fe^2+^, no change can be observed. Fe^3+^, however, shows a major effect, including precipitation and oxidation of FMNH_2_, as Fe^3+^ is reduced to Fe^2+^ . It is hence probable that the ion formation during reaction triggers the pellet formation and thus causes a major part of the initial signal loss. The combination of metal-flavin interactions and overall decrease of solubility at low pHs can ultimately give a coherent explanation for the observed.

Also riboflavin shows a significant decrease in absorption at pH 6 as expected for reduction, but fails do re-oxidize back to the starting concentration (**Fig. 2 E, *SI Appendix*, Figs. S4, S20, and Table S34**). This loss could be due to declining solubility of reduced riboflavin at pH 6 (**Fig. 1 C**) or (pellet-forming) interactions with metals as observed with FMN (***SI Appendix*, Fig. S13 B and S14**)^[26,60]^. Retrieval of lost riboflavin was tried by pH increase after reaction to account for lower solubility at pH 6, with partial success (***SI Appendix*, Fig. S18**).

### Where the electrons come from and go

In a prebiotic environment which is constantly producing H_2,_ such as serpentinizing systems, what impact could the presence of flavins have? To assess the direction of electron flows in our systems, we determined the H_2_ generation by metals under different pHs with and without flavins at 1 bar of N_2_. Then, we compared the redox potential of H_2_ from water to the midpoint potentials of both metal oxidation and flavin reduction (**Fig. 3 A and B**). Based on these calculations and the observation that the H_2_ production significantly drops in the presence of flavins, the electron flow chart shows how flavins can non-enzymatically scavenge electrons from the environment (**Fig. 3 C**). As a result, far more electrons are bound in reduced flavin than in H_2_. Both, when protons and flavins are competing the same reaction, but also compared to the H_2_ formation without flavins in solution. In other words, flavins harvest electrons that otherwise would not even reduce the protons of water readily. Likely, the H_2_ generation observed with cofactors in solution (especially at pH 6) is the result of all dissolved flavin already being reduced or degraded, opening up the electron flow for protons again. Recently, it has been argued in the context of microbial electron shuttling in bio corrosion that Fe would always rather reduce the protons of water^[31]^. Although the authors righteously argue that the microorganisms in question do not use flavins as electron shuttles from extracellular Fe^0^ into the cell, the observation that protons will be rather reduced than flavins might not be universally applicable.

**Figure 3:**
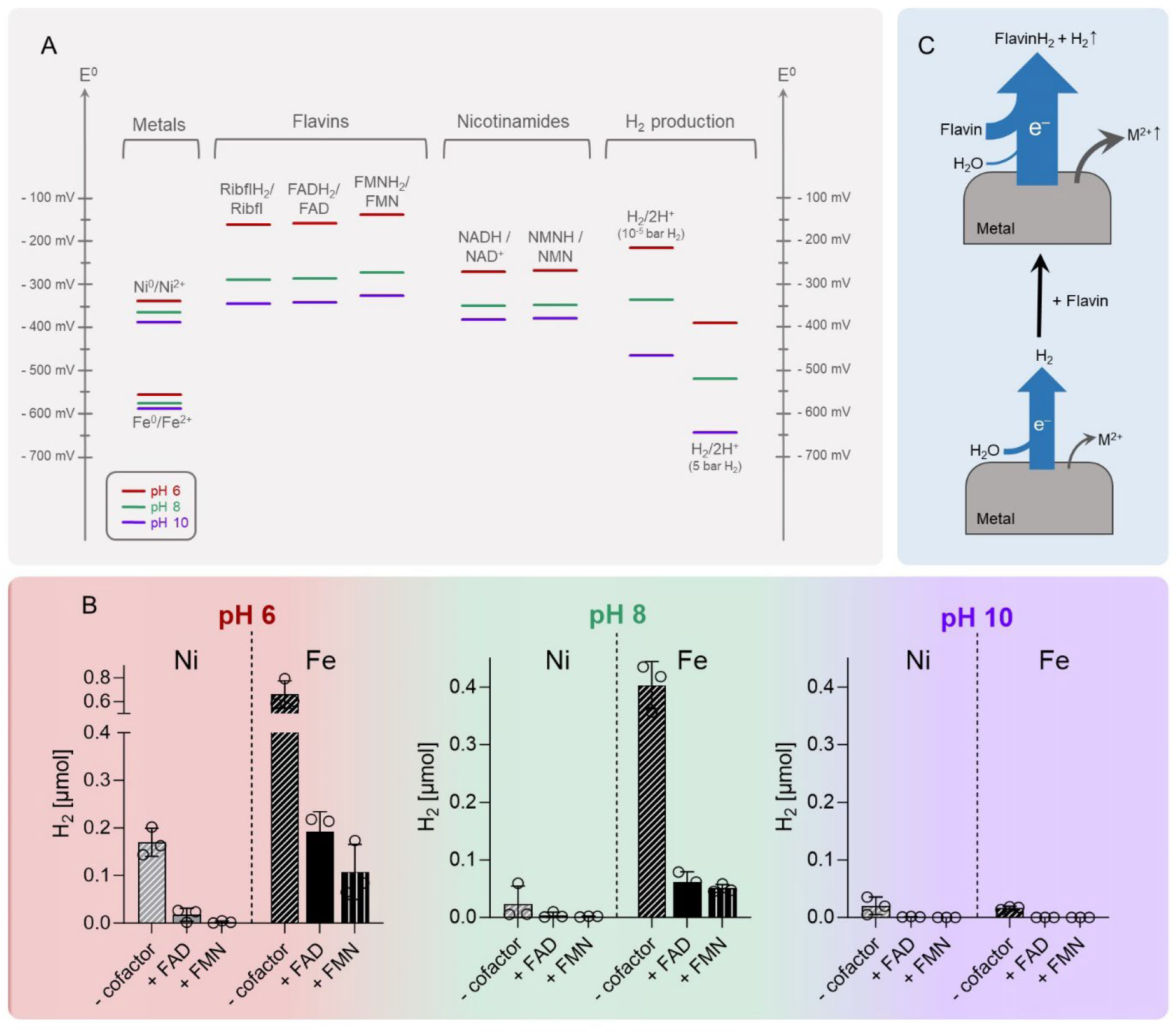
Electron flow within flavin reduction by Ni and Fe. A) Calculated redox potentials of Ni, Fe, flavins, nicotinamides and hydrogen/protons. All calculations based on the Nernst equation are listed in **SI Data 1**. B) Hydrogen production of Ni and Fe under N_2_ atmosphere with and without FAD in solution (***SI Appendix*, Table S23**). This shows how the metals’ electrons rather reduce flavins than protons. Increasing pH decreases both H_2_ production and the yield of the reduced Flavin. C) Proposed electron flow in the metal-flavin-H_2_O systems.

Comparing this artificial setup of electron harvest by flavins, it seems feasible this could happen in any geochemical setting with either native metals or also water-rock interaction systems in which protons are reduced constantly from Fe^2+^. Recently, it was shown that Fe^2+^ indeed can reduce flavins under anaerobic conditions^[61]^, supported also by our observed reduction of FAD with FeS (***SI Appendix*, Figure S5 C**).

### Reduced flavins as electron shuttles and flavin redox cycling

In a prebiotic context, knowing that flavins can scavenge electrons from zero-valent metals opens options for transporting electrons from sources to (micro)environments with completely different conditions, which are more accessible than in H_2_. As mentioned above, there is a variety of microorganisms that shuttle electrons in the form of reduced flavins, more specifically FMN, to reduce Fe^3+^ compounds, making them soluble in the process^[29,62,63]^. As already established, FMN binds metals quicker than FAD^[51]^, echoed by our results displaying significant differences between the metal ion concentrations after FMN and FAD reduction reactions (**Fig. 2 C**). To test if these differences would also influence the reduction of Fe^3+^, we mixed reduced flavins with hematite (Fe_2_O_3_) and magnetite (Fe_3_O_4_) (**Fig. 4 A**). As Fe_3_O_4_ already naturally contains Fe(II) naturally, controls only with oxidized were performed to account for a Fe^2+^ baseline. For the reactions, FMN and FAD are completely reduced with Ni/H_2_ in water, before being mixed with the iron oxides. Fe^3+^ reduction reactions were then performed in unbuffered water to avoid iron phosphate formation, while pHs were monitored to guarantee comparability between samples **(*SI Appendix*, Fig. S11 and Table S29)**. The Fe^3+^ and cofactor concentrations were the same as in the flavin reduction experiments with Ni and Fe, so 4 mM for FMNH_2_ and FADH_2_ and 18 mM for Fe^3+^ per oxide compound. Here, two time points were collected: directly after mixing cofactors with the oxides (t = 0 h) and after a 2 h anaerobic reaction at RT (t = 2 h; Fig. 6). At t = 0 h, we observe a significant difference between FADH_2_ and FMNH_2_ in the Fe_3_O_4_ reaction: FMNH_2_ reduces and dissolves roughly double than FADH_2_. Also, for Fe_2_O_3_ and after 2 h reactions FMNH_2_ always yields more dissolved ions, although the yields get closer to each other. For further experiments we thus decided to focus on FMN and Fe_3_O_4_ as this mineral is also more abundant in serpentinizing systems as it is a major product of the process^[64]^. We also performed additional controls with soluble Fe^3+^ (FeCl_3_) to ensure the robustness of the reduction experiments (cofactor to metal ratio 1:2). Here, all Fe^3+^ was reduced to Fe^2+^ with both FADH_2_ and FMNH_2_ immediately, no difference in yield was detected between them (***SI Appendix*, Fig. S10 and Table S28)**.

**Figure 4:**
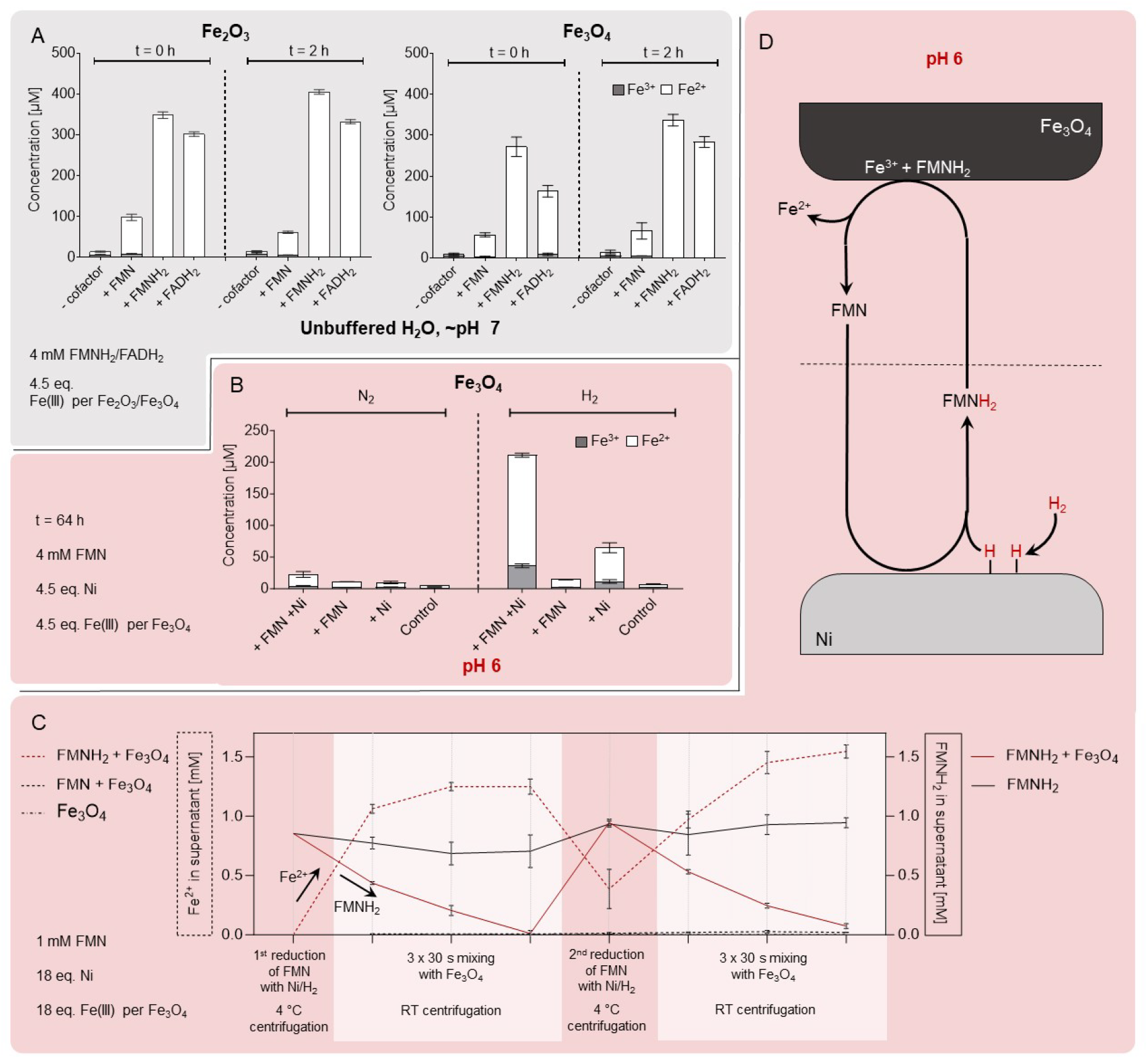
Reduction of Fe^3+^ containing minerals with flavins and redox cyclin of flavins. A) Fe^3+^ reduction with Ni/H_2_ reduced FMNH_2_ (4 mM) from hematite (Fe_2_O_3_) immediately after mixing cofactors and ions and after a 2 h reaction at RT. All reactions in triplicate apart from t = 2 h without flavins (duplicate); Fe^3+^ reduction from magnetite (Fe_3_O_4_) immediately after mixing cofactors and ions together and after a 2 h reaction at RT. All reactions were performed in triplicate. Iron oxides were normalized to 4.5 equivalents (18 mM) of Fe(III). Both Fe^3+^ and Fe^2+^ concentrations were determined via Ferrozine test, the ion concentrations were also determined for the starting solutions of reduced flavin (***SI Appendix*, Table S29**). B) The Fe^2+^ and Fe^3+^ concentration after one pot reactions (64 h) with and without 4 mM of FMN and Fe_3_O_4_ and Ni powder in pH 6 phosphate buffer under 1 bar N_2_ and 5 bar H_2_ atmosphere, respectively. All reactions in triplicate, 4.5 Fe(III) equivalents compared to FMN were added per iron oxide. Both Fe^3+^ and Fe^2+^ concentrations were determined via Ferrozine test (***SI Appendix*, Table S30**). C) Redox cycling of FMN, showing concentrations of FMNH_2_ and Fe^2+^ at all measurement points. FMN was first reduced over Ni with H_2_ gas to FMNH_2_, the supernatant separated from the solid phase to be subsequently mixed for 30 s with Fe_3_O_4_, a process repeated three times so full oxidation back to FMN was achieved. The supernatant was then transferred onto Ni powder again and re-reduced under H_2_ atmosphere before the oxidation process over Fe_3_O_4_ was repeated. All reactions in triplicate, apart from 1^st^ reduction that was performed in bulk. Fe_3_O_4_ were normalized to 4.5 equivalents (18 mM) of Fe(III). Both Fe^3+^ and Fe^2+^ concentrations were determined via Ferrozine test. Controls with FMN and Fe_3_O_4_ were not reduced, Fe_3_O_4_ only was not in the 1^st^ but 2^nd^ reduction reaction, FMNH_2_ control is without Fe_3_O_4_. All controls and exact pHs after reaction are shown in ***SI Appendix* Table S32**. (D) Scheme showing redox cycling of FMN between reduction over Ni/H_2_ and oxidation with Fe^3+^ in Fe_3_O_4_.

As a next step, we combined both Ni and Fe_3_O_4_ powder in pH 6 phosphate buffer under H_2_ and N_2_ atmosphere to investigate if the presence of FMN influences the resulting concentration of Fe^2+^ after 72 h of reaction (**Fig. 4 B**). The addition of acidic buffer was necessary to avoid pH changes triggering ion precipitation during the reaction. Under N_2_, no significant reduction and dissolution of Fe^3+^ (or Fe^2+^) is taking place. Under H_2_, however this changes and the presence of Ni with Fe_3_O_4_ triggers the formation and dissolution of Fe^2+^. This is only outdone by the addition of (oxidized) FMN, although no FMNH_2_ is detected after the reaction indicating that the presence of the flavin facilitates the reduction of Fe(III) in this one-pot setup.

Ultimately, we tested if flavins can be reduced, used as reductant and re-reduced again by separating the reduction (Ni/H_2_) and oxidation (Fe_3_O_4_) reactions (**Figs. 4 C and D**). Also here, we decided to conduct these experiments in pH 6 phosphate buffer to minimize ion precipitation due to pH fluctuations. After the 1^st^ cycle was complete and resulting was FMN re-reduced with Ni, samples were cold centrifuged (4° C) which led to a precipitation of iron ions. During the three subsequent oxidation steps, FMNH_2_ is fully oxidized back to FMN for a second time. Our results clearly show how flavins act as transferring agents between reducing and oxidizing environments, staying very stable at least for the first two cycles shown here. Flavins would additionally serve as transition metal ligands, facilitating the diffusion of otherwise badly soluble ions.

Overall, we see that we can i) couple geochemical flavin reduction with subsequent Fe^3+^ reduction and ii) see a difference between FAD and FMN reactions. This shows how flavins can be used as electron shuttles also in a prebiotic context and that different properties of the mono- and dinucleotide can lead to variances in reaction yields.

## Conclusion

Our results demonstrate that flavins can scavenge electrons that would otherwise be released as H_2_. They effectively function as low-energy electron acceptors capable of harvesting reducing equivalents from metal surfaces under conditions where hydride carriers such as NAD could not. We furthermore show how flavins can be recycled as electron carriers in mineral-rich environments. These conditions were designed roughly after serpentinizing systems in the oceanic crust, but can be transferred to many different reducing environments. For both ocean- and land-based H_2_-generating environments, one could argue that the synthesis of H_2_ represents a loss of energy as the generated gas dissipates into the atmosphere. In that case, flavins could keep these electrons in a chemically accessible form. Moreover, fully reduced flavins (hydroquinone) could access their one-electron chemistry by forming semiquinone radicals, leading to far more negative redox potentials^[65]^. It has been shown that metal ions can stabilize flavin radicals, offering a potential mechanism for changing electron energy without the complex enzymatic mechanism of flavin-based electron bifurcation^[66]^. Our observation of red pellet formation suggests that semiquinones are precipitating during the reactions, possibly opening an abiotic path to downstream radical chemistry^[26,67]^.

The structural differences between riboflavin, FMN, and FAD seem to be relevant beyond simple solubility considerations. We show that FAD yields different results than FMN when it comes to complexation of metal ions, both during the reduction of flavins via Ni and Fe and the reduction of metal oxides by reduced flavins. FAD undergoes intramolecular interactions of its adenine and isoalloxazine moieties in aqueous solution, likely restricting its interactions with metal ions^[33,52,68]^. A similar intramolecular folding in NAD has been proposed to prevent overreduction of the nicotinamide ring, preserving its function as a strictly two-electron hydride carrier^[43]^. Whether the conformation of FAD could also serve a protective function or whether it represents a structural constraint, remains an open question. FMN, unlike its mononucleotide analogue NMN, is not merely a biosynthetic intermediate *en route* to FAD, but a cofactor with its own biological functions. The differences in metal complexation, electron transfer, and aggregation behaviour we observe between FMN and FAD (and also partly in riboflavin) suggest that precise flavin structure matters and that these differences may have been consequential in selecting which flavin species predominated under specific prebiotic geochemical conditions. That extant biology likely only uses flavins as shuttles in the latter direction, could still point toward a primordial electron shuttling role.

In this paper, we have only tested a couple of flavin’s many biological functions that could have been relevant to the origin of life – electron shuttles and recyclable redox entities in an emerging protometabolic context. But looking into the role of flavins in primary carbon metabolism in LUCA, electron transfer, seems a logical beginning. Flavins are among the most ancient cofactors and are central to autotrophic carbon fixation as found in LUCA^[38]^. The chemical stability of the isoalloxazine ring system across different pH values and redox conditions (+/-H_2_) also shows that flavins could have been functional before the emergence of stable enzymatic scaffolds. In this context, flavins may have served in protometabolic networks, enabling electron and metal ion transfer between geochemically distinct microenvironments, keeping a variety of redox reactions simultaneously viable.

## Methods

### Flavin reduction experiments

All sample preparation was conducted in a glove box (Jacomex) under a nitrogen atmosphere and red light conditions to prevent oxidation or photo-induced degradation. Stock solutions of flavins were prepared at a concentration of 4 mM using either Flavin Adenine Dinucleotide Disodium Salt (FAD; 94%, Tokyo Chemical Industry) or Riboflavin 5′-Monophosphate Sodium Salt (FMN; >85%, TCI). These were dissolved in either 0.133 M phosphate buffer (pH 6 or 8) or 0.1 M carbonate buffer (pH 10). Based on preliminary experiments, no significant differences in reduction behaviour were observed between carbonate and phosphate buffer systems at pH 10 (***SI Appendix*, Fig. S24**). Metal powders (18 mM; overview of particle sizes and suppliers in ***SI Appendix*, Table S1**) were weighed into individual 1 mL glass vials (Ni^0^: 1 mg/1.1 μmol; Fe^0^: 1 mg/1.0 μmol). Subsequently, 1 mL aliquots of the flavin stock solution were added to the vials containing the metal powders, along with a magnetic stirring bar. The vials were sealed with pierced crimp caps to allow gas exchange between the vial interior and the reactor environment (***SI Appendix*, Fig. S1**). The prepared samples were placed into a high-pressure 300 mL steel reactor (Berghof). The reactor was sealed, removed from the glove box, and either pressurised with 5 bar H_2_ (99.9% Nippon Gases) or maintained under an inert atmosphere, depending on the experimental conditions. The reaction was carried out at 40 °C for 15 min or 2 h, with continuous stirring at 400 rpm. Directly following the reaction, the reactor was depressurised and flushed three times with argon (Ar) gas. The system was cooled on ice until it returned to ambient temperature to slow down further reduction. Subsequently, the reactor was transferred back into the glove box for processing, such as immediate pH determination of individual samples to guarantee pH stability (***SI Appendix*, Table S21)**. Riboflavin (Avantor/VWR) was subjected to the same experimental procedure as described for FAD and FMN. However, due to its low solubility, the initial concentration were adjusted to 0.1 mM.

### UV-Vis measurements and quantification

Following reactor retrieval, samples were transferred back into the glovebox and centrifuged at 4°C. The supernatant was collected and diluted appropriately for subsequent analyses as described below. For product quantification, the characteristic absorption spectra of flavins were used. Samples were diluted to a final concentration of 0.1 mM for UV-Vis detection (apart from Riboflavin, the concentration of which was already at 0.1 mM). Aliquots were transferred into airtight quartz cuvettes, sealed, and measured over a wavelength range of 320–460 nm using a Cary 60 UV-Vis spectrophotometer (Agilent Technologies) operated with Cary WinUV Scanning Kinetics software (version 5.1.3.1042). An initial spectrum was recorded immediately upon cuvette placement. To assess the fully re-oxidised state, cuvettes were subsequently opened, gently aerated, and mixed by shaking twice before recording a final spectrum. Reduced flavins were quantified based on their characteristic absorbance maximum and minimum, referenced against pH- and concentration-matched calibration curves prepared from the same cofactor (***SI Appendix*, SI Tables S2–20**). The exact sample concentration was first determined from the absorbance maximum of the reoxidised sample. The absorbance minimum was then compared to the theoretical limit to derive a reduction ratio (***SI Appendix*, SI Eqs. S1–5**).

### Inductively Coupled Plasma Mass Spectrometry (ICP-MS)

ICP-MS was used to quantify dissolved metal ions following a reaction (***SI Appendix*, Table S24**). Samples were prepared post-reaction by mixing 100 μL of the centrifuged (4 °C, 30 min, 13 000 rpm) sample with 100 μL of concentrated nitric acid (HNO_3_, 67%) into a 15 mL Falcon tube. These were then heated at 60°C for 3 h in a hot-water bath. Afterwards, the samples were topped up to a volume of 10 mL with HPLC water and vortexed. The Fe and Ni content of the diluted samples was determined using an Agilent 7850 quadrupole ICP-MS (Agilent Technologies, Santa Clara, USA) operated in helium collision mode with kinetic energy discrimination (He-KED). The system was run in low-matrix plasma mode, with an RF power of 1550 W, a sampling depth of 8 mm, and a carrier gas (argon) flow rate of 0.99 L/min. Helium was used as the cell gas at a flow rate of 4.6 mL/min, and the energy discrimination voltage was set to 3 V. Acquisition was performed in He-KED mode, using three replicates per measurement and a dwell time of 0.3 seconds per mass. Samples were introduced via a PFA MicroMist concentric nebulizer into a temperature-controlled quartz cyclonic spray chamber maintained at 2 °C. A standard quartz torch with a 2.5 mm i.d. injector was used, and the interface included Ni sampling and skimmer cones. Calibration was performed using external standards prepared from single-element Fe and Ni salt stock solution (LGC Standards), acidified with 1% (v/v) HNO_3_. The calibration range covered 1–100 ng/mL. A germanium internal standard at 300 ng/mL (LGC Standards) was for internal normalization. Instrument control and data analysis were carried out using MassHunter 5.3 software.

### Gas Chromatography Thermal Conductivity Detection (GC-TCD) of H_2_ production during reaction

GC-TCD was used to monitor the H_2_ generation of the flavin reduction experiments under N_2_ atmosphere. The measurements were performed on a SRI 8610C gas chromatograph (SRI Instruments Europe) with two back-to-back 6-foot Haysep D columns. The GC oven was set to 42 °C, the detector to 308 °C, and the N_2_ carrier gas to 24 °C flow rate was set to 1 mL/min before operating. The program used was PeakSimple Version 4.54. The gas was collected from the headspace of the sample vials with a gas-tight syringe (Hamilton™ 1000er-Serie Gastigh™ Spritzen), which was then inserted into the GC. To secure a gas-tight environment, the vial caps were taped with electrical tape prior to the reaction. The H_2_ concentration of the injected sample was quantified using calibration curves (***SI Appendix*, Fig. S8** and **Table *S*22**). The calibration samples were made by flushing a membrane-capped, and electrical tape-insulated, gas-tight 50 mL bottle with 1 bar of H_2_. The bottle was turned upside down so that its capped top was pointing downwards, secured to a retort stand in the fumehood. The vial’s cap was pierced by in- and outlet syringe needles. The inlet was connected to a hydrogen supply that was set to 1 bar. At the end of the flushing, the outlet syringe was removed first, and the pressure was monitored to be 1 bar from a gauge that was set on the inlet. The inlet was removed afterwards, and the bottle immediately taken to the GC-TCD. The gas bottle was sampled three times and measured consecutively before refilling and flushing it for the next measurement. To set a calibration curve, the extracted hydrogen volume from the stock bottle was altered between 200, 100, and 50 μL, and this was then diluted with atmospheric gas to a total volume of 600 μL. The gases were allowed to mix briefly before inserting a volume of 300, 200, 100, or 50 μL of the mixture gas into the GC-TCD (***SI Appendix*, Table S22**). The molarity of each injection was calculated using the ideal gas law (***SI Appendix*, Eqs. S6 and S7, and Table S23**). The dissolved hydrogen of each sample was subsequently determined following Henry’s law (**Eq. S7**).

### Iron oxide reduction with flavins and Ferrozine assay

FAD and FMN (4 mM) in water (HPLC grade) were reduced under H_2_ with Ni^0^ for 2 h. The samples were centrifuged (4 °C, 13,000 rpm, 20 min), and the supernatant recovered from reaction tube. The pH of all samples was measured immediately and complete reduction was confirmed via UV-Vis (***SI Appendix*, Table S29**). All preparation was done under gloveboxes N_2_ atmosphere before the measurement. The amount of Fe_2_O_3_ and Fe_3_O_4_ (Sigma-Aldrich) was normalized per moles of Fe(III) in each iron oxide (18 mM or 4.5 equivalents relative to the cofactor). 1 mL of reduced flavin solution and stirbar were added to the iron oxides. A 50 μL aliquot was taken from samples 1–24 at t = 0 for the Ferrozine assay. All samples were stirred at room temperature, 400 rpm, for 2 h, under N_2_ atmosphere. All samples were then measured with a Ferrozine assay (***SI Appendix*, Table S29**). Ferrozine stock solution (10 mM) was prepared in 0.1 M ammonium acetate and then diluted to 1 mM with HPLC water before use. Hydroxylamine hydrochloride stock (1.4 M in 2 M HCl) was diluted to 4 mM with HPLC water. A 50 μL aliquot of the sample was added with 50 μL Ferrozine solution (1 mM) and 200 μL of HPLC water. The mixture was vortexed, centrifuged at RT, 13,000 rpm, for 10 min. From this, 100 μL were pipetted into a 96-well plate well. The samples were measured with a plate reader, and the absorbance was recorded at 562 nm. After the measurement, 20 μL of hydroxylamine hydrochloride (4 mM) was added to each sample, gently mixed, and left to incubate in the dark for 10 min, before 20 μL was removed from each sample to keep the volume constant (100 μL) between the measurements. The samples were measured again, and the absorbance values were converted to concentrations according to the standard curve and corrected for dilutions (**Figure 4; *SI Appendix*, Tables S25 and S26**).

### FMNH_2_ redox-cycling with Fe_3_O_4_

25 mL of a 1 mM FMN stock solution was reduced with 5 bar H_2_ over 18 mM of Ni^0^ powder at 40 °C for 2 h. After reaction, the solution was centrifuged at 4 °C, 30 min, 13,000 rpm and was anaerobically stored for further use. The following reactions were performed under anoxic conditions and red light to avoid uncontrolled oxidation or light degradation. The amount of used Fe_3_O_4_ was normalized per moles of Fe(III) in each iron oxide (18 mM or 18 equivalents relative to the cofactor). FMNH_2_ (1 mM, 1.7 mL) was then added to Fe_3_O_4_. The mixture was vortexed for 10 seconds, and then centrifuged at 20 °C for 20 min. 200 μL aliquots of the supernatants were taken for further analysis (pH, Ferrozine, UV-Vis), and the remaining supernatant was transferred to a fresh batch of Fe_3_O_4_, adjusted for volume change (***SI Appendix*, Tables S26 and S32**). This process was repeated three times, after which most of the FMNH_2_ was oxidized to FMN. The remaining supernatant, still containing FMN, was put through another reduction reaction. The reduction of supernatant was done using the same procedure (Ni/H_2_) as before, unless in the control with oxidized FMN instead of FMNH_2_. The full reduction was confirmed using UV-Vis spectroscopy. Then, the reaction was repeated analogously to the first three cycles. The concentration of Fe_3_O_4_ was adjusted to the total volume of the reaction for each cycle to keep the same ratio between cofactor and Fe(III).

## Author contributions

**O.J.L**.: Conceptualization, Methodology, Data Curation, Investigation, Visualization, Writing – Review & Editing **D.P.H.P**.: Conceptualization, Methodology, Investigation, Data Curation, Visualization, Writing – Review & Editing, Supervision. **M.T.Y**.: Methodology, Investigation, Data Curation. **N.P**.: Methodology, Data Curation, Writing – Review & Editing. **M.P**.: Conceptualization, Data Curation, Formal Analysis, Methodology, Visualization, Writing – Original Draft, Supervision, Funding Acquisition.

## Acknowledgements

MP thanks the European Research Council (101221521), the Human Frontiers Science Program (RGEC29/2025), the International Max Planck Research School “Principles of Microbial Life” and the Max Planck Society for funding.

The authors thank Alicia Casitas for assisting with cyclic voltammetric (CV) measurements, Uwe Linne and Heike Mallinger for μXRF measurements, and Mahiob Dawor, Manavika Khanna and Hikmet Baki for exploratory flavin reduction experiments. We further thank Marco W. Fraaije for critical reading of the first draft of this manuscript and Bill Martin, Crispin Lichtenberg, Assma Choudna and Michael von Domaros for discussions.

