## Supplementary material for "Flavin cycling under prebiotic conditions: bidirectional electron transfer and versatility in nickel and iron containing environments": SI_Appendix

**This PDF file includes:**

Supporting text

Figures S1 to S24

Tables S1 to S34

**Other supporting materials for this manuscript include the following:**

SI Data 1, excel file with Redox potential calculations.

**Methods**

**Experimental set up**

**Table S1.** Each mineral/metal powder was standardized to have the same amount of metal atoms (18 μmol).

| Minerals | Molar mass (g/mol) | 18 μmol (mg) | Particle size (nm) | Purity (%) | Vendor |
|---|---|---|---|---|---|
| $Ni^0$ | 58.69 | 1.1 | < 100 | >99 | Sigma-Aldrich |
| $Fe^0$ | 55.845 | 1 | 60-70 | >99.55 | Nanografi |
| NiO | 74.6928 | 1.8 | 14-29 | 76 | Thermo Scientific |
| $Fe_3O_4$ | 231.533 | 1.4 | < 50 | 97 | Sigma-Aldrich |
| FeS | 87.91 | 1.6 | ≈150000 | 99.9 | Thermo Scientific |

**High Pressure Reactions.** For the experiments a 300 mL high-pressure reactor (BR-300, Berghof) was used. The reactor was set to reactor heating system (Berghof Products + Instruments) at 400 rpm and 40 °C. The reaction time was set to 15 minutes or 2 hours once the internal temperature reached 40°C. The atmosphere of the reactor was either filled to 5 bars of $H_2$ (99.9% Nippon Gases), or maintained with the glove-box atmosphere ($N_2$ 99.999%, university faculty). The sample's pH was measured before and after the reaction with an electronic pH meter (FiveEasyPlus pH/mV, Mettler Toledo). Once the reaction was complete, the reactor was depressurised, immediately put in ice water, flushed three times with Ar, and taken into the glovebox for further analysis once the reactor had cooled down back to room temperature. The work was conducted under red light to avoid photodegradation.

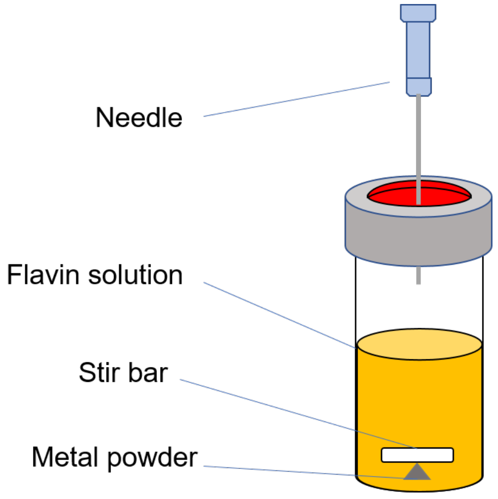


**Fig. S1.** The samples were prepared in 2 mL glass vials. After the solution, a PTFE-coated stirring bar, and metal powder were added. The sample was closed with a membrane lid and pierced with a syringe needle to allow gas exchange.

**Ultraviolet-Visible (UV–Vis) Spectroscopy.** Considering the UV–Vis absorbance spectra of flavins (see **Fig. S2.** ) their spectra were scanned between the wavelengths of 320-460 nm on a Cary 60 UV–Vis with the Cary WinUV, Scanning Kinetics Application, version 5.1.3.1042 program. A baseline of buffer was taken as a reference for carbonate buffer samples, while water was used for phosphate buffer samples. The spectra are measured at room temperature in air-tight UV–quartz cuvettes. For that, each FAD or FMN sample was diluted in buffer to 0.1 mM, after centrifuging (Fresco 17 centrifuge, Thermo Scientific) for 30 min, at 4°C, and 13 000 rpm, to obtain relevant UV–Vis absorption spectra. Riboflavin samples were already 0.1 mM, and were pipetted directly into the cuvettes after the same centrifugation. Sequentially, each sample was opened, gently blown into, and shaken with a cap on, twice over, before measuring it again to collect the spectrum of the fully oxidized sample.

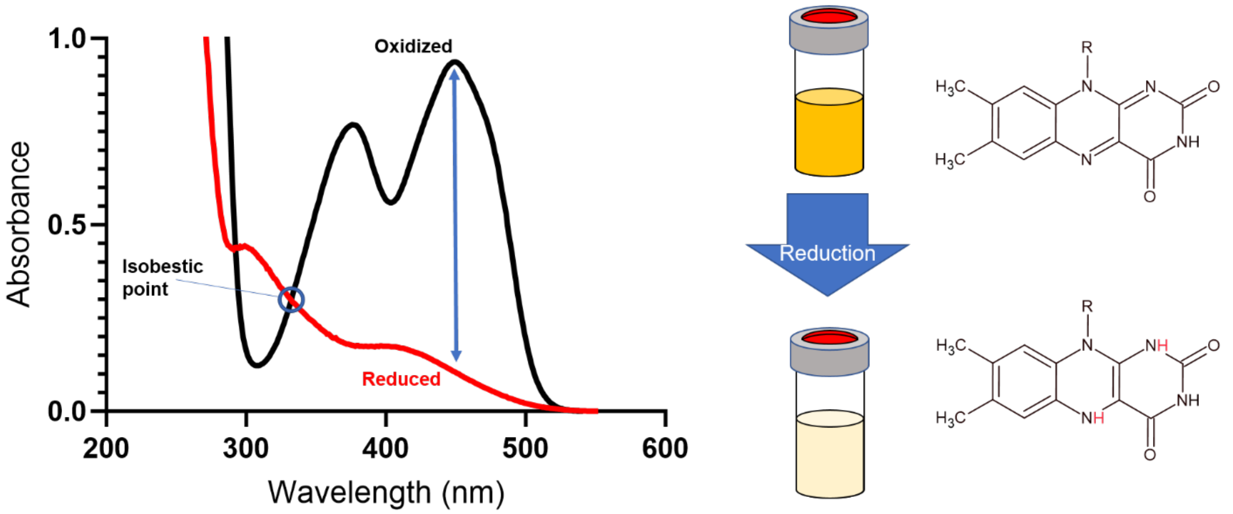


**Fig. S2.** Reduction of the flavin's isoalloxazine ring changes the molecule's ability to absorb light in the 334-500 nm region, which decreases the solution's yellow colour intensity. This can be monitored with UV–Vis spectroscopy.

**Quantification.** To quantify the reduced flavin in solution after each reaction, the flavins' characteristic light absorbance was referenced and monitored through UV-Vis spectroscopy. Reference spectra of fully reduced and oxidised flavins were prepared as standards for a range of concentrations and pHs (see **Table S3─20.**). The standards were made by preparing 10 mM stock solutions of flavins, and from which sequential dilutions were made. For reduction, 100 mM stock solution of sodium hydrosulfite (85% Thermo Scientific) was prepared in water. Aliquots corresponding to a final threefold molar excess relative to the standards were added to obtain reduction. According to the Lambert-Beer Law, the equations obtained from the standards, should describe a linear relationship between absorbance ($\boldsymbol{Abs}$) and concentration ($\boldsymbol{c}$), where the light path ($\boldsymbol{l}$) is always 1, and thus the slope ($\boldsymbol{m}$) is the inverse of the extinction coefficient ($\boldsymbol{\varepsilon}$) of the molecule at a specific wavelength (see **Eq. S1.**). Each calibration curve was done with stock solutions of at least 5 different concentrations, which were measured thrice. Conventionally, isobestic points are used as reference points for spectrophotometric quantification. Due to our samples containing other species that absorb at the same wavelength, such as iron phosphates, samples were quantified based on the flavins' highest absorbance peak after reoxidation (see **Eq. S3.**)**,** where less overlapping absorptions from other species are expected.

**Eq. S1.** Lambert-Beer Law

$$Abs = \varepsilon * c * l$$

**Eq. S2.** Standard curve equation

$$c = \frac{1}{\varepsilon} * Abs$$

Abs = Absorbance value
c = concentration (mM)
l = path lenght (1)
ε = extinction coefficient

**Table S2.** The UV-Vis spectroscopy calibration curves' equations in pHs 6, 8, and 10, with their respective $r^2$ values in brackets, are specified for each cofactor. The x-axis is absorption, and the y-axis is concentration of cofactor in μM. Riboflavin's maximum absorbance peak is at 446 nm, FMN's is 448 nm, and FAD's is 450 nm. The equations on the left relate to the fully oxidised standard solutions, while the equations on the right relate to the fully reduced standard solutions.

| Cofactor | pH | $c_s = m_{max} * Abs_{max}$ | $c_s = m_{min} * Abs_{min}$ |
|---|---|---|---|
| **FAD** | 6 | y = 89.20976x (0.9883) | y = 774.2874x (0.9971) |
| | 8 | y = 90.6254x (0.9958) | y = 1379.544x (0.9893) |
| | 10 | y = 108.006x (0.9874) | y = 1335.27x (0.991) |
| **FMN** | 6 | y = 86.122x (0.9924) | y = 663.0255x (0.9959) |
| | 8 | y = 64.7785x (0.9983) | y = 1338.495x (0.9959) |
| | 10 | y = 76.23395x (0.9966) | y = 1642.285x (0.9969) |
| **Riboflavin** | 6 | y = 78.104x (0.9921) | y = 729.43x (0.9829) |
| | 8 | y = 89.018x (0.997) | y = 1383.6x (0.9896) |
| | 10 | y = 93.654x (0.996) | y = 1511.2x (0.9924) |

**Table S3.** Calibration curve of maximum absorbance value of FAD pH 10. using a spectrophotometer at λ = 448 nm. To each concentration point (y), three replicas were measured to obtain the corresponding absorbance values (x).

<table>
<tr><th colspan="3">FAD pH10</th></tr>
<tr><td>Concentration vs $\mathbf{Abs_{max}}$ (450nm)</td><td>Concentration (µM)</td><td>Absorbance</td></tr>
<tr><td rowspan="15">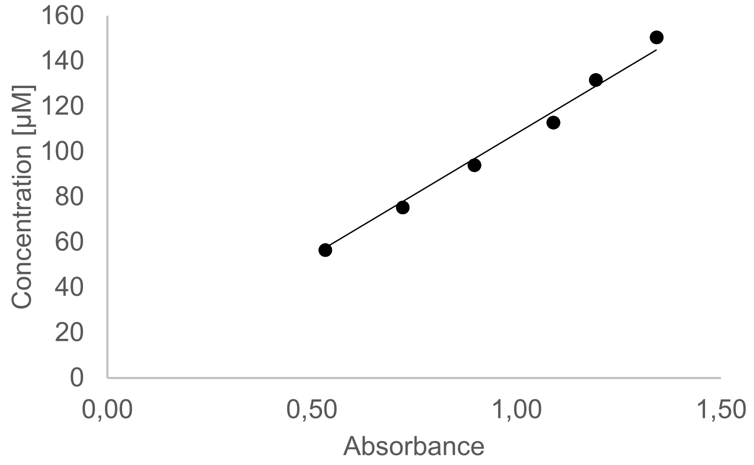</td><td rowspan="3">150.4</td><td>1.342</td></tr>
<tr><td>1.342</td></tr>
<tr><td>1.342</td></tr>
<tr><td rowspan="3">131.6</td><td>1.194</td></tr>
<tr><td>1.194</td></tr>
<tr><td>1.194</td></tr>
<tr><td rowspan="3">112.8</td><td>1.090</td></tr>
<tr><td>1.090</td></tr>
<tr><td>1.090</td></tr>
<tr><td rowspan="3">94</td><td>0.897</td></tr>
<tr><td>0.897</td></tr>
<tr><td>0.897</td></tr>
<tr><td rowspan="3">75.2</td><td>0.722</td></tr>
<tr><td>0.722</td></tr>
<tr><td>0.722</td></tr>
<tr><td rowspan="3">$\mathbf{y = 108.006x;\ R^2 = 0.9874}$</td><td rowspan="3">56.4</td><td>0.533</td></tr>
<tr><td>0.532</td></tr>
<tr><td>0.532</td></tr>
</table>

**Table S4.** Calibration curve of maximum absorbance value of FMN pH 10 using a spectrophotometer at λ = 448 nm. To each concentration point (y), three replicas were measured to obtain the corresponding absorbance values (x).

<table>
<tr><th colspan="3">FMN pH10</th></tr>
<tr><td>Concentration vs $\mathbf{Abs_{max}}$ (448nm)</td><td>Concentration (µM)</td><td>Absorbance</td></tr>
<tr><td rowspan="15">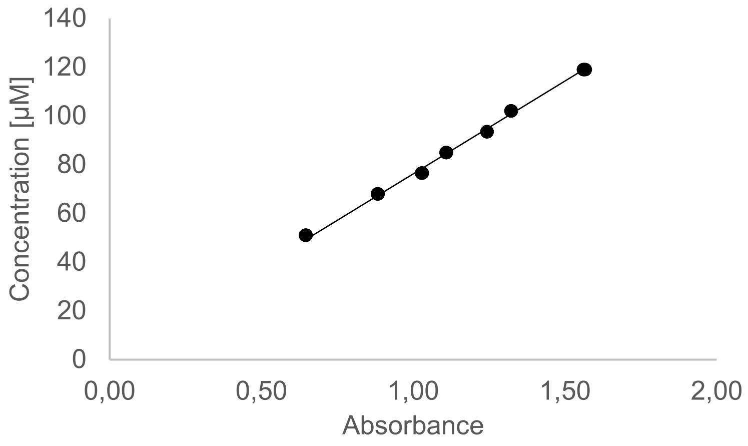</td><td rowspan="3">119</td><td>1.561</td></tr>
<tr><td>1.563</td></tr>
<tr><td>1.567</td></tr>
<tr><td rowspan="3">102</td><td>1.322</td></tr>
<tr><td>1.323</td></tr>
<tr><td>1.322</td></tr>
<tr><td rowspan="3">93.5</td><td>1.243</td></tr>
<tr><td>1.243</td></tr>
<tr><td>1.243</td></tr>
<tr><td rowspan="3">85</td><td>1.110</td></tr>
<tr><td>1.109</td></tr>
<tr><td>1.108</td></tr>
<tr><td rowspan="3">76.5</td><td>1.028</td></tr>
<tr><td>1.028</td></tr>
<tr><td>1.030</td></tr>
<tr><td rowspan="3">$\mathbf{y = 76.234x;\ R^2 = 0.9966}$</td><td rowspan="3">68</td><td>0.885</td></tr>
<tr><td>0.884</td></tr>
<tr><td>0.882</td></tr>
</table>

**Table S5.** Calibration curve of minimum absorbance value of FAD pH 10 using a spectrophotometer at λ = 450 nm. To each concentration point (y), three replicas were measured to obtain the corresponding absorbance values (x).

<table>
<tr><th colspan="3">FAD pH10</th></tr>
<tr><td>Concentration vs $\mathbf{Abs_{min}}$ (450nm)</td><td>Concentration (µM)</td><td>Absorbance</td></tr>
<tr><td rowspan="15">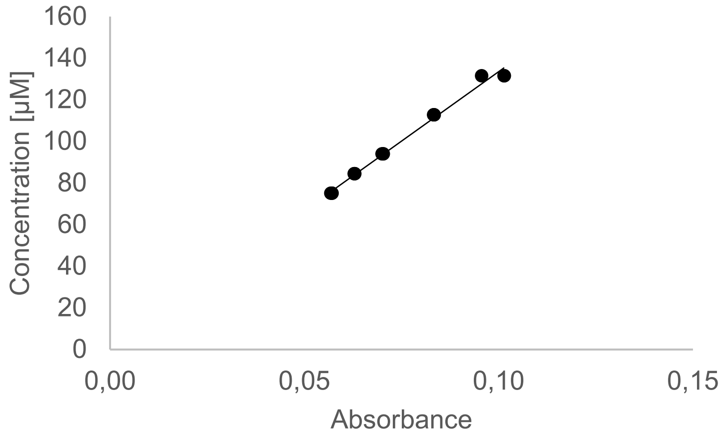
</td><td rowspan="3">131.6</td><td>0.096</td></tr>
<tr><td>0.101</td></tr>
<tr><td>0.101</td></tr>
<tr><td rowspan="3">112.8</td><td>0.083</td></tr>
<tr><td>0.083</td></tr>
<tr><td>0.083</td></tr>
<tr><td rowspan="3">94</td><td>0.070</td></tr>
<tr><td>0.070</td></tr>
<tr><td>0.070</td></tr>
<tr><td rowspan="3">84.6</td><td>0.063</td></tr>
<tr><td>0.063</td></tr>
<tr><td>0.063</td></tr>
<tr><td rowspan="3">75.2</td><td>0.057</td></tr>
<tr><td>0.057</td></tr>
<tr><td>0.057</td></tr>
<tr><td rowspan="3">$\mathbf{y = 1335.3x;\ R^2 = 0.991}$</td><td rowspan="3">131.6</td><td>0.096</td></tr>
<tr><td>0.101</td></tr>
<tr><td>0.101</td></tr>
</table>

**Table S6.** Calibration curve of minimum absorbance value of FMN pH 10 using a spectrophotometer at λ = 448 nm. To each concentration point (y), three replicas were measured to obtain the corresponding absorbance values (x).

<table>
<tr><th colspan="3">FMN pH10</th></tr>
<tr><td>Concentration vs $\mathbf{Abs_{min}}$ (448nm)</td><td>Concentration (µM)</td><td>Absorbance</td></tr>
<tr><td rowspan="15">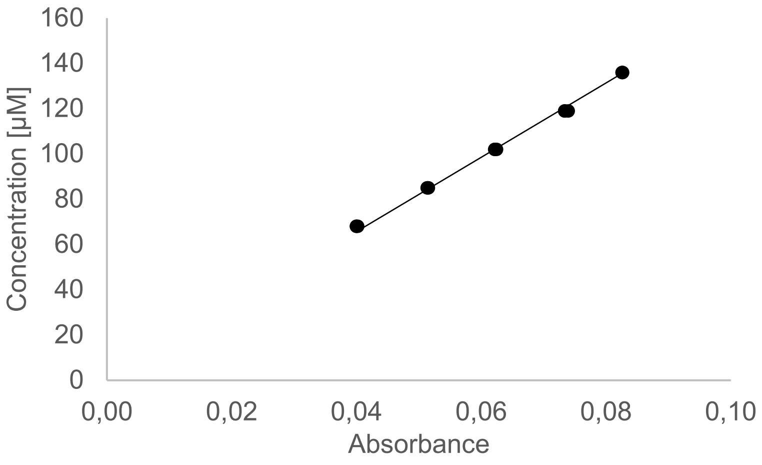
</td><td rowspan="3">136</td><td>0.083</td></tr>
<tr><td>0.083</td></tr>
<tr><td>0.083</td></tr>
<tr><td rowspan="3">119</td><td>0.074</td></tr>
<tr><td>0.073</td></tr>
<tr><td>0.073</td></tr>
<tr><td rowspan="3">102</td><td>0.062</td></tr>
<tr><td>0.062</td></tr>
<tr><td>0.062</td></tr>
<tr><td rowspan="3">85</td><td>0.051</td></tr>
<tr><td>0.052</td></tr>
<tr><td>0.051</td></tr>
<tr><td rowspan="3">68</td><td>0.040</td></tr>
<tr><td>0.040</td></tr>
<tr><td>0.040</td></tr>
<tr><td rowspan="3">$\mathbf{y = 1642.3x;\ R^2 = 0.9969}$</td><td rowspan="3">136</td><td>0.083</td></tr>
<tr><td>0.083</td></tr>
<tr><td>0.083</td></tr>
</table>

**Table S7.** Calibration curve of maximum absorbance value of FAD pH 8 using a spectrophotometer at λ = 450 nm. To each concentration point (y), three replicas were measured to obtain the corresponding absorbance values (x).

| FAD pH8 | | |
|---|---|---|
| *Concentration vs* $\mathbf{Abs_{max}}$ *(450nm)* | *Concentration (µM)* | *Absorbance* |
| 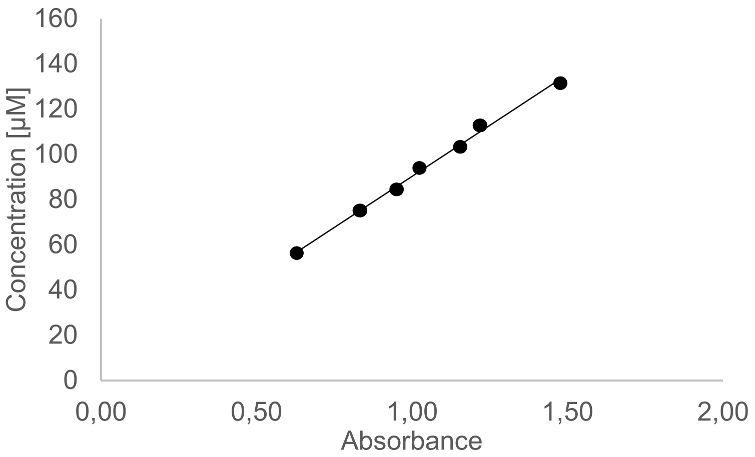  | **131.6** | **1.478** |
| | | **1.476** |
| | | **1.476** |
| | **112.8** | **1.216** |
| | | **1.220** |
| | | **1.221** |
| | **94** | **1.153** |
| | | **1.155** |
| | | **1.156** |
| | **84.6** | **1.023** |
| | | **1.024** |
| | | **1.025** |
| | **75.2** | **0.949** |
| | | **0.950** |
| | | **0.951** |
| $\mathbf{y = 90.295x;\ R^2 = 0.9958}$ | **131.6** | **0.831** |
| | | **0.832** |
| | | **0.834** |

**Table S8.** Calibration curve of maximum absorbance value of FMN pH 8 using a spectrophotometer at λ = 448 nm. To each concentration point (y), three replicas were measured to obtain the corresponding absorbance values (x).

| FMN pH8 | | |
|---|---|---|
| *Concentration vs* $\mathbf{Abs_{max}}$ *(448nm)* | *Concentration (µM)* | *Absorbance* |
| 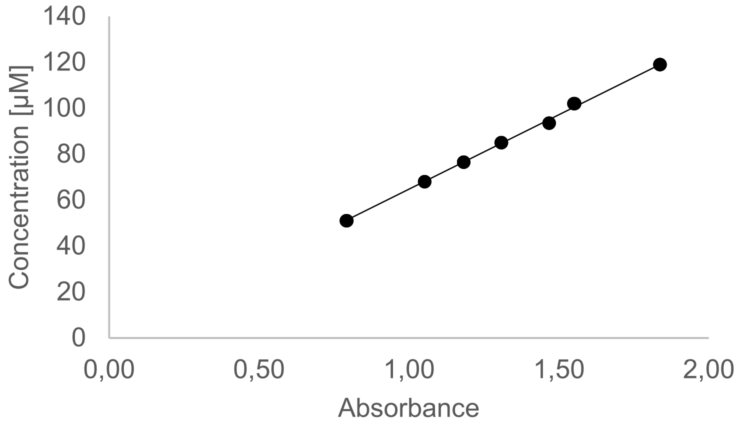  | **119** | **1.838** |
| | | **1.836** |
| | | **1.837** |
| | **102** | **1.552** |
| | | **1.550** |
| | | **1.550** |
| | **93.5** | **1.468** |
| | | **1.468** |
| | | **1.468** |
| | **85** | **1.309** |
| | | **1.308** |
| | | **1.308** |
| | **76.5** | **1.182** |
| | | **1.182** |
| | | **1.183** |
| $\mathbf{y = 64.778x;\ R^2 = 0.9983}$ | **68** | **1.052** |
| | | **1.052** |
| | | **1.052** |

**Table S9.** Calibration curve of minimum absorbance value of FAD pH 8 using a spectrophotometer at λ = 450 nm. To each concentration point (y), three replicas were measured to obtain the corresponding absorbance values (x).

<table>
<tr><th colspan="3">FAD pH8</th></tr>
<tr><td>Concentration vs $\mathbf{Abs_{min}}$ (450nm)</td><td>Concentration (μM)</td><td>Absorbance</td></tr>
<tr><td rowspan="15">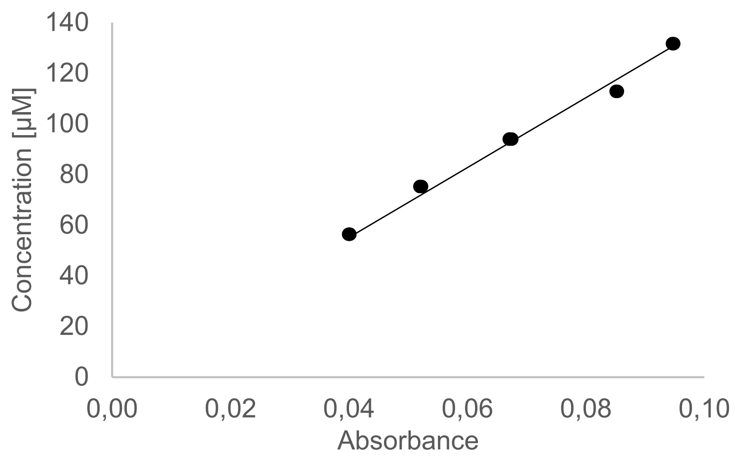
</td><td rowspan="3">131.6</td><td>0.095</td></tr>
<tr><td>0.095</td></tr>
<tr><td>0.095</td></tr>
<tr><td rowspan="3">112.8</td><td>0.085</td></tr>
<tr><td>0.085</td></tr>
<tr><td>0.085</td></tr>
<tr><td rowspan="3">94</td><td>0.067</td></tr>
<tr><td>0.067</td></tr>
<tr><td>0.067</td></tr>
<tr><td rowspan="3">84.6</td><td>0.052</td></tr>
<tr><td>0.052</td></tr>
<tr><td>0.052</td></tr>
<tr><td rowspan="3">75.2</td><td>0.040</td></tr>
<tr><td>0.040</td></tr>
<tr><td>0.040</td></tr>
<tr><td rowspan="3">$\mathbf{y = 1379.5x;\ R^2 = 0.9893}$</td><td rowspan="3">131.6</td><td>0.095</td></tr>
<tr><td>0.095</td></tr>
<tr><td>0.095</td></tr>
</table>

**Table S10**. Calibration curve of minimum absorbance of FMN pH 8 using a spectrophotometer at λ = 448 nm. To each concentration point (y), three replicas were measured to obtain the corresponding absorbance values (x).

<table>
<tr><th colspan="3">FMN pH8</th></tr>
<tr><td>Concentration vs $\mathbf{Abs_{min}}$ (448nm)</td><td>Concentration (μM)</td><td>Absorbance</td></tr>
<tr><td rowspan="15">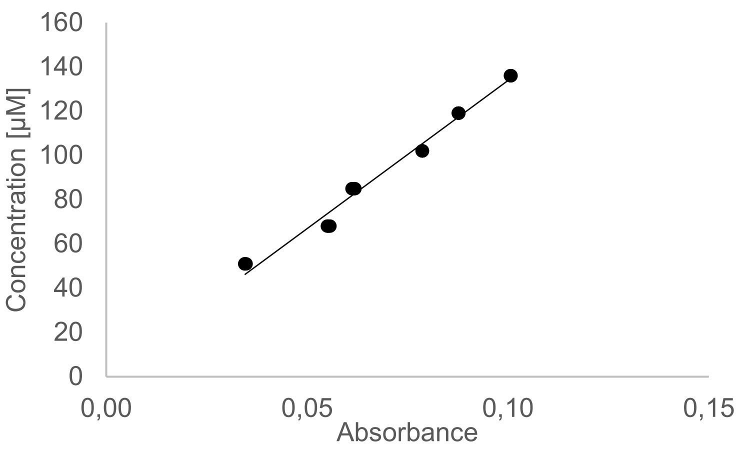
</td><td rowspan="3">136</td><td>0.101</td></tr>
<tr><td>0.101</td></tr>
<tr><td>0.101</td></tr>
<tr><td rowspan="3">119</td><td>0.088</td></tr>
<tr><td>0.088</td></tr>
<tr><td>0.088</td></tr>
<tr><td rowspan="3">102</td><td>0.079</td></tr>
<tr><td>0.079</td></tr>
<tr><td>0.079</td></tr>
<tr><td rowspan="3">85</td><td>0.062</td></tr>
<tr><td>0.062</td></tr>
<tr><td>0.061</td></tr>
<tr><td rowspan="3">68</td><td>0.055</td></tr>
<tr><td>0.056</td></tr>
<tr><td>0.055</td></tr>
<tr><td rowspan="3">$\mathbf{y = 1338.5x;\ R^2 = 0.9959}$</td><td rowspan="3">136</td><td>0.035</td></tr>
<tr><td>0.034</td></tr>
<tr><td>0.034</td></tr>
</table>

**Table S11.** Calibration curve of maximum absorbance value of FAD pH 6 using a spectrophotometer at λ = 450 nm. To each concentration point (y), three replicas were measured to obtain the corresponding absorbance values (x).

**FAD pH6**

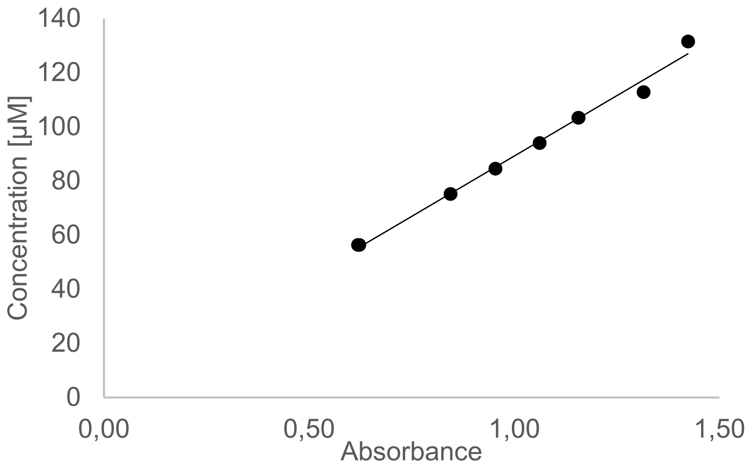


| *Concentration vs* $\mathbf{Abs_{max}}$ *(450nm)* | *Concentration (µM)* | *Absorbance* |
|---|---|---|
| | **131.6** | **1.424** |
| | | **1.424** |
| | | **1.424** |
| | **112.8** | **1.315** |
| | | **1.315** |
| | | **1.316** |
| | **94** | **1.156** |
| | | **1.157** |
| | | **1.157** |
| | **84.6** | **1.062** |
| | | **1.062** |
| | | **1.062** |
| | **75.2** | **0.955** |
| | | **0.954** |
| | | **0.954** |
| | | **0.844** |
| $\mathbf{y = 89.21x;\ R^2 = 0.9883}$ | **131.6** | **0.845** |
| | | **0.845** |

**Table S12.** Calibration curve of maximum absorbance value of FMN pH 6 using a spectrophotometer at λ = 448 nm. To each concentration point (y), three replicas were measured to obtain the corresponding absorbance values (x).

**FMN pH6**

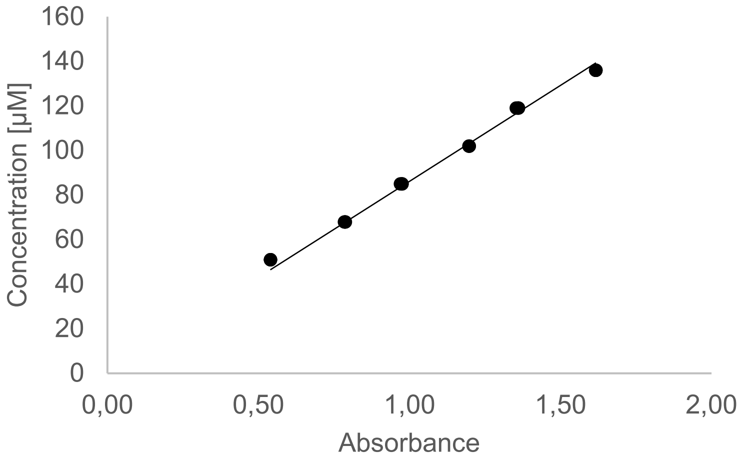


| *Concentration vs* $\mathbf{Abs_{max}}$ *(448nm)* | *Concentration (µM)* | *Absorbance* |
|---|---|---|
| | **136** | **1.617** |
| | | **1.617** |
| | | **1.617** |
| | **119** | **1.360** |
| | | **1.360** |
| | | **1.354** |
| | **102** | **1.198** |
| | | **1.198** |
| | | **1.197** |
| | **85** | **0.975** |
| | | **0.976** |
| | | **0.970** |
| | **68** | **0.786** |
| | | **0.788** |
| | | **0.785** |
| | | **0.540** |
| $\mathbf{y = 86.122x\ ;\ R^2 = 0.9924}$ | **51** | **0.539** |
| | | **0.539** |

**Table S13.** Calibration curve of minimum absorbance value of FAD pH 6 using a spectrophotometer at λ = 450 nm. To each concentration point (y), three replicas were measured to obtain the corresponding absorbance values (x).

<table>
<tr><th colspan="3">FAD pH6</th></tr>
<tr><td>Concentration vs $\mathbf{Abs_{min}}$ (450nm)</td><td>Concentration (µM)</td><td>Absorbance</td></tr>
<tr><td rowspan="15">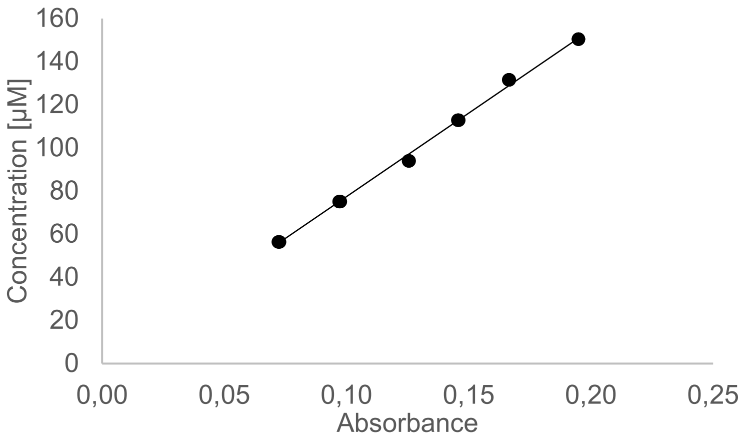
</td><td rowspan="3">150.4</td><td>0.195</td></tr>
<tr><td>0.195</td></tr>
<tr><td>0.195</td></tr>
<tr><td rowspan="3">131.6</td><td>0.166</td></tr>
<tr><td>0.167</td></tr>
<tr><td>0.166</td></tr>
<tr><td rowspan="3">112.8</td><td>0.146</td></tr>
<tr><td>0.146</td></tr>
<tr><td>0.145</td></tr>
<tr><td rowspan="3">94</td><td>0.126</td></tr>
<tr><td>0.125</td></tr>
<tr><td>0.125</td></tr>
<tr><td rowspan="3">75.2</td><td>0.097</td></tr>
<tr><td>0.097</td></tr>
<tr><td>0.097</td></tr>
<tr><td rowspan="3">$\mathbf{y = 774.29x;\ R^2 = 0.9971}$</td><td rowspan="3">56.4</td><td>0.072</td></tr>
<tr><td>0.073</td></tr>
<tr><td>0.072</td></tr>
</table>

**Table S14.** Calibration curve of minimum absorbance value of FMN pH 6 using a spectrophotometer at λ = 448 nm. To each concentration point (y), three replicas were measured to obtain the corresponding absorbance values (x).

<table>
<tr><th colspan="3">FMN pH6</th></tr>
<tr><td>Concentration vs $\mathbf{Abs_{min}}$ (448nm)</td><td>Concentration (µM)</td><td>Absorbance</td></tr>
<tr><td rowspan="15">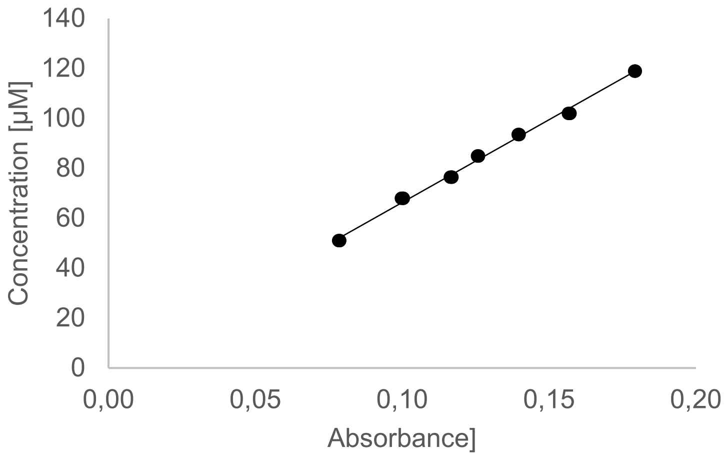
</td><td rowspan="3">119</td><td>0.179</td></tr>
<tr><td>0.179</td></tr>
<tr><td>0.179</td></tr>
<tr><td rowspan="3">102</td><td>0.157</td></tr>
<tr><td>0.157</td></tr>
<tr><td>0.157</td></tr>
<tr><td rowspan="3">93.5</td><td>0.140</td></tr>
<tr><td>0.140</td></tr>
<tr><td>0.140</td></tr>
<tr><td rowspan="3">85</td><td>0.126</td></tr>
<tr><td>0.126</td></tr>
<tr><td>0.126</td></tr>
<tr><td rowspan="3">76.5</td><td>0.117</td></tr>
<tr><td>0.117</td></tr>
<tr><td>0.116</td></tr>
<tr><td rowspan="3">$\mathbf{y = 663.03x;\ R^2 = 0.9959}$</td><td rowspan="3">68</td><td>0.100</td></tr>
<tr><td>0.100</td></tr>
<tr><td>0.100</td></tr>
</table>

**Table S15.** Calibration curve of maximum absorbance of riboflavin pH 10 using a spectrophotometer at λ = 446 nm. To each concentration point (y), three replicas were measured to obtain the corresponding absorbance values (x).

<table>
<tr><th colspan="3">Riboflavin pH10</th></tr>
<tr><th>Concentration vs $\mathbf{Abs_{max}}$ (446nm)</th><th>Concentration (μM)</th><th>Absorbance</th></tr>
<tr><td rowspan="15">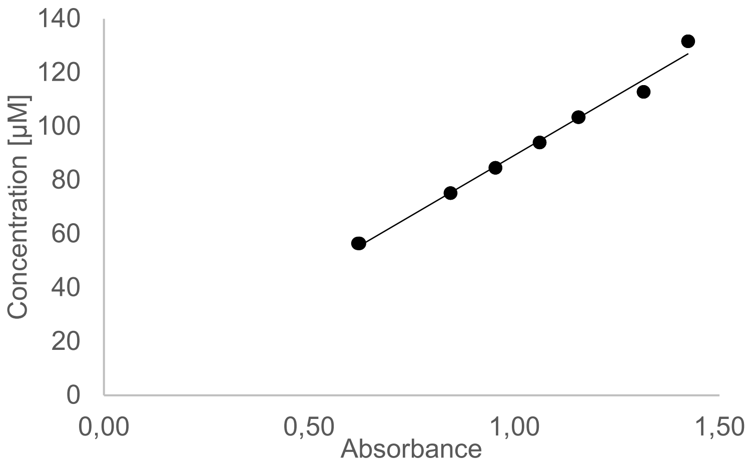
</td><td rowspan="3">160</td><td>1.7100</td></tr>
<tr><td>1.7015</td></tr>
<tr><td>1.6949</td></tr>
<tr><td rowspan="3">130</td><td>1.3435</td></tr>
<tr><td>1.3587</td></tr>
<tr><td>1.3559</td></tr>
<tr><td rowspan="3">100</td><td>1.1018</td></tr>
<tr><td>1.0845</td></tr>
<tr><td>1.0786</td></tr>
<tr><td rowspan="3">90</td><td>0.9708</td></tr>
<tr><td>0.9677</td></tr>
<tr><td>0.9642</td></tr>
<tr><td rowspan="3">80</td><td>0.8668</td></tr>
<tr><td>0.8759</td></tr>
<tr><td>0.8706</td></tr>
<tr><td rowspan="3">$\mathbf{y = 93.654x;\ R^2 = 0.996}$</td><td rowspan="3">70</td><td>0.7676</td></tr>
<tr><td>0.7681</td></tr>
<tr><td>0.7649</td></tr>
</table>

**Table S16.** Calibration curve of minimum absorbance of riboflavin pH 10 using a spectrophotometer at λ = 446 nm. To each concentration point (y), three replicas were measured to obtain the corresponding Absorbance values (x).

<table>
<tr><th colspan="3">Riboflavin pH10</th></tr>
<tr><th>Concentration vs $\mathbf{Abs_{min}}$ (446nm)</th><th>Concentration (μM)</th><th>Absorbance</th></tr>
<tr><td rowspan="15">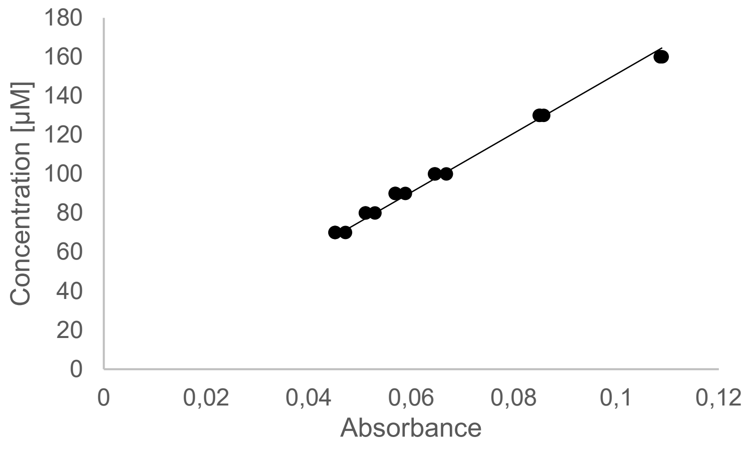
</td><td rowspan="3">160</td><td>0.1087</td></tr>
<tr><td>0.1086</td></tr>
<tr><td>0.1089</td></tr>
<tr><td rowspan="3">130</td><td>0.0850</td></tr>
<tr><td>0.0851</td></tr>
<tr><td>0.0859</td></tr>
<tr><td rowspan="3">100</td><td>0.0646</td></tr>
<tr><td>0.0645</td></tr>
<tr><td>0.0669</td></tr>
<tr><td rowspan="3">90</td><td>0.0569</td></tr>
<tr><td>0.0569</td></tr>
<tr><td>0.0589</td></tr>
<tr><td rowspan="3">80</td><td>0.0510</td></tr>
<tr><td>0.0511</td></tr>
<tr><td>0.0529</td></tr>
<tr><td rowspan="3">$\mathbf{y = 1511.2x;\ R^2 = 0.9924}$</td><td rowspan="3">70</td><td>0.0451</td></tr>
<tr><td>0.0452</td></tr>
<tr><td>0.0472</td></tr>
</table>

**Table S17.** Calibration curve of maximum absorbance of riboflavin pH 8 using a spectrophotometer at λ = 446 nm. To each concentration point (y), three replicas were measured to obtain the corresponding absorbance values (x).

<table>
<tr><th colspan="3">Riboflavin pH8</th></tr>
<tr><td>Concentration vs $\mathbf{Abs_{max}}$ (446nm)</td><td>Concentration (μM)</td><td>Absorbance</td></tr>
<tr><td rowspan="15">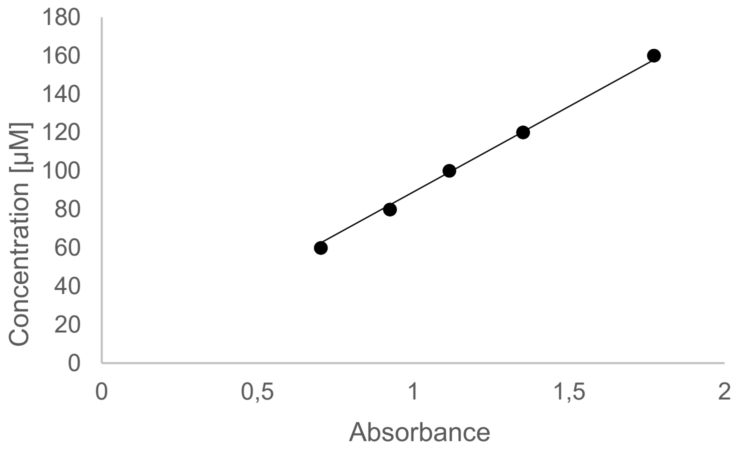
</td><td rowspan="3">160</td><td>1.7722</td></tr>
<tr><td>1.7723</td></tr>
<tr><td>1.7741</td></tr>
<tr><td rowspan="3">130</td><td>1.3534</td></tr>
<tr><td>1.3524</td></tr>
<tr><td>1.3527</td></tr>
<tr><td rowspan="3">100</td><td>1.1160</td></tr>
<tr><td>1.1160</td></tr>
<tr><td>1.1158</td></tr>
<tr><td rowspan="3">90</td><td>0.9251</td></tr>
<tr><td>0.9252</td></tr>
<tr><td>0.9256</td></tr>
<tr><td rowspan="3">80</td><td>0.7032</td></tr>
<tr><td>0.7035</td></tr>
<tr><td>0.7036</td></tr>
<tr><td rowspan="3">$\mathbf{y = 89.018x;\ R^2 = 0.997}$</td><td rowspan="3">70</td><td>1.7722</td></tr>
<tr><td>1.7723</td></tr>
<tr><td>1.7741</td></tr>
</table>

**Table S18.** Calibration curve of minimum absorbance of riboflavin pH 8 using a spectrophotometer at λ = 446 nm. To each concentration point (y), three replicas were measured to obtain the corresponding absorbance values (x).

<table>
<tr><th colspan="3">Riboflavin pH8</th></tr>
<tr><td>Concentration vs $\mathbf{Abs_{min}}$ (446nm)</td><td>Concentration (μM)</td><td>Absorbance</td></tr>
<tr><td rowspan="15">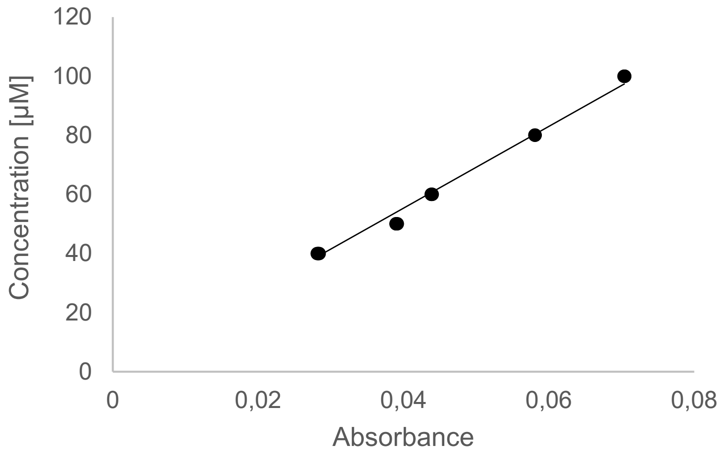
</td><td rowspan="3">100</td><td>0.0704</td></tr>
<tr><td>0.0704</td></tr>
<tr><td>0.0703</td></tr>
<tr><td rowspan="3">80</td><td>0.0581</td></tr>
<tr><td>0.0581</td></tr>
<tr><td>0.0581</td></tr>
<tr><td rowspan="3">60</td><td>0.0439</td></tr>
<tr><td>0.0439</td></tr>
<tr><td>0.0438</td></tr>
<tr><td rowspan="3">50</td><td>0.0390</td></tr>
<tr><td>0.0390</td></tr>
<tr><td>0.0391</td></tr>
<tr><td rowspan="3">40</td><td>0.0283</td></tr>
<tr><td>0.0284</td></tr>
<tr><td>0.0281</td></tr>
<tr><td colspan="3">$\mathbf{y = 1383.6x;\ R^2 = 0.9896}$</td></tr>
</table>

**Table S19.** Calibration curve of maximum absorbance of riboflavin pH 6 using a spectrophotometer at λ = 446 nm. To each concentration point (y), three replicas were measured to obtain the corresponding absorbance values (x).

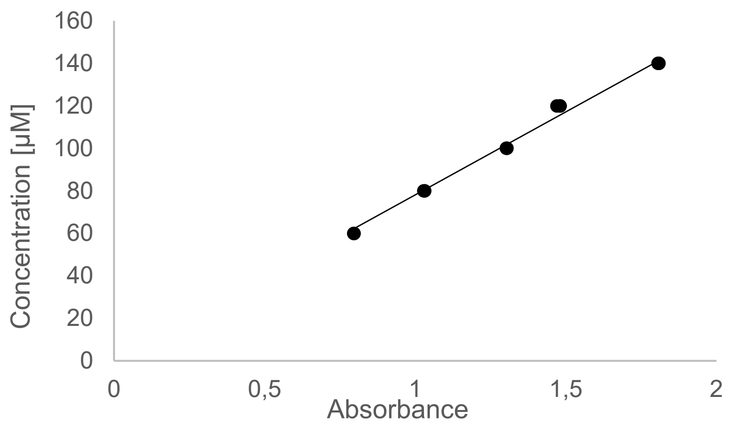


| Riboflavin pH6 | | |
|---|---|---|
| *Concentration vs* $\mathbf{Abs_{max}}$ *(446nm)* | *Concentration (µM)* | *Absorbance* |
| (graph above) | **140** | **1.8064** |
| | | **1.8092** |
| | | **1.8079** |
| | **120** | **1.4807** |
| | | **1.4808** |
| | | **1.4701** |
| | **100** | **1.3042** |
| | | **1.3045** |
| | | **1.3032** |
| | **80** | **1.0307** |
| | | **1.0318** |
| | | **1.0286** |
| | **60** | **0.7955** |
| | | **0.7968** |
| | | **0.7973** |
| $\mathbf{y = 78.104x;\ R^2 = 0.9921}$ | | |

**Table S20.** Calibration curve of minimum absorbance of riboflavin pH 6 using a spectrophotometer at λ = 446 nm. To each concentration point (y), three replicas were measured to obtain the corresponding absorbance values (x).

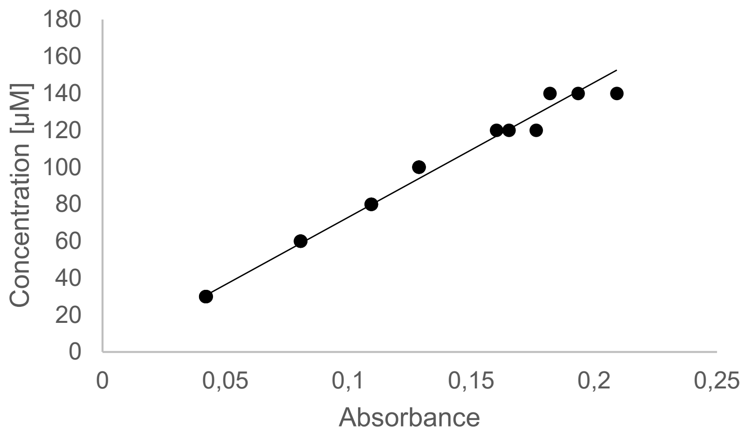


| Riboflavin pH6 | | |
|---|---|---|
| *Concentration vs* $\mathbf{Abs_{min}}$ *(446nm)* | *Concentration (µM)* | *Absorbance* |
| (graph above) | **140** | **0.1935** |
| | | **0.2092** |
| | | **0.1820** |
| | **120** | **0.1653** |
| | | **0.1764** |
| | | **0.1603** |
| | **100** | **0.1286** |
| | | **0.1287** |
| | | **0.1288** |
| | **80** | **0.1093** |
| | | **0.1093** |
| | | **0.1093** |
| | **60** | **0.0806** |
| | | **0.0806** |
| | | **0.0805** |
| $\mathbf{y = 729.43x;\ R^2 = 0.9829}$ | **30** | **0.0419** |
| | | **0.0419** |
| | | **0.0421** |

After reaction, the sample's absorbance value ($Abs_s$) was recorded in an anaerobic air-tight cuvette. Afterwards, the sample was reoxidised by opening it and mixing it with air. The spectra were collected again to obtain the maximum absorbance value ($Abs_{max}$).

**Quantification calculations.** After reoxidation, the sample's concentration ($\boldsymbol{c_s}$) was determined from its $\boldsymbol{Abs_{max}}$, which can be used to determine the theoretical $\boldsymbol{Abs_{min}}$ of the sample (see **Table S3.** ). Knowing the absorbance values of the fully reduced and oxidised sample, the amount of reduced Flavin within the sample can be determined according to **Eq. S3. ,** where $\mathbf{d}$ is the dilution factor of 40 (for FAD and FMN) or 1 (for Rib). This assumes that $\mathbf{c_s}$ is 100% pure flavin, which is an approximation. By comparing the sample to the control ($\mathbf{c_t}$), according to **Eq. S4.** , changes in the sample can be partly accounted for and discussed.

**Eq. S3.** Concentration of reduced flavin after the reaction (µM)

$$c_{red} = d * m_{max} * \left(\frac{Abs_{max} - Abs_s}{Abs_{max} - Abs_{min}}\right)$$

$$d_{FAD,FMN} = 40 \ ; \ d_{Rib} = 1$$

**Eq. S4.** Concentration difference between the sample and the control (µM)

$$\Delta c = d * m_{max} * (Abs_c - Abs_{max})$$

**Eq. S5.** Concentration of the control, described by the respective sample (µM)

$$c_t = c_{red} + c_{ox} + \Delta c$$

$c_{red}$ = concentration of reduced Flavin in the sample
$c_{ox}$ = concentration of oxidized Flavin in the sample
d = dilution fold of the sample for UV-Vis spectroscopy analysis
$m_{max}$ = extinction coefficient of the Flavin at a specific pH when fully oxidized
$Abs_{max}$ = absorbance of the diluted sample after fully oxidized
$Abs_c$ = absorbance of the diluted control after fully oxidized
$Abs_s$ = absorbance of the diluted sample after the reaction
$Abs_{min}$ = theoretical absorbance of the diluted sample if fully reduced, relative to calculated $c_s$
$c_t$ = concentration of the control
$\Delta_c$ = concentration difference between the sample and the control

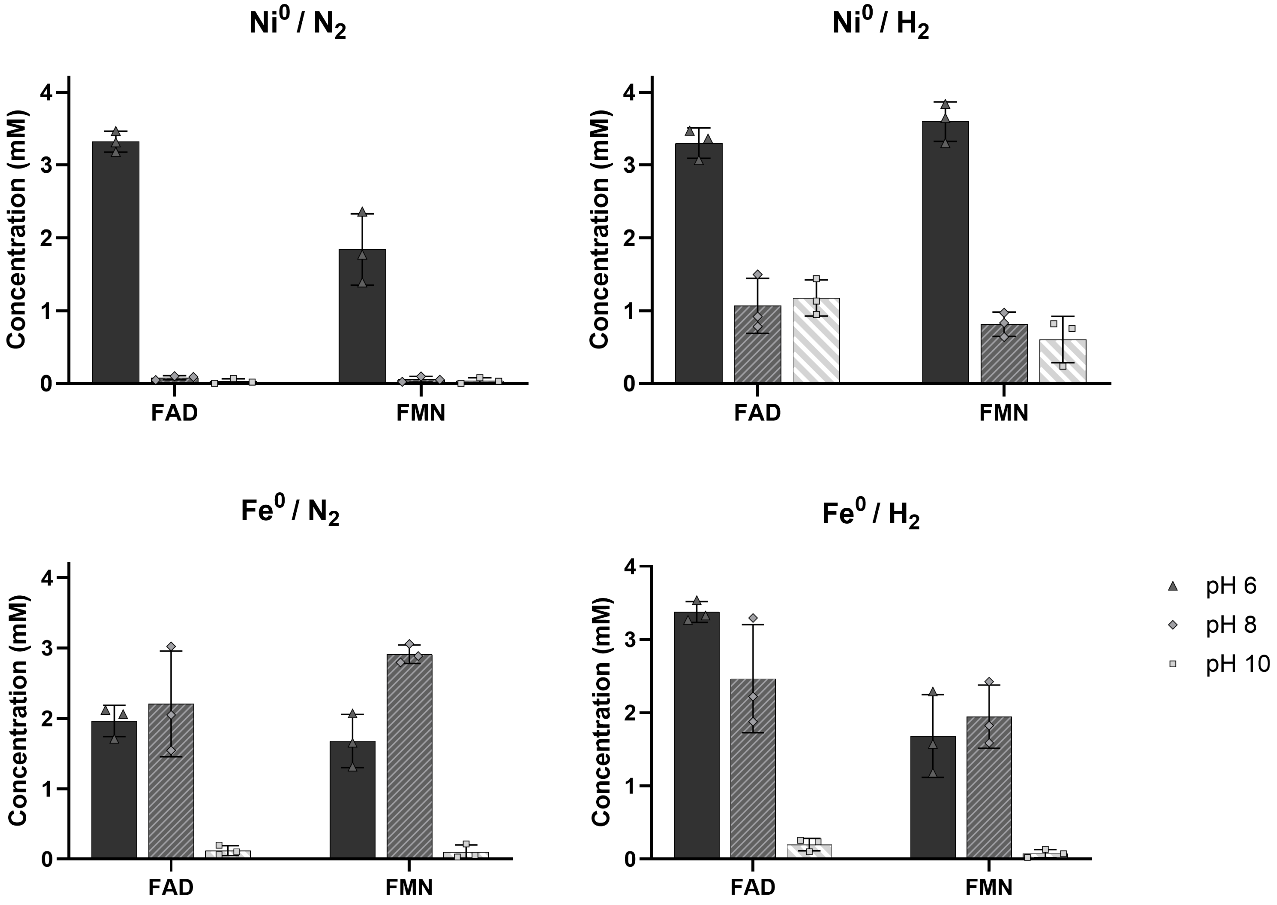


**Fig. S3.** Concentrations of reduced FAD and FMN after 15 minute reaction.

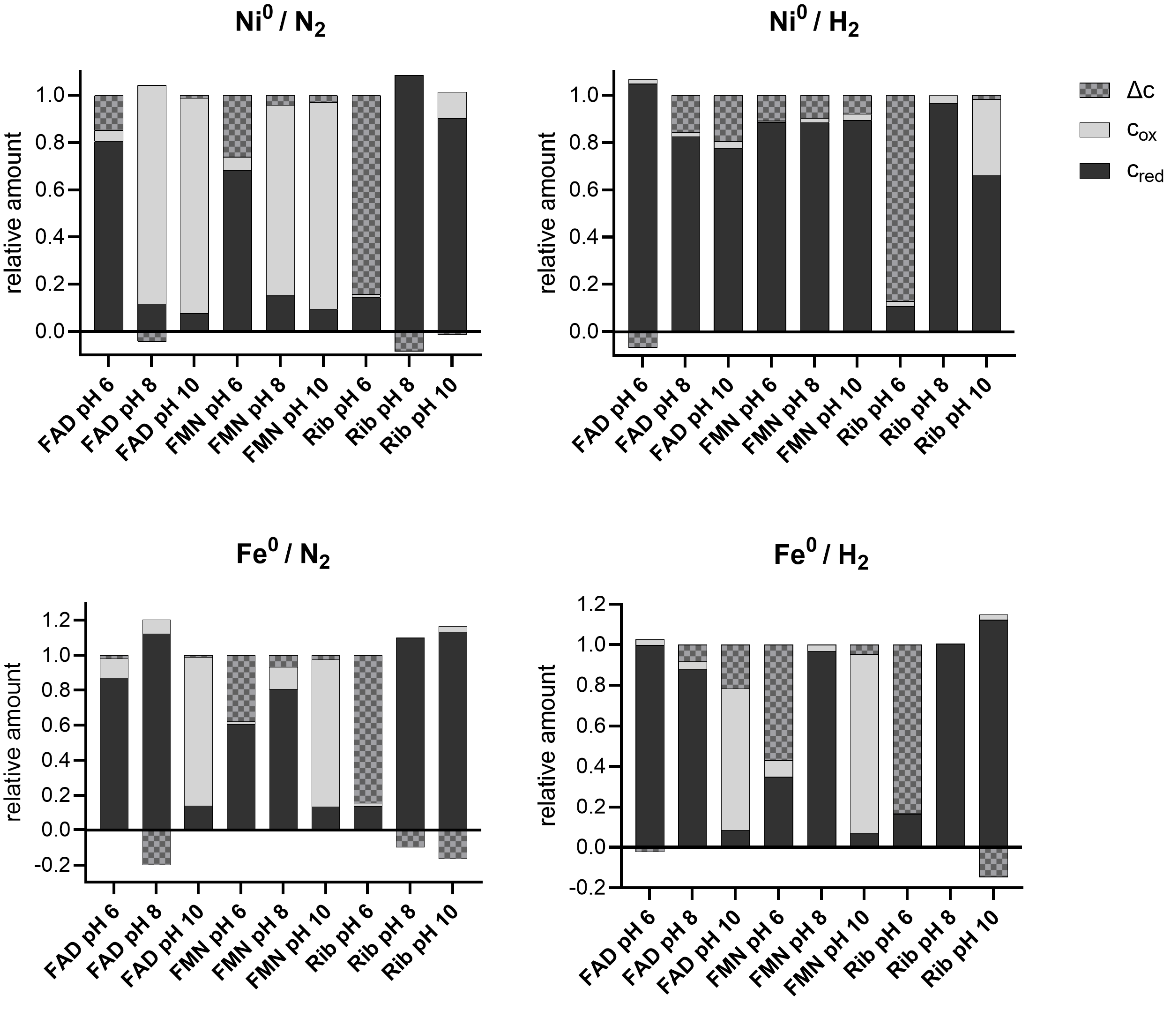


**Fig. S4**. FAD, FMN and Riboflavin's ratios of the reduced supernatant ($c_{red}$), oxidised ($c_{ox}$), and lost or gained flavin signal (Δc) after 2 hour reaction, in relation to the control sample. Values are according to **Table S21.**

**Table S21.** Post reaction absorbance values and pHs, and calculated concentration of the samples' reduced and oxidised proportions. The pH in the first column is that of the starting pH of the buffer, and in the pH on the second column is the pH of the sample after a 15 min or 2 h long reaction.

| Sample (metal+cofactor+pH+atmosphere+time) | pH after reaction | Sample Abs | Reoxidized sample Abs | Control Abs | Reduced (µM) | Oxidised (µM) | Control (µM) | Change (µM) |
|---|---|---|---|---|---|---|---|---|
| $Ni^0$ FAD pH 6 $H_2$ 15 min | 5.90 | 0.15 | 0.91 | 0.91 | 3068.49 | 177.60 | 3241.14 | -4.94 |
| | 5.77 | 0.16 | 0.99 | 0.91 | 3361.96 | 185.98 | 3241.14 | -306.79 |
| | 5.60 | 0.14 | 1.00 | 0.91 | 3469.82 | 93.48 | 3241.14 | -322.16 |
| $Ni^0$ FAD pH 8 $H_2$ 15 min | 7.78 | 0.84 | 1.04 | 1.03 | 782.10 | 2998.88 | 3733.38 | -47.60 |
| | 7.82 | 0.69 | 1.07 | 1.03 | 1494.82 | 2383.97 | 3733.38 | -145.41 |
| | 7.85 | 0.86 | 1.10 | 1.03 | 917.50 | 3061.85 | 3733.38 | -245.97 |
| $Ni^0$ FAD pH 10 $H_2$ 15 min | 9.86 | 0.68 | 0.99 | 1.01 | 1441.06 | 2827.93 | 4364.35 | 95.37 |
| | 9.86 | 0.80 | 1.04 | 1.01 | 1131.67 | 3360.80 | 4364.35 | -128.11 |
| | 9.87 | 0.84 | 1.04 | 1.01 | 948.05 | 3541.87 | 4364.35 | -125.56 |
| $Ni^0$ FAD pH 6 $H_2$ 15 min | 5.96 | 0.19 | 1.01 | 0.94 | 3314.12 | 298.13 | 3363.43 | -248.81 |
| | 5.98 | 0.17 | 0.96 | 0.94 | 3181.30 | 224.79 | 3363.43 | -42.66 |
| | 5.90 | 0.18 | 1.04 | 0.94 | 3468.37 | 229.28 | 3363.43 | -334.21 |
| $Ni^0$ FAD pH 8 $H_2$ 15 min | 7.90 | 0.93 | 0.96 | 0.94 | 89.94 | 3374.29 | 3408.69 | -55.54 |
| | 7.85 | 0.84 | 0.87 | 0.94 | 100.42 | 3044.32 | 3408.69 | 263.96 |
| | 7.85 | 0.78 | 0.79 | 0.94 | 47.09 | 2816.29 | 3408.69 | 545.31 |
| $Ni^0$ FAD pH 10 $H_2$ 15 min | 9.73 | 0.73 | 0.74 | 0.73 | 20.98 | 3160.57 | 3159.89 | -21.66 |
| | 9.73 | 0.69 | 0.70 | 0.73 | 67.47 | 2974.75 | 3159.89 | 117.67 |
| | 9.73 | 0.65 | 0.65 | 0.73 | -15.56 | 2800.00 | 3159.89 | 375.46 |
| $Ni^0$ FAD pH 6 $H_2$ 2 h | 5.82 | 0.12 | 0.97 | 0.83 | 3433.22 | 26.23 | 2969.69 | -489.76 |
| | 5.86 | 0.12 | 0.87 | 0.83 | 3055.99 | 57.67 | 2969.69 | -143.97 |
| | 5.82 | 0.12 | 0.82 | 0.83 | 2848.10 | 86.02 | 2969.69 | 35.56 |
| $Ni^0$ FAD pH 8 $H_2$ 2 h | 8.10 | 0.07 | 0.78 | 1.11 | 2754.07 | 68.78 | 4029.07 | 1206.23 |
| | 7.90 | 0.09 | 1.04 | 1.11 | 3681.12 | 72.84 | 4029.07 | 275.11 |
| | 7.90 | 0.08 | 1.00 | 1.11 | 3542.55 | 65.33 | 4029.07 | 421.19 |
| $Ni^0$ FAD pH 10 $H_2$ 2 h | 10.00 | 0.13 | 0.76 | 0.99 | 2951.30 | 329.47 | 4278.13 | 997.36 |
| | 9.90 | 0.07 | 0.81 | 0.99 | 3474.31 | 5.18 | 4278.13 | 798.64 |
| | 10.00 | 0.07 | 0.82 | 0.99 | 3546.05 | 15.14 | 4278.13 | 716.94 |
| $Ni^0$ FAD pH 6 $H_2$ 2 h | 5.93 | 0.13 | 0.97 | 1.05 | 3366.61 | 88.28 | 3729.80 | 274.91 |
| | 5.94 | 0.14 | 0.74 | 1.05 | 2392.07 | 234.16 | 3729.80 | 1103.57 |
| | 5.92 | 0.16 | 0.97 | 1.05 | 3258.14 | 195.70 | 3729.80 | 275.96 |
| $Ni^0$ FAD pH 8 $H_2$ 2 h | 8.07 | 0.87 | 0.97 | 1.00 | 401.47 | 3109.57 | 3637.48 | 126.44 |
| | 8.11 | 0.90 | 1.04 | 1.00 | 570.82 | 3208.42 | 3637.48 | -141.76 |
| | 8.16 | 1.05 | 1.13 | 1.00 | 299.43 | 3781.44 | 3637.48 | -443.39 |
| $Ni^0$ FAD pH 10 $H_2$ 2 h | 10.30 | 0.90 | 0.97 | 1.01 | 355.29 | 3847.88 | 4340.85 | 137.69 |
| | 10.08 | 0.96 | 1.01 | 1.01 | 271.69 | 4108.48 | 4340.85 | -39.31 |
| | 10.00 | 0.91 | 0.99 | 1.01 | 371.96 | 3907.66 | 4340.85 | 61.23 |

| | | | | | | | | |
|---|---|---|---|---|---|---|---|---|
| Fe$^0$ FAD pH 6 $H_2$ 15 min | 5.96 | 0.15 | 0.98 | 0.91 | 3326.70 | 157.20 | 3241.14 | -242.76 |
| | 6.06 | 0.18 | 0.99 | 0.91 | 3268.69 | 257.21 | 3241.14 | -284.76 |
| | 5.70 | 0.14 | 1.02 | 0.91 | 3538.10 | 93.67 | 3241.14 | -390.63 |
| Fe$^0$ FAD pH 8 $H_2$ 15 min | 8.30 | 0.46 | 1.04 | 1.03 | 2222.46 | 1528.92 | 3733.38 | -18.00 |
| | 8.50 | 0.17 | 1.02 | 1.03 | 3294.64 | 414.54 | 3733.38 | 24.20 |
| | 7.95 | 0.55 | 1.03 | 1.03 | 1879.77 | 1868.69 | 3733.38 | -15.09 |
| Fe$^0$ FAD pH 10 $H_2$ 15 min | 9.84 | 0.97 | 1.03 | 1.01 | 257.85 | 4180.27 | 4364.35 | -73.76 |
| | 9.86 | 0.97 | 1.02 | 1.01 | 238.30 | 4167.38 | 4364.35 | -41.33 |
| | 9.81 | 1.02 | 1.04 | 1.01 | 102.50 | 4393.94 | 4364.35 | -132.08 |
| Fe$^0$ FAD pH 6 $H_2$ 15 min | 6.00 | 0.09 | 0.60 | 0.94 | 2061.02 | 64.09 | 3363.43 | 1238.32 |
| | 5.96 | 0.10 | 0.63 | 0.94 | 2119.52 | 110.27 | 3363.43 | 1133.64 |
| | 5.98 | 0.42 | 0.84 | 0.94 | 1709.52 | 1291.43 | 3363.43 | 362.48 |
| Fe$^0$ FAD pH 8 $H_2$ 15 min | 8.26 | 0.12 | 0.89 | 0.94 | 3021.68 | 220.64 | 3408.69 | 166.37 |
| | 8.03 | 0.39 | 0.92 | 0.94 | 2049.67 | 1268.58 | 3408.69 | 90.44 |
| | 8.02 | 0.40 | 0.79 | 0.94 | 1545.92 | 1331.62 | 3408.69 | 531.15 |
| Fe$^0$ FAD pH 10 $H_2$ 15 min | 9.74 | 0.62 | 0.63 | 0.73 | 59.73 | 2658.29 | 3159.89 | 441.88 |
| | 9.73 | 0.90 | 0.94 | 0.73 | 197.54 | 3857.23 | 3159.89 | -894.88 |
| | 9.73 | 0.99 | 1.01 | 0.73 | 104.55 | 4250.13 | 3159.89 | -1194.78 |
| Fe$^0$ FAD pH 6 $H_2$ 2 h | 5.90 | 0.17 | 1.13 | 1.15 | 3899.95 | 139.75 | 4094.65 | 54.95 |
| | 5.91 | 0.17 | 1.22 | 1.15 | 4230.62 | 107.49 | 4094.65 | -243.45 |
| | 5.96 | 0.15 | 1.18 | 1.15 | 4129.75 | 69.93 | 4094.65 | -105.03 |
| Fe$^0$ FAD pH 8 $H_2$ 2 h | 8.28 | 0.14 | 1.05 | 1.11 | 3525.63 | 288.57 | 4029.07 | 214.87 |
| | 9.62 | 0.09 | 1.02 | 1.11 | 3627.52 | 82.77 | 4029.07 | 318.79 |
| | 9.55 | 0.09 | 0.98 | 1.11 | 3466.78 | 99.73 | 4029.07 | 462.56 |
| Fe$^0$ FAD pH 10 $H_2$ 2 h | 10.00 | 0.60 | 0.70 | 0.99 | 497.31 | 2543.65 | 4278.13 | 1237.17 |
| | 10.00 | 0.73 | 0.83 | 0.99 | 508.25 | 3091.60 | 4278.13 | 678.27 |
| | 10.00 | 0.77 | 0.79 | 0.99 | 79.22 | 3337.11 | 4278.13 | 861.79 |
| Fe$^0$ FAD pH 6 $N_2$ 2 h | 6.40 | 0.21 | 1.03 | 1.04 | 3329.75 | 355.16 | 3693.47 | 8.55 |
| | 6.34 | 0.24 | 1.03 | 1.04 | 3185.42 | 499.60 | 3693.47 | 8.45 |
| | 6.05 | 0.20 | 0.98 | 1.04 | 3137.96 | 364.61 | 3693.47 | 190.90 |
| Fe$^0$ FAD pH 8 $N_2$ 2 h | 9.55 | 0.17 | 1.32 | 1.01 | 4461.91 | 321.92 | 3656.962 | -1126.87 |
| | 9.85 | 0.16 | 1.13 | 1.01 | 3778.55 | 315.17 | 3656.96 | -436.76 |
| | 9.80 | 0.14 | 1.19 | 1.01 | 4086.76 | 229.86 | 3656.96 | -659.65 |
| Fe$^0$ FAD pH 10 $N_2$ 2 h | 10.00 | 0.79 | 0.95 | 1.01 | 710.00 | 3371.48 | 4340.85 | 259.38 |
| | 10.00 | 0.91 | 1.01 | 1.01 | 440.68 | 3911.97 | 4340.85 | -11.80 |
| | 10.01 | 0.89 | 1.03 | 1.01 | 680.92 | 3772.54 | 4340.85 | -112.60 |
| Ni$^0$ FMN pH 6 $N_2$ 15 min | 5.90 | 0.14 | 1.13 | 1.19 | 3643.94 | 233.90 | 4084.73 | 206.89 |
| | 5.95 | 0.13 | 1.18 | 1.19 | 3839.25 | 211.51 | 4084.73 | 33.97 |
| | 5.90 | 0.16 | 1.06 | 1.19 | 3303.92 | 335.41 | 4084.73 | 445.40 |
| Ni$^0$ FMN pH 8 $N_2$ 15 min | 7.72 | 1.26 | 1.48 | 0.99 | 638.40 | 3191.43 | 2561.83 | -1268.00 |
| | 7.73 | 1.22 | 1.51 | 0.99 | 830.08 | 3079.86 | 2561.83 | -1348.10 |
| | 7.74 | 1.20 | 1.54 | 0.99 | 972.67 | 3017.29 | 2561.83 | -1428.13 |
| | 9.96 | 1.02 | 1.27 | 1.34 | 819.01 | 3055.76 | 4072.44 | 197.67 |

| | | | | | | | | |
|---|---|---|---|---|---|---|---|---|
| $Ni^0$ FMN pH 10 $N_2$ 15 min | 9.99 | 1.26 | 1.33 | 1.34 | 236.35 | 3831.93 | 4072.44 | 4.16 |
| | 10.00 | 1.04 | 1.28 | 1.34 | 754.26 | 3145.28 | 4072.44 | 172.90 |
| $Ni^0$ FMN pH 6 $N_2$ 15 min | 5.92 | 0.25 | 0.89 | 0.81 | 2359.00 | 695.84 | 2803.95 | -250.90 |
| | 5.92 | 0.14 | 0.62 | 0.81 | 1769.88 | 366.64 | 2803.95 | 667.43 |
| | 5.93 | 0.15 | 0.52 | 0.81 | 1389.05 | 413.71 | 2803.95 | 1001.19 |
| $Ni^0$ FMN pH 8 $N_2$ 15 min | 7.78 | 1.00 | 1.00 | 1.32 | 22.41 | 2578.43 | 3412.09 | 811.25 |
| | 7.80 | 1.19 | 1.22 | 1.32 | 96.93 | 3075.09 | 3412.09 | 240.06 |
| | 7.74 | 1.24 | 1.26 | 1.32 | 52.43 | 3219.38 | 3412.09 | 140.27 |
| $Ni^0$ FMN pH 10 $N_2$ 15 min | 9.90 | 0.80 | 0.82 | 1.19 | 80.03 | 2428.60 | 3619.37 | 1110.74 |
| | 9.90 | 0.82 | 0.81 | 1.19 | -40.58 | 2507.22 | 3619.37 | 1152.73 |
| | 9.90 | 0.82 | 0.83 | 1.19 | 29.59 | 2507.41 | 3619.37 | 1082.37 |
| $Ni^0$ FMN pH 6 $N_2$ 2 h | 5.87 | 0.15 | 0.91 | 1.29 | 2833.02 | 315.67 | 4434.93 | 1286.23 |
| | 5.83 | 0.16 | 1.28 | 1.29 | 4146.64 | 276.49 | 4434.93 | 11.80 |
| | 5.84 | 0.17 | 1.25 | 1.29 | 3985.99 | 323.08 | 4434.93 | 125.87 |
| $Ni^0$ FMN pH 8 $N_2$ 2 h | 7.76 | 0.09 | 1.27 | 1.36 | 3372.40 | -89.33 | 3514.13 | 231.06 |
| | 7.82 | 0.10 | 1.16 | 1.36 | 3064.38 | -47.46 | 3514.13 | 497.21 |
| | 7.77 | 0.12 | 1.31 | 1.36 | 3413.40 | -23.40 | 3514.13 | 124.12 |
| $Ni^0$ FMN pH 10 $N_2$ 2 h | 9.97 | 0.08 | 1.26 | 1.38 | 3788.89 | 55.27 | 4220.87 | 376.72 |
| | 10.00 | 0.08 | 1.36 | 1.38 | 4073.84 | 59.49 | 4220.87 | 87.55 |
| | 9.92 | 0.07 | 1.14 | 1.38 | 3446.28 | 38.44 | 4220.87 | 736.15 |
| $Ni^0$ FMN pH 6 $N_2$ 2 h | 5.97 | 0.17 | 1.06 | 1.25 | 3277.66 | 383.08 | 4314.17 | 140.72 |
| | 5.94 | 0.19 | 0.90 | 1.25 | 2605.21 | 486.82 | 4314.17 | 1059.11 |
| | 5.93 | 0.18 | 0.82 | 1.25 | 2343.27 | 471.84 | 4314.17 | 1473.15 |
| $Ni^0$ FMN pH 8 $N_2$ 2 h | 7.77 | 1.11 | 1.22 | 1.31 | 329.32 | 2840.02 | 3380.59 | 211.25 |
| | 7.85 | 1.18 | 1.28 | 1.31 | 295.03 | 3022.51 | 3380.59 | 63.06 |
| | 7.86 | 0.90 | 1.25 | 1.31 | 999.51 | 2235.41 | 3380.59 | 145.67 |
| $Ni^0$ FMN pH 10 $N_2$ 2 h | 9.96 | 0.93 | 1.07 | 1.13 | 432.02 | 2826.51 | 3435.33 | 176.79 |
| | 9.98 | 1.02 | 1.12 | 1.13 | 323.59 | 3079.06 | 3435.33 | 32.68 |
| | 9.98 | 1.02 | 1.09 | 1.13 | 231.89 | 3087.81 | 3435.33 | 115.63 |
| $Fe^0$ FMN pH 6 $N_2$ 15 min | 5.93 | 0.26 | 0.69 | 1.19 | 1576.63 | 803.72 | 4084.73 | 1704.38 |
| | 5.93 | 0.43 | 0.75 | 1.19 | 1180.78 | 1398.92 | 4084.73 | 1505.03 |
| | 5.99 | 0.08 | 0.71 | 1.19 | 2292.58 | 136.07 | 4084.73 | 1656.08 |
| $Fe^0$ FMN pH 8 $N_2$ 15 min | 7.96 | 0.43 | 1.28 | 0.99 | 2425.17 | 887.84 | 2561.83 | -751.18 |
| | 7.97 | 0.61 | 1.25 | 0.99 | 1823.76 | 1410.84 | 2561.83 | -672.78 |
| | 8.00 | 0.72 | 1.27 | 0.99 | 1588.25 | 1705.73 | 2561.83 | -732.16 |
| $Fe^0$ FMN pH 10 $N_2$ 15 min | 9.98 | 1.30 | 1.32 | 1.34 | 75.28 | 3961.88 | 4072.44 | 35.28 |
| | 9.98 | 1.28 | 1.29 | 1.34 | 24.23 | 3915.54 | 4072.44 | 132.66 |
| | 9.98 | 1.30 | 1.34 | 1.34 | 133.86 | 3945.05 | 4072.44 | -6.47 |
| $Fe^0$ FMN pH 6 $N_2$ 15 min | 5.94 | 0.23 | 0.68 | 0.81 | 1652.94 | 676.22 | 2803.95 | 474.78 |
| | 5.92 | 0.06 | 0.42 | 0.81 | 1315.84 | 123.22 | 2803.95 | 1364.88 |
| | 5.92 | 0.19 | 0.75 | 0.81 | 2069.25 | 514.26 | 2803.95 | 220.44 |
| | 8.16 | 0.17 | 1.17 | 1.32 | 2884.81 | 157.56 | 3412.09 | 369.72 |

| | | | | | | | | |
|---|---|---|---|---|---|---|---|---|
| $Fe^0$ FMN pH 8 $N_2$ 15 min | 8.00 | 0.15 | 1.21 | 1.32 | 3055.60 | 82.73 | 3412.09 | 273.76 |
| | 7.86 | 0.19 | 1.17 | 1.32 | 2795.49 | 227.12 | 3412.09 | 389.48 |
| $Fe^0$ FMN pH 10 $N_2$ 15 min | 9.90 | 0.68 | 0.75 | 1.19 | 216.14 | 2066.47 | 3619.37 | 1336.77 |
| | 9.90 | 0.85 | 0.86 | 1.19 | 28.56 | 2588.85 | 3619.37 | 1001.96 |
| | 9.90 | 0.82 | 0.84 | 1.19 | 54.88 | 2507.04 | 3619.37 | 1057.45 |
| $Fe^0$ FMN pH 6 $N_2$ 2 h | 5.92 | 0.17 | 0.57 | 1.29 | 1464.07 | 481.38 | 4434.93 | 2489.48 |
| | 5.75 | 0.14 | 0.54 | 1.29 | 1461.25 | 382.75 | 4434.93 | 2590.93 |
| | 5.84 | 0.18 | 0.56 | 1.29 | 1388.27 | 528.26 | 4434.93 | 2518.40 |
| $Fe^0$ FMN pH 8 $N_2$ 2 h | 8.13 | 0.13 | 1.37 | 1.36 | 3564.68 | -17.58 | 3514.13 | -32.98 |
| | 8.44 | 0.10 | 1.31 | 1.36 | 3484.78 | -89.95 | 3514.13 | 119.29 |
| | 9.02 | 0.09 | 1.39 | 1.36 | 3717.73 | -118.81 | 3514.13 | -84.79 |
| $Fe^0$ FMN pH 10 $N_2$ 2 h | 9.96 | 1.14 | 1.35 | 1.38 | 678.37 | 3444.40 | 4220.87 | 98.10 |
| | 9.94 | 1.34 | 1.36 | 1.38 | 78.64 | 4081.88 | 4220.87 | 60.35 |
| | 9.92 | 1.22 | 1.24 | 1.38 | 82.51 | 3705.02 | 4220.87 | 433.34 |
| $Fe^0$ FMN pH 6 $N_2$ 2 h | 5.96 | 0.09 | 0.56 | 1.25 | 1737.99 | 188.82 | 4314.17 | 2387.36 |
| | 6.00 | 0.16 | 0.91 | 1.25 | 2756.91 | 367.49 | 4314.17 | 1189.77 |
| | 6.08 | 0.12 | 0.88 | 1.25 | 2795.44 | 223.46 | 4314.17 | 1295.27 |
| $Fe^0$ FMN pH 8 $N_2$ 2 h | 8.26 | 0.24 | 1.27 | 1.31 | 2946.23 | 332.17 | 3380.59 | 102.19 |
| | 8.43 | 0.22 | 1.23 | 1.31 | 2875.77 | 299.37 | 3380.59 | 205.45 |
| | 8.66 | 0.18 | 1.16 | 1.31 | 2811.22 | 194.62 | 3380.59 | 374.75 |
| $Fe^0$ FMN pH 10 $N_2$ 2 h | 9.98 | 0.92 | 1.11 | 1.13 | 608.45 | 2781.89 | 3435.33 | 44.99 |
| | 9.97 | 0.98 | 1.09 | 1.13 | 351.40 | 2975.61 | 3435.33 | 108.32 |
| | 9.97 | 0.96 | 1.09 | 1.13 | 437.57 | 2891.44 | 3435.33 | 106.32 |
| $Ni^0$ Riboflavin pH 6 $N_2$ 2 h | 6.06 | 0.04 | 0.17 | 729.43 | 10.76 | 2.14 | 98.61 | 85.71 |
| | 6.05 | 0.05 | 0.17 | 729.43 | 10.59 | 2.70 | 98.61 | 85.31 |
| | 5.99 | 0.04 | 0.15 | 729.43 | 10.22 | 1.65 | 98.61 | 86.74 |
| $Ni^0$ Riboflavin pH 6 $N_2$ 2 h | 6.09 | 0.04 | 0.20 | 729.43 | 13.90 | 1.74 | 93.92 | 78.27 |
| | 6.10 | 0.04 | 0.19 | 729.43 | 13.28 | 1.38 | 93.92 | 79.26 |
| | 6.11 | 0.03 | 0.18 | 729.43 | 13.59 | 0.49 | 93.92 | 79.83 |
| $Ni^0$ Riboflavin pH 10 $N_2$ 2 h | 10.32 | 0.09 | 1.09 | 1511.20 | 99.59 | 2.62 | 95.19 | -7.02 |
| | 10.30 | 0.11 | 1.11 | 1511.20 | 100.69 | 3.57 | 95.19 | -9.07 |
| | 10.34 | 0.10 | 1.08 | 1511.20 | 97.50 | 3.13 | 95.19 | -5.45 |
| $Ni^0$ Riboflavin pH 10 $N_2$ 2 h | 10.29 | 0.25 | 1.08 | 1511.20 | 82.37 | 18.47 | 93.14 | -7.70 |
| | 10.28 | 0.27 | 1.10 | 1511.20 | 82.09 | 20.42 | 93.14 | -9.36 |
| | 10.23 | 0.23 | 1.11 | 1511.20 | 87.69 | 16.17 | 93.14 | -10.72 |
| $Fe^0$ Riboflavin pH 6 $N_2$ 2 h | 6.28 | 0.02 | 0.20 | 729.43 | 15.10 | 0.21 | 98.61 | 83.30 |
| | 6.37 | 0.04 | 0.22 | 729.43 | 15.80 | 1.32 | 98.61 | 81.49 |
| | 6.28 | 0.02 | 0.22 | 729.43 | 17.02 | -0.19 | 98.61 | 81.78 |
| $Fe^0$ Riboflavin pH 6 $N_2$ 2 h | 6.31 | 0.04 | 0.18 | 729.43 | 11.76 | 2.09 | 93.92 | 80.08 |
| | 6.27 | 0.04 | 0.19 | 729.43 | 13.40 | 1.49 | 93.92 | 79.03 |
| | 6.39 | 0.04 | 0.20 | 729.43 | 13.51 | 1.75 | 93.92 | 78.66 |
| | 10.25 | 0.08 | 1.09 | 1511.20 | 100.90 | 0.97 | 95.19 | -6.69 |

| | | | | | | | | |
|---|---|---|---|---|---|---|---|---|
| $Fe^0$ Riboflavin pH 10 $N_2$ 2 h | 10.30 | 0.10 | 1.10 | 1511.20 | 100.22 | 3.12 | 95.19 | -8.15 |
| | 10.29 | 0.08 | 1.11 | 1511.20 | 103.04 | 1.20 | 95.19 | -9.06 |
| $Fe^0$ Riboflavin pH 10 $N_2$ 2 h | 10.22 | 0.08 | 1.10 | 1511.20 | 102.70 | 0.69 | 93.14 | -10.25 |
| | 10.28 | 0.07 | 1.09 | 1511.20 | 101.11 | 0.56 | 93.14 | -8.52 |
| | 10.26 | 0.09 | 1.11 | 1511.20 | 102.48 | 1.77 | 93.14 | -11.11 |
| $Ni^0$ Riboflavin pH 8 $N_2$ 2 h | 8.00 | 0.11 | 0.97 | 1383.60 | 90.95 | 4.27 | 94.49 | -0.73 |
| | 8.00 | 0.08 | 0.93 | 1383.60 | 90.24 | 1.16 | 94.49 | 3.08 |
| | 8.00 | 0.10 | 0.98 | 1383.60 | 92.81 | 3.42 | 94.49 | -1.74 |
| $Ni^0$ Riboflavin pH 8 $N_2$ 2 h | 8.00 | 0.07 | 0.98 | 1383.60 | 96.10 | -0.26 | 86.05 | -9.79 |
| | 8.00 | 0.07 | 0.94 | 1383.60 | 91.76 | 0.36 | 86.05 | -6.07 |
| | 8.00 | 0.07 | 0.94 | 1383.60 | 91.92 | -0.17 | 86.05 | -5.70 |
| $Fe^0$ Riboflavin pH 8 $N_2$ 2 h | 8.50 | 0.07 | 0.97 | 1383.60 | 94.89 | 0.41 | 94.49 | -0.81 |
| | 8.50 | 0.07 | 0.96 | 1383.60 | 93.75 | -0.13 | 94.49 | 0.87 |
| | 8.50 | 0.07 | 0.98 | 1383.60 | 96.13 | -0.21 | 94.49 | -1.43 |
| $Fe^0$ Riboflavin pH 8 $N_2$ 2 h | 8.70 | 0.06 | 0.97 | 1383.60 | 95.66 | -0.51 | 86.05 | -9.09 |
| | 8.70 | 0.07 | 0.96 | 1383.60 | 93.20 | 0.50 | 86.05 | -7.65 |
| | 8.70 | 0.08 | 0.97 | 1383.60 | 94.61 | 0.76 | 86.05 | -9.33 |

### Reactivity with different metals

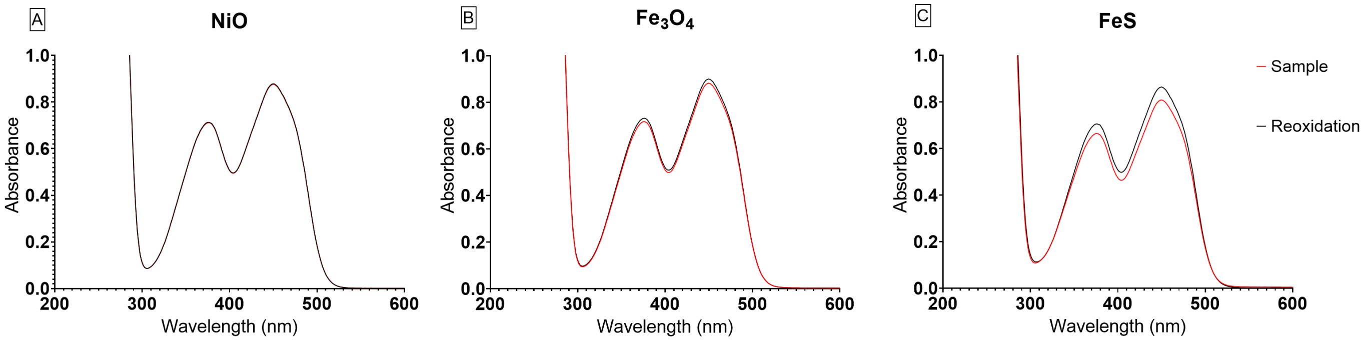


**Fig S5.** UV-Vis spectra (200–600 nm) of FAD after 4h, at pH 8, 0.133 M phosphate buffer, at 40 °C, under 5 bars of $H_2$, with 18 mM of NiO (A), $Fe_3O_4$ (B) and FeS (C). The red line of the graphs shows the samples' lowest spectrum after the reaction. The black line is the absorbance spectra after reoxidation.

### Reaction Mechanism

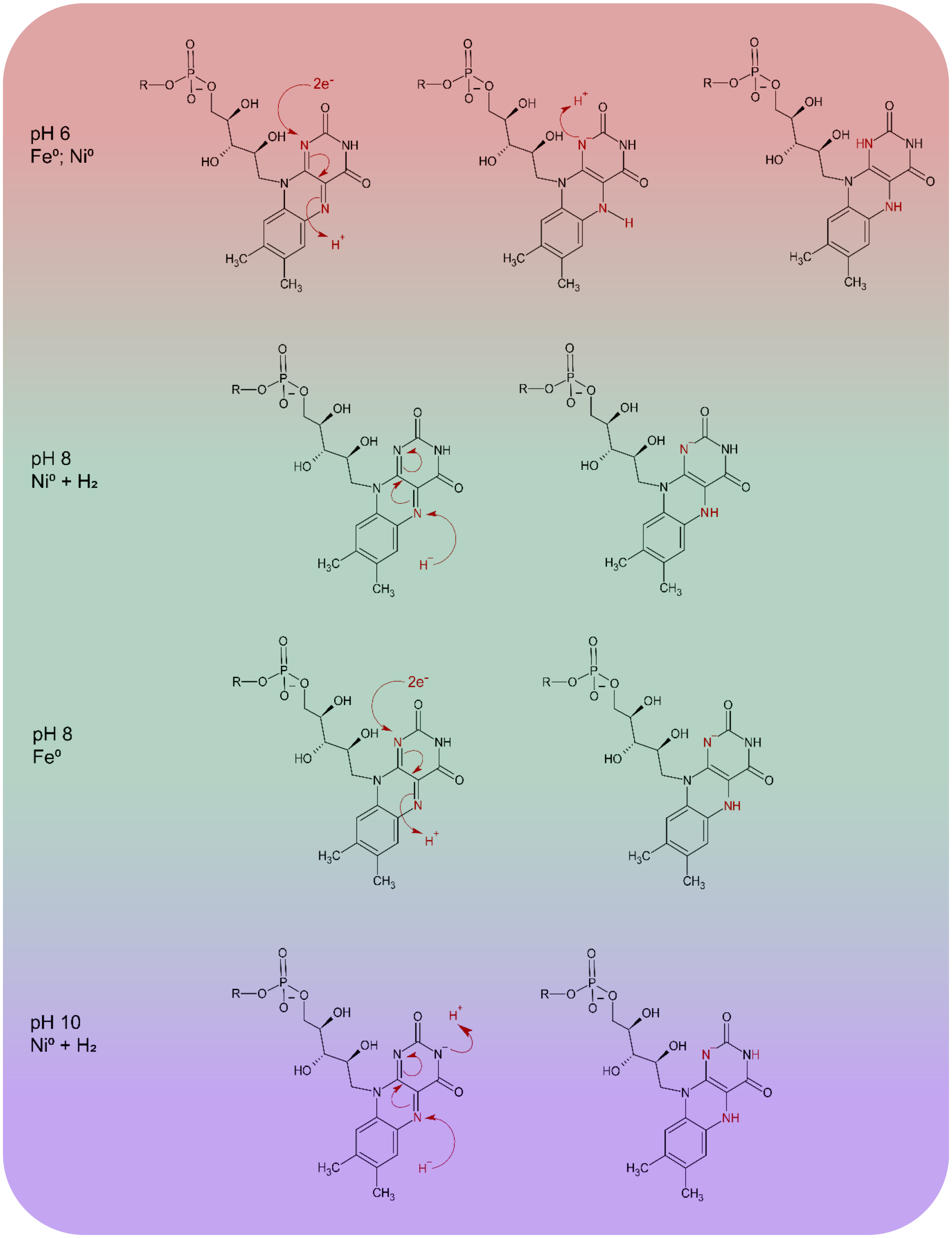


**Fig. S6.** Proposed reaction mechanisms for each pH and metal where significant reduction was observed. Considering the most common species of oxidized flavin that exist at pHs 6, 8 and 10, the reduction reaction was designed considering the dominant electron donor (metal or $H_2$) and the species of reduced Flavin at the same pH. In the case of FAD, the group R is adenosine monophosphate, while for FMN it is either a proton at pH 6, or nothing at pHs 8 and 10.

### Cyclic Voltammetry

Standards of 2 mM FAD and FMN were characterized at pHs 6, 8, and 10 through cyclic voltammetry with the corresponding buffers for the reactions (see **Fig. S6**).The measurements were performed at room temperature, under a $N_2$ atmosphere and purged solutions. The electrochemical cell consisted of three electrodes: glassy carbon as the working electrode, platinum wire as the counter electrode, and Ag/AgCl (3 M NaCl) as the reference electrode. The scan rate was set to 100 mV/s, starting from a cathodic current.

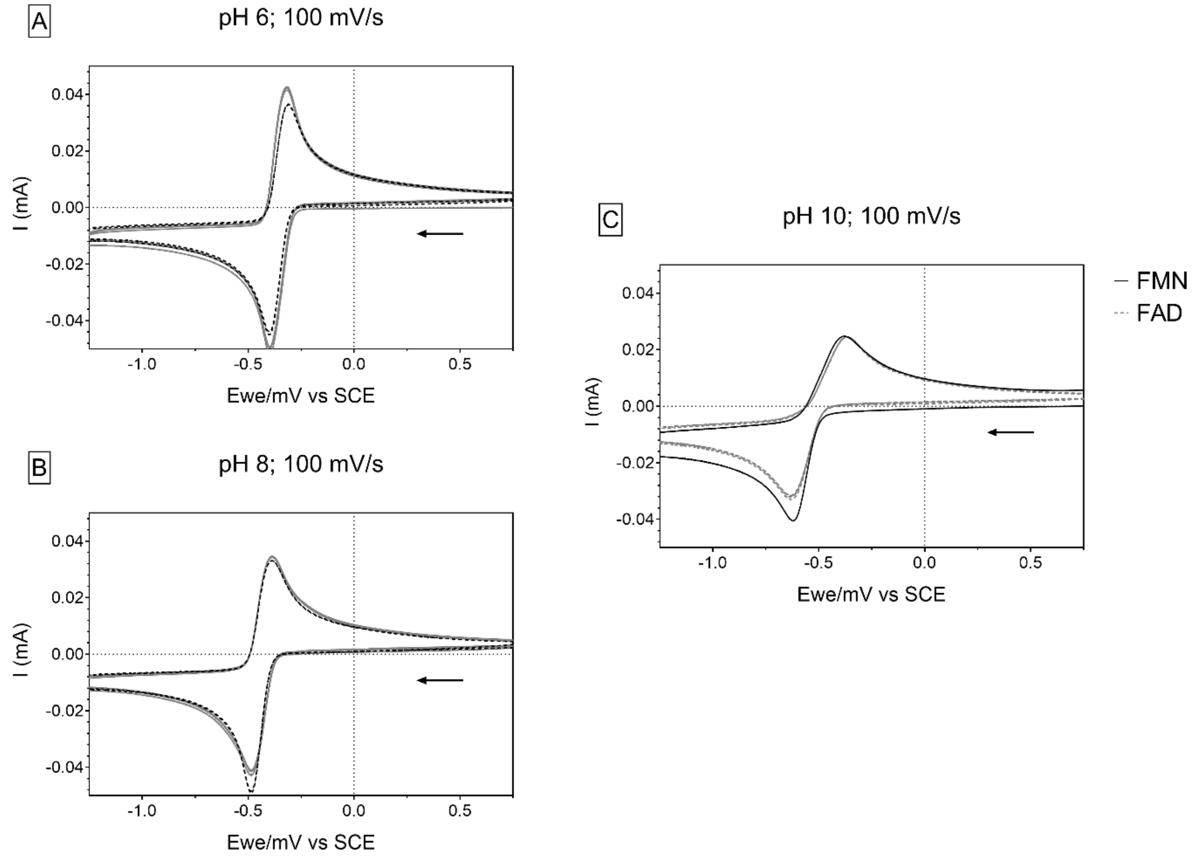


**Fig. S7.** Cyclic voltammetry graphs of 2 mM FAD (gray dashed line) and FMN (black solid line) in pHs of 6 (A), 8 (B), and 10 (C). For each measurement three cycles were completed as displayed, starting from the cathodic current.

### Gas Chromatography Thermal Conductivity Detection (GC-TCD).

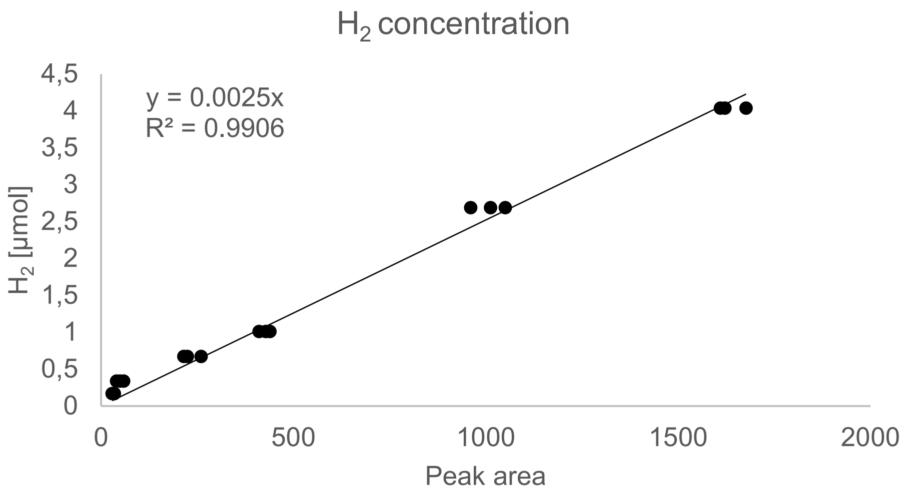


**Fig. S8.** A hydrogen calibration curve to quantify the produced hydrogen in the samples. On the x-axis is the peak area, and on the y-axis is the amount of hydrogen in moles, according to **Table S22**. The measurements were made from an injected ratio of atmospheric oxygen and stock hydrogen.

**Table S22.** Table of hydrogen's calibration curve data points. The moles of hydrogen were obtained from **Eq. S6.**

| $H_2$ + Atmosphere (µL) | injected (µL) | Hydrogen (µmol) | peak area | Ratio of hydrogen volume |
|---|---|---|---|---|
| 200+400 | 200 | 2.690806429 | 960.5636 | 0.333333333 |
| 200+400 | 200 | 2.690806429 | 1050.3232 | 0.333333333 |
| 200+400 | 200 | 2.690806429 | 1012.1944 | 0.333333333 |
| 200+400 | 300 | 4.036209644 | 1621.519 | 0.333333333 |
| 200+400 | 300 | 4.036209644 | 1676.6072 | 0.333333333 |
| 200+400 | 300 | 4.036209644 | 1609.7978 | 0.333333333 |
| 100+500 | 100 | 0.672701607 | 214.878 | 0.166666667 |
| 100+500 | 100 | 0.672701607 | 224.3788 | 0.166666667 |
| 100+500 | 100 | 0.672701607 | 259.7712 | 0.166666667 |
| 100+500 | 50 | 0.336350804 | 49.661 | 0.166666667 |
| 100+500 | 50 | 0.336350804 | 40.3538 | 0.166666667 |
| 100+500 | 50 | 0.336350804 | 58.5595 | 0.166666667 |
| 50+550 | 50 | 0.168175402 | 28.8734 | 0.083333333 |
| 50+550 | 50 | 0.168175402 | 34.3241 | 0.083333333 |
| 50+550 | 50 | 0.168175402 | 34.0682 | 0.083333333 |
| 50+550 | 300 | 1.009052411 | 439.14 | 0.083333333 |
| 50+550 | 300 | 1.009052411 | 427.9248 | 0.083333333 |
| 50+550 | 300 | 1.009052411 | 410.7557 | 0.083333333 |

**Eq. S6.** Ideal gas law

$$PV = nRT$$

P = pressure
V = volume
n = moles
R = gas constant
T = temperature

**Eq. S7.** Henry's law

$$C = Pk_H$$

C = dissolved gas concentration
P = partial pressure of the gas
$k_H$ = Henry's law constant ($7.8*10^{-4}$ mol $L^{-1}$ $atm^{-1}$)

**Table S23.** Table of standard samples' hydrogen content after a reaction. The caps were sealed with electrical tape to trap the released $H_2$ inside the sample vial. The molarity was calculated using **Fig. S11**.

| Sample (metal+cofactor +pH) | Peak area | Injected (μL) | Hydrogen in solution (μmol) | Hydrogen in headspace (μmol) | Total hydrogen (μmol) |
|---|---|---|---|---|---|
| $Ni^0$ FAD pH 6 | 29.8100 | 200 | 0.0013 | 0.0007 | 0.0020 |
| | 33.3900 | 200 | 0.0014 | 0.0008 | 0.0023 |
| | 2.6429 | 100 | 0.0001 | 0.0001 | 0.0002 |
| FAD pH 6 | 0.0000 | 100 | 0.0000 | 0.0000 | 0.0000 |
| $Ni^0$ FMN pH 6 | 2.4507 | 200 | 0.0001 | 0.0001 | 0.0002 |
| | 6.4377 | 200 | 0.0003 | 0.0002 | 0.0004 |
| | 1.0328 | 200 | 0.0000 | 0.0000 | 0.0001 |
| FMN pH 6 | 0.0000 | 100 | 0.0000 | 0.0000 | 0.0000 |
| pH 6 buffer | 0.0000 | 100 | 0.0000 | 0.0000 | 0.0000 |
| $Fe^0$ FAD pH 6 | 281.0780 | 100 | 0.0134 | 0.0070 | 0.0204 |
| | 275.8614 | 100 | 0.0132 | 0.0069 | 0.0200 |
| | 183.7432 | 100 | 0.0088 | 0.0046 | 0.0134 |
| $Fe^0$ FMN pH 6 | 108.5978 | 100 | 0.0052 | 0.0027 | 0.0079 |
| | 83.8232 | 100 | 0.0040 | 0.0021 | 0.0061 |
| | 223.5010 | 100 | 0.0107 | 0.0056 | 0.0162 |
| $Ni^0$ FAD pH 8 | 12.2906 | 200 | 0.0005 | 0.0003 | 0.0008 |
| | 1.9507 | 200 | 0.0001 | 0.0000 | 0.0001 |
| | 1.1682 | 200 | 0.0001 | 0.0000 | 0.0001 |
| FAD pH 8 | 0.0000 | 200 | 0.0000 | 0.0000 | 0.0000 |
| $Ni^0$ FMN pH 8 | 2.4798 | 200 | 0.0001 | 0.0001 | 0.0002 |
| | 2.7848 | 200 | 0.0001 | 0.0001 | 0.0002 |
| | 1.2832 | 200 | 0.0001 | 0.0000 | 0.0001 |
| FMN pH 8 | 0.0000 | 200 | 0.0000 | 0.0000 | 0.0000 |

| | | | | | |
|---|---|---|---|---|---|
| pH 8 buffer | 0.0000 | 200 | 0.0000 | 0.0000 | 0.0000 |
| $Fe^0$ FAD pH 8 | 150.6680 | 200 | 0.0065 | 0.0038 | 0.0103 |
| | 79.8149 | 200 | 0.0035 | 0.0020 | 0.0055 |
| | 118.3708 | 200 | 0.0051 | 0.0030 | 0.0081 |
| $Fe^0$ FMN pH 8 | 89.9082 | 200 | 0.0039 | 0.0022 | 0.0061 |
| | 85.9556 | 200 | 0.0037 | 0.0021 | 0.0059 |
| | 111.4898 | 200 | 0.0048 | 0.0028 | 0.0076 |
| $Ni^0$ FAD pH 10 | 0.5160 | 100 | 0.0000 | 0.0000 | 0.0000 |
| | 1.4588 | 100 | 0.0001 | 0.0000 | 0.0001 |
| | 1.0827 | 100 | 0.0001 | 0.0000 | 0.0001 |
| FAD pH 10 | 0.0000 | 100 | 0.0000 | 0.0000 | 0.0000 |
| $N^0$i FMN pH 10 | 0.0000 | 100 | 0.0000 | 0.0000 | 0.0000 |
| | 0.0000 | 100 | 0.0000 | 0.0000 | 0.0000 |
| | 0.0000 | 100 | 0.0000 | 0.0000 | 0.0000 |
| FMN pH 10 | 0.0000 | 100 | 0.0000 | 0.0000 | 0.0000 |
| pH 10 buffer | 0.0000 | 100 | 0.0000 | 0.0000 | 0.0000 |
| $Fe^0$ FAD pH 10 | 0.0000 | 100 | 0.0000 | 0.0000 | 0.0000 |
| | 0.0000 | 100 | 0.0000 | 0.0000 | 0.0000 |
| | 0.0000 | 100 | 0.0000 | 0.0000 | 0.0000 |
| $Fe^0$ FMN pH 10 | 0.0000 | 100 | 0.0000 | 0.0000 | 0.0000 |
| | 0.0000 | 100 | 0.0000 | 0.0000 | 0.0000 |
| | 0.0000 | 100 | 0.0000 | 0.0000 | 0.0000 |
| $Ni^0$ pH 6 | 203.0344 | 200 | 0.0088 | 0.0051 | 0.0139 |
| | 251.4409 | 200 | 0.0109 | 0.0063 | 0.0172 |
| | 179.3238 | 200 | 0.0078 | 0.0045 | 0.0123 |
| $Fe^0$ pH 6 | 751.4790 | 200 | 0.0326 | 0.0188 | 0.0514 |
| | 726.9309 | 200 | 0.0315 | 0.0182 | 0.0497 |
| | 987.3112 | 200 | 0.0428 | 0.0247 | 0.0675 |
| $Ni^0$ pH 8 | 73.8920 | 200 | 0.0032 | 0.0018 | 0.0051 |
| | 7.4240 | 200 | 0.0003 | 0.0002 | 0.0005 |
| | 6.8294 | 200 | 0.0003 | 0.0002 | 0.0005 |
| $Fe^0$ pH 8 | 521.2754 | 200 | 0.0226 | 0.0130 | 0.0356 |
| | 444.2809 | 200 | 0.0193 | 0.0111 | 0.0304 |
| | 541.9534 | 200 | 0.0235 | 0.0135 | 0.0370 |
| $Ni^0$ pH 10 | 44.2085 | 200 | 0.0019 | 0.0011 | 0.0030 |
| | 24.5514 | 200 | 0.0011 | 0.0006 | 0.0017 |
| | 6.3836 | 200 | 0.0003 | 0.0002 | 0.0004 |
| $Fe^0$ pH 10 | 23.2461 | 200 | 0.0010 | 0.0006 | 0.0016 |
| | 21.7676 | 200 | 0.0009 | 0.0005 | 0.0015 |
| | 16.9285 | 200 | 0.0007 | 0.0004 | 0.0012 |

### Inductively Coupled Plasma Mass Spectrometry (ICP-MS)

**Table S24.** Dissolved Fe and Ni ions of the samples after the 2-hour reactions. The concentration was adjusted from the dilution 100/10000.

| Sample (metal + cofactor + pH + atmosphere) | Fe (mM) | Ni (mM) | | Sample (metal + pH + atmosphere) | Fe (mM) | Ni (mM) |
|---|---|---|---|---|---|---|
| $Ni^0$ FAD pH 6 $N_2$ | 0.006 | 0.702 | | pH 6 $N_2$ | 0.008 | 0.003 |
| | 0.044 | 1.115 | | $Ni^0$ pH 6 $N_2$ | 0.037 | 1184 |
| | 0.005 | 1.178 | | | 0.031 | 1103 |
| FAD pH 6 $N_2$ | 0.007 | 0.002 | | | 0.070 | 0.537 |
| $Ni^0$ FMN pH 6 $N_2$ | 0.020 | 3.147 | | | | |
| | 0.033 | 3.349 | | | | |
| | 0.015 | 2.881 | | | | |
| FMN pH 6 $N_2$ | 0.009 | 0.004 | | | | |
| $Fe^0$ FAD pH 6 $N_2$ | 0.144 | 0.003 | | $Fe^0$ pH 6 $N_2$ | 0.136 | 0.001 |
| | 0.158 | 0.002 | | | 0.129 | 0.001 |
| | 0.187 | 0.002 | | | 0.138 | 0.002 |
| $Fe^0$ FMN pH 6 $N_2$ | 1.210 | 0.002 | | | | |
| | 1.365 | 0.003 | | | | |
| | 0.916 | 0.002 | | | | |
| $Ni^0$ FAD pH 8 $N_2$ | 0.009 | 0.195 | | pH 8 $N_2$ | 0.008 | 0.002 |
| | 0.014 | 0.385 | | $Ni^0$ pH 8 $N_2$ | 0.014 | 0.184 |
| | 0.010 | 0.473 | | | 0.013 | 0.260 |
| FAD pH 8 $N_2$ | 0.014 | 0.005 | | | 0.011 | 0.202 |
| $Ni^0$ FMN pH 8 $N_2$ | 0.017 | 1.163 | | | | |
| | 0.007 | 0.445 | | | | |
| | 0.010 | 0.338 | | | | |
| FMN pH 8 $N_2$ | 0.011 | 0.003 | | | | |
| $Fe^0$ FAD pH 8 $N_2$ | 0.026 | 0.002 | | $Fe^0$ pH 8 $N_2$ | 0.068 | 0.002 |
| | 0.021 | 0.002 | | | 0.057 | 0.008 |
| | 0.037 | 0.002 | | | 0.078 | 0.003 |
| $Fe^0$ FMN pH 8 $N_2$ | 0.034 | 0.002 | | | | |
| | 0.048 | 0.002 | | | | |
| | 0.030 | 0.002 | | | | |
| $Ni^0$ FAD pH 10 $N_2$ | 0.007 | 0.025 | | pH 10 $N_2$ | 0.007 | 0.001 |
| | 0.008 | 0.011 | | $Ni^0$ pH 10 $N_2$ | 0.006 | 0.031 |
| | 0.006 | 0.022 | | | 0.005 | 0.039 |
| FAD pH 10 $N_2$ | 0.007 | 0.002 | | | 0.005 | 0.023 |

| | | | | | | |
|---|---|---|---|---|---|---|
| $Ni^0$ FMN pH 10 $N_2$ | 0.005 | 0.025 | | | | |
| | 0.006 | 0.016 | | | | |
| | 0.007 | 0.023 | | | | |
| FMN pH 10 $N_2$ | 0.007 | 0.001 | | | | |
| $Fe^0$ FAD pH 10 $N_2$ | 0.008 | 0.002 | | $Fe^0$ pH 10 $N_2$ | 0.017 | 0.002 |
| | 0.009 | 0.002 | | | 0.011 | 0.001 |
| | 0.009 | 0.003 | | | 0.013 | 0.001 |
| $Fe^0$ FMN pH 10 $N_2$ | 0.008 | 0.002 | | | | |
| | 0.007 | 0.003 | | | | |
| | 0.006 | 0.002 | | | | |
| $Ni^0$ FAD pH 6 $H_2$ | 0.051 | 0.278 | | pH 6 $H_2$ | 0.004 | 0.003 |
| | 0.043 | 0.291 | | $Ni^0$ pH 6 $H_2$ | 0.039 | 0.293 |
| | 0.028 | 0.492 | | | 0.026 | 0.273 |
| FAD pH 6 $H_2$ | 0.006 | 0.004 | | | 0.019 | 0.250 |
| $Ni^0$ FMN pH 6 $H_2$ | 0.025 | 0.651 | | | | |
| | 0.041 | 0.484 | | | | |
| | 0.111 | 0.602 | | | | |
| FMN pH 6 $H_2$ | 0.010 | 0.039 | | | | |
| $Fe^0$ FAD pH 6 $H_2$ | 0.121 | 0.008 | | $Fe^0$ pH 6 $H_2$ | 0.125 | 0.002 |
| | 0.152 | 0.005 | | | 0.104 | 0.006 |
| | 0.234 | 0.006 | | | 0.121 | 0.001 |
| $Fe^0$ FMN pH 6 $H_2$ | 0.818 | 0.007 | | | | |
| | 1.070 | 0.007 | | | | |
| | 1.310 | 0.009 | | | | |
| $Ni^0$ FAD pH 8 $H_2$ | 0.021 | 0.129 | | pH 8 $H_2$ | 0.006 | 0.003 |
| | 0.024 | 0.194 | | $Ni^0$ pH 8 $H_2$ | 0.022 | 0.013 |
| | 0.017 | 0.117 | | | 0.015 | 0.012 |
| FAD pH 8 $H_2$ | 0.025 | 0.007 | | | 0.015 | 0.014 |
| $Ni^0$ FMN pH 8 $H_2$ | 0.029 | 0.335 | | | | |
| | 0.015 | 0.225 | | | | |
| | 0.016 | 0.272 | | | | |
| FMN pH 8 $H_2$ | 0.011 | 0.005 | | | | |
| $Fe^0$ FAD pH 8 $H_2$ | 0.014 | 0.004 | | $Fe^0$ pH 8 $H_2$ | 0.051 | 0.003 |
| | 0.031 | 0.056 | | | 0.061 | 0.004 |
| | 0.015 | 0.034 | | | 0.046 | 0.002 |
| $Fe^0$ FMN pH 8 $H_2$ | 0.032 | 0.004 | | | | |
| | 0.043 | 0.004 | | | | |
| | 0.028 | 0.010 | | | | |

| | | | | | |
|---|---|---|---|---|---|
| $Ni^0$ FAD pH 10 $H_2$ | 0.078 | 0.013 | pH 10 $H_2$ | 0.024 | 0.044 |
| | 0.014 | 0.020 | $Ni^0$ pH 10 $H_2$ | 0.006 | 0.012 |
| | 0.007 | 0.010 | | 0.006 | 0.012 |
| FAD pH 10 $H_2$ | 0.009 | 0.019 | | 0.005 | 0.016 |
| $Ni^0$ FMN pH 10 $H_2$ | 0.042 | 0.011 | | | |
| | 0.019 | 0.014 | | | |
| | 0.008 | 0.027 | | | |
| FMN pH 10 $H_2$ | 0.042 | 0.005 | | | |
| $Fe^0$ FAD pH 10 $H_2$ | 0.051 | 0.027 | $Fe^0$ pH 10 $H_2$ | 0.012 | 0.003 |
| | 0.039 | 0.073 | | 0.008 | 0.002 |
| | 0.019 | 0.005 | | 0.029 | 0.001 |
| $Fe^0$ FMN pH 10 $H_2$ | 0.017 | 0.012 | | | |
| | 0.012 | 0.010 | | | |
| | 0.008 | 0.006 | | | |

### Ferrozine test

To distinguish between $Fe^{2+}$ and $Fe^{3+}$ ions, a colorimetric ferrozine assay was used. 0.01 M of ferrozine (95+%, Thermo Scientific) was prepared in 0.1 M ammonium acetate solution, and stored in the dark at 4°C. For the reducing agent, 1.4 M of hydroxylamine hydrochloride is prepared in 2 M HCl. Standards were made from a 17.9 mM $FeCl_3$ (0.01 M HCl) stock by diluting it with a 0.15 M NaCl solution. In order to get absorbance values between 0.1–1, different volumes were aliquoted from the original samples at each pH: 9 µL (pH 6), 36 µL (pH 8), and 100 µL (pH 10) samples. Each aliquot was diluted to 100 µL with water before further reagent addition, excpet pH 10 samples. To measure the total amount of Fe ions, 100 µL of the reducing agent and 100 µL of the ferrozine solution were added to the sample. Prior to analysis the samples' pH was measured and then adjusted with 10 M of ammonium acetate (pH of 9.5) or 1 M HCL to pH 5. The amount of $Fe^{2+}$ was determined from aliquots in which reducing agent was replaced with equal volume of HPLC water. The samples were left to incubate for 30 min in the dark, and then measured on a 96-well plate through UV-Vis spectroscopy, using a Gen5 Microplate Reader and Imager Software (version 3.03, Agilent Technologies) (see **Table S26.** and **Fig. S9.**). The $Fe^{3+}$ concentration was calculated by subtracting the $Fe^{2+}$ from total Fe ions. All the tests were done under a nitrogen atmosphere and in low light conditions, until the measurement.

**Table S25.** Calibration curve of the ferrozine assay to quantify $FADH_2$ and $FMNH_2$ reducing hematite and magnetite samples. $Fe^{3+}$ triplicate standards (17.9 mM $FeCl_3$ stock diluted with 0.15 M NaCl) were reduced with 1.4 M hydroxylamine hydrochloride (2 M HCl), and then added into 0.1 M ferrozine ( 0.1 M ammonium acetate) in 96-well microplate with 100 µL volume. The plates were incubate at room temperature in 30 minutes in the dark, and then their absorbance was measured from 562 nm wavelength with a microplate reader.

| **Ferrozine assay for hematite and magnetite** | | | | | |
|---|---|---|---|---|---|
| *Absorbance vs $Fe^{2+}$ concentration* | | | | *Concentration (µM)* | *Absorbance* |
| 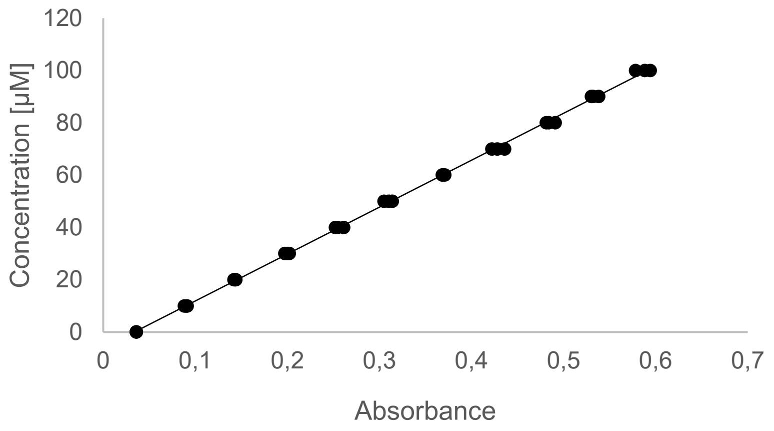 | | | | 0 | **0.036** |
| | | | | | **0.036** |
| | | | | | **0.036** |
| | | | | 10 | **0.088** |
| | | | | | **0.09** |
| | | | | | **0.091** |
| | | | | 20 | **0.143** |
| | | | | | **0.142** |
| | | | | | **0.144** |
| | | | | 30 | **0.202** |
| | | | | | **0.2** |
| | | | | | **0.197** |
| | | | | 40 | **0.252** |
| | | | | | **0.261** |
| | | | | | **0.254** |
| $\mathbf{y = 179.42x - 6.0592, R^2 = 0.9993}$ | | | | 50 | **0.314** |
| | | | | | **0.31** |
| | | | | | **0.305** |
| 60 | **0.369** | 70 | **0.428** | 80 | **0.484** |
| | **0.368** | | **0.436** | | **0.481** |
| | **0.371** | | **0.422** | | **0.491** |
| 90 | **0.532** | 100 | **0.594** | | |
| | **0.538** | | **0.578** | | |
| | **0.53** | | **0.588** | | |

**Table S26.** Calibration curve of the ferrozine assay to quantify $Fe^{2+}$. $Fe^{3+}$ triplicate standards (17.9 mM $FeCl_3$ stock diluted with 0.15 M NaCl) were reduced with 1.4 M hydroxylamine hydrochloride (2 M HCl), and then added into 0.1 M ferrozine ( 0.1 M ammonium acetate) in 96-well microplate with 300 µL volume. The plates were incubate at room temperature in 30 minutes in the dark, and then their absorbance was measured from 562 nm wavelength with a microplate reader.

**Ferrozine assay for $FeCl_3$**

*Absorbance vs **$Fe^{2+}$ concentration***

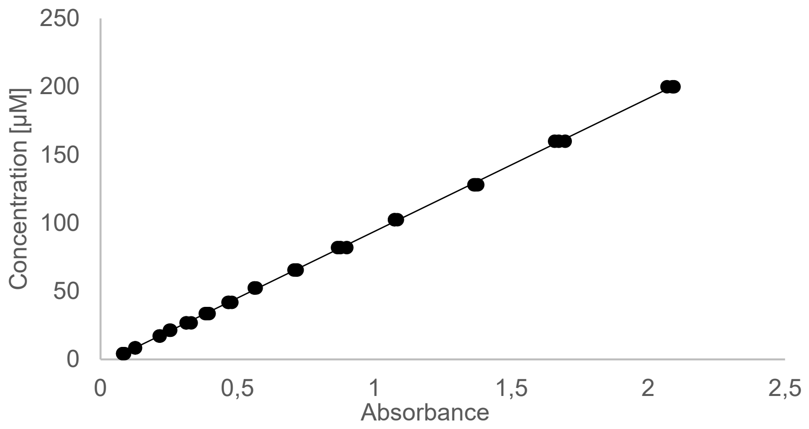


$$y = 97.584x - 3.8091, R^2 = 0.9997$$

| *Concentration (µM)* | *Absorbance* |
|---|---|
| **4.29** | **0.081** |
| | **0.088** |
| | **0.081** |
| **8.589** | **0.127** |
| | **0.127** |
| | **0.125** |
| **17.179** | **0.216** |
| | **0.217** |
| | **0.214** |
| **21.47** | **0.252** |
| | **0.256** |
| | **0.254** |
| **33.554** | **0.397** |
| | **0.388** |
| | **0.382** |
| **41.943** | **0.469** |
| | **0.479** |
| | **0.465** |

| | | | | | |
|---|---|---|---|---|---|
| **52.428** | **0.563** | **65.536** | **0.706** | **81.92** | **0.876** |
| | **0.568** | | **0.718** | | **0.899** |
| | **0.561** | | **0.714** | | **0.865** |
| **102.4** | **1.074** | **128** | **1.363** | **160** | **1.673** |
| | **1.084** | | **1.378** | | **1.697** |
| | **1.073** | | **1.369** | | **1.658** |
| **200** | **2.069** | | | | |
| | **2.088** | | | | |
| | **2.094** | | | | |

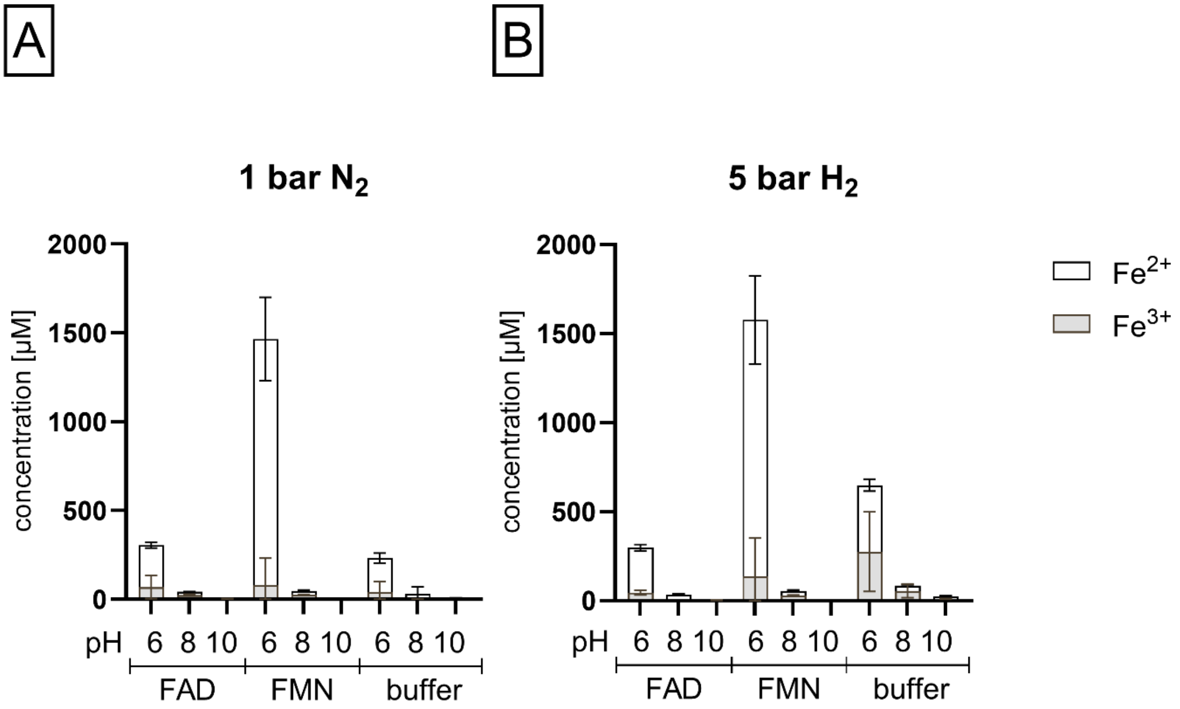


**Fig. S9.** $Fe^{2+}$ and $Fe^{3+}$ formed during standard reaction conditions under $H_2$ (A) or $N_2$ (B) were determined through ferrozine colometric assays as described in the method above. The samples were quantified based on **Table S30.**

**Table S27.** Table of samples' $Fe^{2+}$ and $Fe^{3+}$ ions as determined per a ferrozine assay and following calculations as described in the methods above. The molarity was corrected after calculating the concentration from the absorbance, as pH 6 samples were 6/100 diluted and pH 8 samples were 36/100 diluted. The samples were quantified based on **Table S30.**

| **Sample (cofactor+pH+atmosphere+ion)** | | **Absorbance (652 nm)** | **Ion concentration (mM)** |
|---|---|---|---|
| FAD pH 6 $H_2$ | $Fe^{2+}$ | 0.193 | 250.4 |
| | | 0.206 | 271.6 |
| | | 0.184 | 235.8 |
| | $Fe^{2+}$/ $Fe^{3+}$ | 0.221 | 295.9 |
| | | 0.243 | 331.7 |
| | | 0.204 | 268.3 |
| | $Fe^{3+}$ | N/A | 45.5 |
| | | N/A | 60.2 |
| | | N/A | 32.5 |
| FMN pH 6 $H_2$ | $Fe^{2+}$ | 1.099 | 1723.9 |
| | | 0.851 | 1320.6 |
| | | 0.822 | 1273.4 |
| | $Fe^{2+}$/ $Fe^{3+}$ | 1.08 | 1693.0 |
| | | 1.085 | 1701.2 |
| | | 0.861 | 1336.8 |
| | $Fe^{3+}$ | N/A | -30.9 |
| | | N/A | 380.6 |
| | | N/A | 63.4 |
| FAD pH 6 $N_2$ | $Fe^{2+}$ | 0.195 | 253.7 |
| | | 0.178 | 226.0 |
| | | 0.178 | 226.0 |
| | $Fe^{2+}$/ $Fe^{3+}$ | 0.281 | 393.5 |
| | | 0.185 | 237.4 |
| | | 0.214 | 284.6 |
| | $Fe^{3+}$ | N/A | 139.9 |
| | | N/A | 11.4 |
| | | N/A | 58.6 |
| FMN pH 6 $N_2$ | $Fe^{2+}$ | 0.737 | 1135.2 |
| | | 1.025 | 1603.6 |
| | | 0.91 | 1416.5 |
| | $Fe^{2+}$/ $Fe^{3+}$ | 0.791 | 1223.0 |
| | | 1.166 | 1832.9 |
| | | 0.864 | 1341.7 |
| | $Fe^{3+}$ | N/A | 87.8 |
| | | N/A | 229.3 |
| | | N/A | -74.8 |
| FAD pH 8 $H_2$ | $Fe^{2+}$ | 0.163 | 33.6 |
| | | 0.168 | 35.0 |

| | | | |
|---|---|---|---|
| | | 0.191 | 41.2 |
| | $Fe^{2+}$/ $Fe^{3+}$ | 0.163 | 33.6 |
| | | 0.161 | 33.1 |
| | | 0.179 | 37.9 |
| | $Fe^{3+}$ | N/A | 0.0 |
| | | N/A | -1.9 |
| | | N/A | -3.3 |
| FMN pH 8 $H_2$ | $Fe^{2+}$ | 0.157 | 32.0 |
| | | 0.116 | 20.9 |
| | | 0.129 | 24.4 |
| | $Fe^{2+}$/ $Fe^{3+}$ | 0.266 | 61.5 |
| | | 0.236 | 53.4 |
| | | 0.226 | 50.7 |
| | $Fe^{3+}$ | N/A | 29.5 |
| | | N/A | 32.5 |
| | | N/A | 26.3 |
| FAD pH 8 $N_2$ | $Fe^{2+}$ | 0.104 | 17.6 |
| | | 0.1 | 16.5 |
| | | 0.115 | 20.6 |
| | $Fe^{2+}$/ $Fe^{3+}$ | 0.212 | 46.9 |
| | | 0.174 | 36.6 |
| | | 0.198 | 43.1 |
| | $Fe^{3+}$ | N/A | 29.3 |
| | | N/A | 20.1 |
| | | N/A | 22.5 |
| FMN pH 8 $N_2$ | $Fe^{2+}$ | 0.094 | 14.9 |
| | | 0.122 | 22.5 |
| | | 0.128 | 24.1 |
| | $Fe^{2+}$/ $Fe^{3+}$ | 0.195 | 42.3 |
| | | 0.219 | 48.8 |
| | | 0.228 | 51.2 |
| | $Fe^{3+}$ | N/A | 27.4 |
| | | N/A | 26.3 |
| | | N/A | 27.1 |
| FAD pH 10 $H_2$ | $Fe^{2+}$ | 0.043 | 0.4 |
| | | 0.046 | 0.7 |
| | | 0.046 | 0.7 |
| | $Fe^{2+}$/ $Fe^{3+}$ | 0.077 | 3.7 |
| | | 0.093 | 5.3 |
| | | 0.063 | 2.3 |
| | $Fe^{3+}$ | N/A | 3.3 |
| | | N/A | 4.6 |
| | | N/A | 1.7 |

| | | | |
|---|---|---|---|
| FMN pH 10 $H_2$ | $Fe^{2+}$ | 0.05 | 1.1 |
| | | 0.05 | 1.1 |
| | | 0.05 | 1.1 |
| | $Fe^{2+}$/ $Fe^{3+}$ | 0.066 | 2.6 |
| | | 0.06 | 2.0 |
| | | 0.061 | 2.1 |
| | $Fe^{3+}$ | N/A | 1.6 |
| | | N/A | 1.0 |
| | | N/A | 1.1 |
| FAD pH 10 $N_2$ | $Fe^{2+}$ | 0.043 | 0.4 |
| | | 0.043 | 0.4 |
| | | 0.045 | 0.6 |
| | $Fe^{2+}$/ $Fe^{3+}$ | 0.058 | 1.9 |
| | | 0.06 | 2.0 |
| | | 0.087 | 4.7 |
| | $Fe^{3+}$ | N/A | 1.5 |
| | | N/A | 1.7 |
| | | N/A | 4.1 |
| FMN pH 10 $N_2$ | $Fe^{2+}$ | 0.045 | 0.6 |
| | | 0.045 | 0.6 |
| | | 0.045 | 0.6 |
| | $Fe^{2+}$/ $Fe^{3+}$ | 0.054 | 1.5 |
| | | 0.059 | 1.9 |
| | | 0.054 | 1.5 |
| | $Fe^{3+}$ | N/A | 0.9 |
| | | N/A | 1.4 |
| | | N/A | 0.9 |
| pH 6 $H_2$ | $Fe^{2+}$ | 0.245 | 334.983 |
| | | 0.278 | 388.6542 |
| | | 0.283 | 396.7862 |
| | $Fe^{2+}$/ $Fe^{3+}$ | 0.384 | 561.0526 |
| | | 0.328 | 469.9742 |
| | | 0.603 | 917.2342 |
| | $Fe^{3+}$ | N/A | 162.5846 |
| | | N/A | 17.835 |
| | | N/A | 456.963 |
| pH 8 $H_2$ | $Fe^{2+}$ | 0.133 | 25.47103 |
| | | 0.171 | 35.77157 |
| | | 0.133 | 25.47103 |
| | $Fe^{2+}$/ $Fe^{3+}$ | 0.19 | 40.92183 |
| | | 0.401 | 98.1169 |
| | | 0.466 | 115.7362 |
| | $Fe^{3+}$ | N/A | 4.869967 |

| | | | |
|---|---|---|---|
| | | N/A | 51.7645 |
| | | N/A | 79.68437 |
| pH 10 $H_2$ | $Fe^{2+}$ | 0.129 | 8.779236 |
| | | 0.199 | 15.61012 |
| | | 0.224 | 18.04972 |
| | $Fe^{2+}$/ $Fe^{3+}$ | 0.229 | 18.53764 |
| | | 0.33 | 28.39362 |
| | | 0.358 | 31.12597 |
| | $Fe^{3+}$ | N/A | 5.9493 |
| | | N/A | 8.974404 |
| | | N/A | 9.267156 |
| pH 6 $N_2$ | $Fe^{2+}$ | 0.155 | 188.607 |
| | | 0.175 | 221.135 |
| | | 0.14 | 164.211 |
| | $Fe^{2+}$/ $Fe^{3+}$ | 0.139 | 162.5846 |
| | | 0.211 | 279.6854 |
| | | 0.195 | 253.663 |
| | $Fe^{3+}$ | N/A | -89.5074 |
| | | N/A | -4.9346 |
| | | N/A | 25.967 |
| pH 8 $N_2$ | $Fe^{2+}$ | 0.076 | 10.02023 |
| | | 0.057 | 4.869967 |
| | | 0.325 | 77.51583 |
| | $Fe^{2+}$/ $Fe^{3+}$ | 0.076 | 10.02023 |
| | | 0.063 | 6.496367 |
| | | 0.314 | 74.5341 |
| | $Fe^{3+}$ | N/A | -10.5808 |
| | | N/A | -8.95443 |
| | | N/A | -13.5626 |
| pH 10 $N_2$ | $Fe^{2+}$ | 0.147 | 10.53575 |
| | | 0.132 | 9.071988 |
| | | 0.133 | 9.169572 |
| | $Fe^{2+}$/ $Fe^{3+}$ | 0.143 | 10.14541 |
| | | 0.126 | 8.486484 |
| | | 0.128 | 8.681652 |
| | $Fe^{3+}$ | N/A | -4.19944 |
| | | N/A | -4.3946 |
| | | N/A | -4.29702 |

**$FADH_2$ and $FMNH_2$ reducing free $Fe^{3+}$.** FAD and FMN (4 mM) in HPLC water were reduced under $H_2$ with $Ni^0$ for 2 hours. The samples were centrifuged (4°C, 13 000 rpm, 20 min), and the supernatant recovered from the reaction tube. Complete reduction was confirmed with UV-visible spectroscopy. All preparation was done under gloveboxes $N_2$ atmosphere before the measurement. The samples were diluted to 80 µM.

From 17.9 mM stock solution of $FeCl_3$ (98%; Grüssing GmbH) a 160 µM dilution was made in HPLC-grade water. Ferrozine stock solution (10mM) was prepared in 0.1 M ammonium acetate and then diluted to 1 mM with HPLC water before use. 100 µL of the $FADH_2$ or $FMNH_2$, 100 µL of the 160 µM $FeCl_3$, and 100 µL of the 1 mM ferrozine solution were pipetted into a reaction tube and vortexed. The mixture was transferred onto a 96-well plate and measured with microwellplate reader from the 562 nm. The absorbance was converted into a concentration based on the

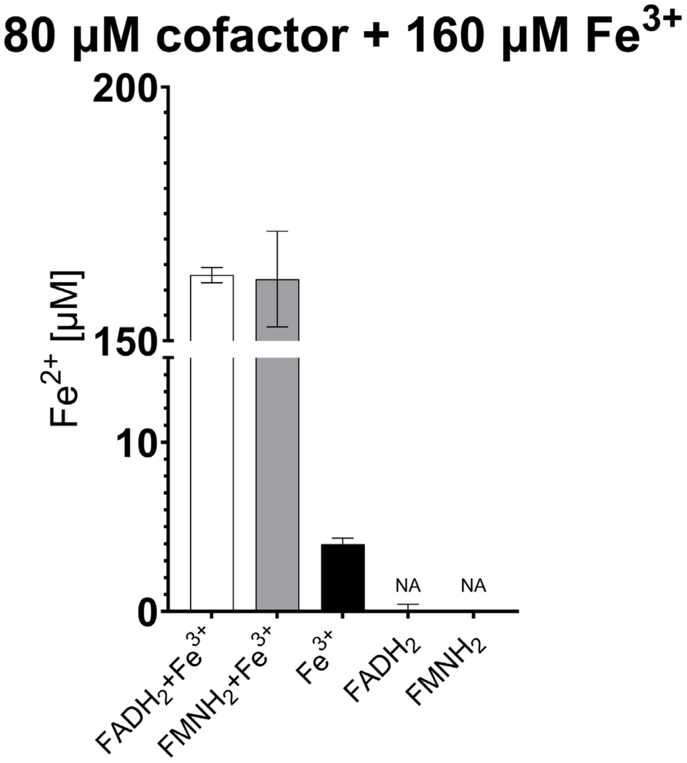


**Fig. S10.** Ferrozine colometric assay graphs showing $Fe^{2+}$ concentration right after mixing 80 µM of each reduced flavin and double the amount of $FeCl_3$. The samples were quantified based on the standard curve in **Table S25.**

**Table S28.** $Fe^{2+}$ concentration in the sample, after adding 80 µM of reduced flavin to 160 µM $FeCl_3$, and controls without.

| | $FADH_2$+$Fe^{3+}$ | $FMNH_2$+$Fe^{3+}$ | $Fe^{3+}$ | $FADH_2$ | $FMNH_2$ |
|---|---|---|---|---|---|
| $Fe^{2+}$ | 161.30 | 168.62 | 3.80 | -0.30 | -0.01 |
| | 163.35 | 166.57 | 3.80 | 0.28 | -0.01 |
| | 164.22 | 151.34 | 4.38 | 0.28 | -0.01 |

**$FADH_2$ and $FMNH_2$ reducing minerals.**

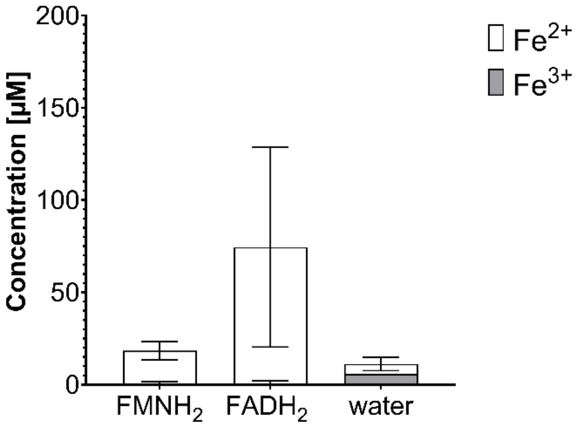


**Fig. S11.** Ferrozine colometric assay was tested with reduced flavin controls, showing the baseline amount of $Fe^{2+}$ and $Fe^{3+}$ after two hour stirring at room temperature (t = 2). The samples were quantified based on **Table S25.**

**Table S29.** Table of sample mixtures of reduced flavins with hematite or magnetite powders, tested with ferrozine assay at t=0 and t=2 to determined the reduction and dissolved $Fe^{2+}$ formed as a reaction product. (*)The negative value of $Fe^{3+}$ indicates an issue with the measurement, so both values were not included in the graph.

| # | $FMNH_2$ (mM) | FMN (mM) | $FADH_2$ (mM) | Hematite (mM) | Magnetite (mM) | $Fe^{2+}$ (µM) | | $Fe^{3+}$ (µM) | | pH after reaction |
|---|---|---|---|---|---|---|---|---|---|---|
| | | | | | | t = 0 | t = 2 | t = 0 | t = 2 | |
| 1 | 4 | | | 18 | | 357,05 | 410,63 | 0,00 | 0,00 | 7.42 |
| 2 | 4 | | | 18 | | 346,34 | 404,38 | 0,00 | 0,00 | 7.34 |
| 3 | 4 | | | 18 | | 340,98 | 399,91 | 0,00 | 0,00 | 7.27 |
| 4 | | 4 | | 18 | | 98,13 | 56,16 | 6,88 | 6,13 | 6.22 |
| 5 | | 4 | | 18 | | 82,95 | 54,38 | 9,56 | 4,79 | 6.25 |
| 6 | | 4 | | 18 | | 90,98 | 59,73 | 5,69 | 3,60 | 6.27 |
| 7 | | | 4 | 18 | | 307,95 | 326,70 | 0,00 | 0,00 | 7.2 |
| 8 | | | 4 | 18 | | 298,13 | 332,95 | 0,00 | 0,00 | 7.52 |
| 9 | | | 4 | 18 | | 299,02 | 336,52 | 0,00 | 0,00 | 8.02 |
| 10 | | 4 | | | 18 | 48,13 | 38,30 | 4,79 | 5,23 | 6.35 |
| 11 | | 4 | | | 18 | 57,95 | 74,02 | 4,34 | 5,98 | 6.2 |
| 12 | | 4 | | | 18 | 52,59 | 71,34 | 3,45 | 5,54 | 6.15 |
| 13 | 4 | | | | 18 | 279,38 | 329,38 | 0,00 | 0,00 | 6.47 |
| 14 | 4 | | | | 18 | 290,98 | 353,48 | 0,00 | 0,00 | 6.58 |
| 15 | 4 | | | | 18 | 245,45 | 328,48 | 0,00 | 0,00 | 6.45 |
| 16 | | | 4 | | 18 | 169,55 | 277,59 | 6,31 | 0,00 | 6.83 |
| 17 | | | 4 | | 18 | 150,80 | 274,91 | 10,47 | 0,00 | 6.87 |
| 18 | | | 4 | | 18 | 141,88 | 299,02 | 12,11 | 0,00 | 6.83 |
| 19 | | | | 18 | | 6,16 | *236,52** | 6,11 | *-5,43** | 4.1 |
| 20 | | | | 18 | | 7,95 | 7,95 | 5,37 | 10,58 | 8.8 |
| 21 | | | | 18 | | 8,84 | 4,38 | 7,60 | 5,82 | 8.84 |
| 22 | | | | | 18 | 7,05 | 14,20 | 4,18 | 4,33 | 7.77 |
| 23 | | | | | 18 | 5,27 | 5,27 | 5,96 | 5,96 | 7.2 |
| 24 | | | | | 18 | 3,48 | 8,84 | 5,67 | 6,56 | 7.78 |
| 25 | 4 | | | | | n/a | 18,66 | n/a | 0,00 | 6.2 |
| 26 | 4 | | | | | n/a | 22,23 | n/a | 0,00 | 6.27 |
| 27 | 4 | | | | | n/a | 12,41 | n/a | 1,95 | 6.22 |
| 28 | | | 4 | | | n/a | 77,59 | n/a | 2,41 | 6.52 |
| 29 | | | 4 | | | n/a | 17,77 | n/a | 0,00 | 6.54 |
| 30 | | | 4 | | | n/a | 125,80 | n/a | 0,00 | 6.55 |
| 31 | | | | | | n/a | 9,73 | n/a | 5,67 | 8.89 |
| 32 | | | | | | n/a | 3,48 | n/a | 5,67 | 8.9 |
| 33 | | | | | | n/a | 3,48 | n/a | 5,67 | 8.28 |

**$FMNH_2$ recycling with magnetite.**

**Table S30.** Fe ion concentrations of FMN one pot reaction and controls. (*)One of the FMN+$Fe_3O_4$ samples had an iron contamination from a dirty stirring bar so its measurement was excluded.

| | | $Fe^{2+}$ | | | $Fe^{3+}$ | | |
|---|---|---|---|---|---|---|---|
| **FMN +$Ni^0$+$Fe_3O_4$** | $H_2$ | 7.57672 | 11.70338 | 7.75614 | 1.949763 | 2.561686 | 1.985733 |
| **FMN+$Fe_3O_4$** | | 4.34716 | 4.52658 | *310.7965** | 1.08691 | 0.90749 | *94.83246** |
| **$Ni^0$+$Fe_3O_4$** | | 3.45006 | 2.9118 | 4.706 | 1.337839 | 1.014539 | 1.15885 |
| **$Fe_3O_4$** | | 1.65586 | 1.29702 | 1.83528 | 0.547357 | 0.690807 | 0.798718 |
| **FMN+$Ni^0$+$Fe_3O_4$** | $N_2$ | 19.39184 | 18.56668 | 16.37767 | 85.98326 | 87.2392 | 89.21282 |
| **FMN+$Fe_3O_4$** | | 1.195251 | 1.159281 | 1.267191 | 5.96194 | 5.78252 | 6.32078 |
| **$Ni^0$+$Fe_3O_4$** | | 5.116429 | 7.236083 | 4.576446 | 26.59524 | 30.7219 | 22.82742 |
| **$Fe_3O_4$** | | 0.870658 | 0.942598 | 1.014539 | 2.19412 | 2.55296 | 2.9118 |

**Table S31.** Volume-adjusted amounts of magnetite in each redox cycle displayed in **Table S32**.

| Cycle | Magnetite [mg] | Starting volume [mL] | Aliquot [µL] | Remaining Volume [mL] |
|---|---|---|---|---|
| 1 | 3.5 | 1.7 | 200 | 1.5 |
| 2 | 3.1 | 1.5 | 200 | 1.3 |
| 3 | 2.7 | 1.3 | 300 | 1 |
| 4 | 1.7 | 0.8 | 200 | 0.6 |
| 5 | 1.3 | 0.6 | 200 | 0.4 |
| 6 | 0.9 | 0.4 | 200 | 0.2 |

**Table S32.** $Fe^{2+}$, $Fe^{3+}$, pH, and $FMNH_2$ concentrations of samples after $Ni^0$ reduction and reactions with $Fe_3O_4$. Fe amounts were determined through a ferrozine assay and falvins amounts through UV-Vis spectroscopy as described by the corresponding methods. (*) Two samples were excluded due to issues handling the caps and mainting an air-tight environment.

| | **$FMNH_2$ + magnetite** | | | | | | | | | | | |
|---|---|---|---|---|---|---|---|---|---|---|---|---|
| **Cycle** | $[Fe^{2+}]$ | | | $[Fe^{3+}]$ | | | pH | | | $[FMNH_2]$ | | |
| **$Ni^0$ 1** | n/a | n/a | n/a | n/a | n/a | n/a | 5.92 | 5.92 | 5.92 | 0.85 | 0.85 | 0.85 |
| **$Fe_3O_4$ 1** | 1.09 | 1.07 | 1.02 | -0.01 | -0.11 | -0.06 | 5.95 | 5.88 | 5.87 | 0.44 | 0.42 | 0.44 |
| **$Fe_3O_4$ 2** | 1.21 | 1.28 | 1.26 | -0.10 | -0.03 | -0.01 | 5.9 | 5.91 | 5.91 | 0.25 | 0.17 | 0.19 |
| **$Fe_3O_4$ 3** | 1.25 | 1.18 | 1.31 | -0.11 | -0.03 | -0.03 | 5.9 | 5.9 | 5.9 | 0.04 | 0.00 | -0.01 |
| **$Ni^0$ 2** | 0.36 | 0.24 | 0.56 | -0.01 | 0.00 | -0.04 | 5.89 | 5.89 | 5.85 | 0.93 | 0.98 | 0.93 |
| **$Fe_3O_4$ 4** | 0.99 | 1.03 | 0.89 | n/a | n/a | n/a | 5.89 | 5.89 | 5.89 | 0.52 | 0.52 | 0.55 |
| **$Fe_3O_4$ 5** | 1.44 | 1.55 | 1.36 | n/a | n/a | n/a | 5.85 | 5.89 | 5.88 | 0.22 | 0.24 | 0.27 |
| **$Fe_3O_4$ 6** | 1.49 | 1.54 | 1.60 | -0.02 | -0.09 | -0.15 | 5.88 | 5.85 | 5.87 | 0.06 | 0.06 | 0.10 |
| | **FMN + magnetite** | | | | | | | | | | | |
| | $[Fe^{2+}]$ | | | $[Fe^{3+}]$ | | | pH | | | | | |
| **$Ni^0$ 1** | n/a | n/a | n/a | n/a | n/a | n/a | 6 | 6 | 6 | | | |
| **$Fe_3O_4$ 1** | 0.01 | 0.01 | 0.01 | 0.00 | 0.00 | 0.00 | 5.83 | 5.82 | 5.82 | | | |
| **$Fe_3O_4$ 2** | 0.01 | 0.01 | 0.01 | 0.00 | 0.00 | 0.00 | 5.8 | 5.82 | 5.81 | | | |
| **$Fe_3O_4$ 3** | 0.01 | 0.01 | 0.01 | 0.00 | 0.00 | 0.00 | 5.8 | 5.8 | 5.8 | | | |

| | | | | | | | | | | | | |
|---|---|---|---|---|---|---|---|---|---|---|---|---|
| **$Ni^0$ 2** | 0.02 | 0.01 | 0.01 | -0.01 | 0.00 | 0.00 | 5.81 | 5.8 | 5.82 | | | |
| **$Fe_3O_4$ 4** | 0.02 | 0.02 | 0.02 | 0.00 | 0.00 | 0.00 | 5.81 | 5.81 | 5.81 | | | |
| **$Fe_3O_4$ 5** | 0.02 | 0.02 | 0.04 | 0.00 | 0.00 | 0.00 | 5.8 | 5.8 | 5.8 | | | |
| **$Fe_3O_4$ 6** | 0.02 | 0.02 | 0.02 | 0.00 | 0.00 | 0.00 | 5.76 | 5.76 | 5.74 | | | |
| | **magnetite** | | | | | | | | | | | |
| | $[Fe^{2+}]$ | | | $[Fe^{3+}]$ | | | pH | | | | | |
| **$Ni^0$ 1** | n/a | n/a | n/a | n/a | n/a | n/a | 6 | 6 | 6 | | | |
| **$Fe_3O_4$ 1** | 0.00 | 0.00 | 0.00 | 0.00 | 0.00 | 0.00 | 5.9 | 5.82 | 5.9 | | | |
| **$Fe_3O_4$ 2** | 0.00 | 0.00 | 0.00 | 0.00 | 0.00 | 0.00 | 5.9 | 5.9 | 5.9 | | | |
| **$Fe_3O_4$ 3** | 0.00 | 0.00 | 0.00 | 0.00 | 0.00 | 0.00 | 5.9 | 5.9 | 5.85 | | | |
| **$Ni^0$ 2** | 0.00 | 0.01 | 0.00 | 0.00 | 0.00 | 0.00 | 5.8 | 5.8 | 5.8 | | | |
| **$Fe_3O_4$ 4** | 0.00 | 0.00 | 0.00 | 0.00 | 0.00 | 0.00 | 5.8 | 5.8 | 5.81 | | | |
| **$Fe_3O_4$ 5** | 0.00 | 0.00 | 0.00 | 0.00 | 0.00 | 0.00 | 5.8 | 5.8 | 5.78 | | | |
| **$Fe_3O_4$ 6** | 0.00 | 0.00 | 0.00 | 0.00 | 0.00 | 0.00 | 5.76 | 5.73 | 5.73 | | | |
| | **$FMNH_2$** | | | | | | | | | | | |
| | $[Fe^{2+}]$ | | | $[Fe^{3+}]$ | | | pH | | | $[FMNH_2]$ | | |
| **$Ni^0$ 1** | n/a | n/a | n/a | n/a | n/a | n/a | 5.92 | 5.92 | 5.92 | 0.85 | 0.85 | 0.85 |
| **$Fe_3O_4$ 1** | n/a | n/a | n/a | n/a | n/a | n/a | 5.94 | 5.86 | 5.9 | 0.81 | 0.72 | 0.79 |
| **$Fe_3O_4$ 2** | n/a | n/a | n/a | n/a | n/a | n/a | 5.9 | 5.9 | 5.9 | 0.60* | 0.67 | 0.79 |
| **$Fe_3O_4$ 3** | n/a | n/a | n/a | n/a | n/a | n/a | 5.85 | 5.85 | 5.87 | 0.82 | 0.74 | 0.55* |
| **$Ni^0$ 2** | n/a | n/a | n/a | n/a | n/a | n/a | 5.81 | 5.81 | 5.81 | 0.94 | 0.91 | 0.96 |
| **$Fe_3O_4$ 4** | n/a | n/a | n/a | n/a | n/a | n/a | 5.81 | 5.81 | 5.81 | 0.65 | 0.93 | 0.96 |
| **$Fe_3O_4$ 5** | n/a | n/a | n/a | n/a | n/a | n/a | 5.78 | 5.78 | 5.78 | 0.98 | 0.83 | 0.97 |
| **$Fe_3O_4$ 6** | 0.02 | 0.02 | 0.02 | -0.000413 | -0.0007 | 0.0004478 | 5.77 | 5.76 | 5.76 | 0.97 | 0.90 | 0.96 |
| | **buffer pH 6** | | | | | | | | | | | |
| | $[Fe^{2+}]$ | | | $[Fe^{3+}]$ | | | pH | | | | | |
| **$Ni^0$ 1** | n/a | n/a | n/a | n/a | n/a | n/a | 6 | 6 | 6 | | | |
| **$Fe_3O_4$ 1** | n/a | n/a | n/a | n/a | n/a | n/a | 5.94 | 5.86 | 5.8 | | | |
| **$Fe_3O_4$ 2** | n/a | n/a | n/a | n/a | n/a | n/a | 5.9 | 5.9 | 5.9 | | | |
| **$Fe_3O_4$ 3** | n/a | n/a | n/a | n/a | n/a | n/a | 5.8 | 5.8 | 5.8 | | | |
| **$Ni^0$ 2** | n/a | n/a | n/a | n/a | n/a | n/a | 5.89 | 5.85 | 5.84 | | | |
| **$Fe_3O_4$ 4** | n/a | n/a | n/a | n/a | n/a | n/a | 5.8 | 5.8 | 5.8 | | | |
| **$Fe_3O_4$ 5** | n/a | n/a | n/a | n/a | n/a | n/a | 5.79 | 5.79 | 5.79 | | | |
| **$Fe_3O_4$ 6** | 0.00 | 0.00 | 0.00 | 0.0007342 | 0.0003036 | 0.0003754 | 5.79 | 5.76 | 5.75 | | | |

### Absorbance loss and precipitation

**Effect of metal ions in falvin absorbance.** To test the effects of dissolved metals in the absorbance of flavins, several tests were carried out with equimolar amounts of metal ions to oxidized or reduced flavins (4 mM). Stock solutions were prepared in a $N_2$-filled glovebox at 8 mM, from $NiCl_2{\cdot}6H_2O$ (97%; Thermo Scientific), $FeCl_2{\cdot}H_2O$ (Grüssing GmbH), $FeCl_3$ (98%; Grüssing GmbH), and FMN, dissolved with 0.133 M phosphate buffer at pH 6. $FMNH_2$ and $FADH_2$ solutions were also prepared to 8 mM, starting from 16 mM solutions of the oxidized flavins, mixed with equimolar amounts of sodium dithionite. Samples were prepared in 1.5 mL Reaction tube tubes by mixing 0.5 mL of a metal stock with the same volume of a flavin stock. Controls were prepared by mixing either stock with the respective buffer solution. In the case of Riboflavin, the same procedure was followed with lower concentrations of metal and flavin, to a final concentration of 0.1 mM of each, and with 0.133 M phosphate buffer and 0.1 M carbonate buffer at pHs 8 and 10 respectively. The solutions were analysed accordingly to the UV-Vis spectroscopy and quantification methods.

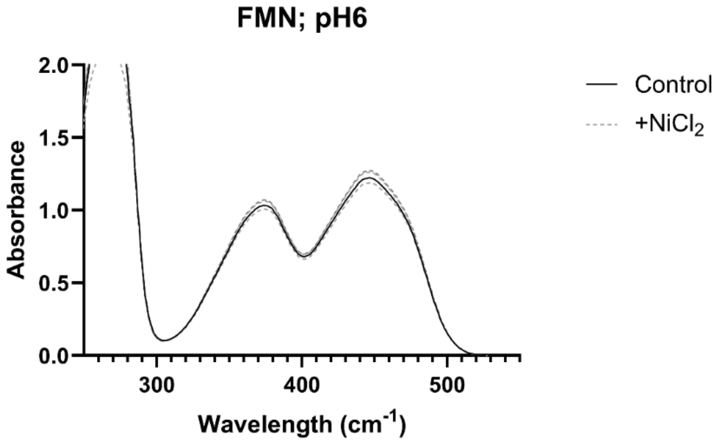


**Fig S12.** UV-Vis spectrum (200–600 nm) of dissolved samples ($\mathbf{d}_{\mathbf{FMN}}$) of originally 4 mM of FMN in 0.133M phosphate buffer at pH6, with (dashed; triplicates) and without (straight; single) 4 mM of $NiCl_2$. A white pellet was observed during sample preparation.

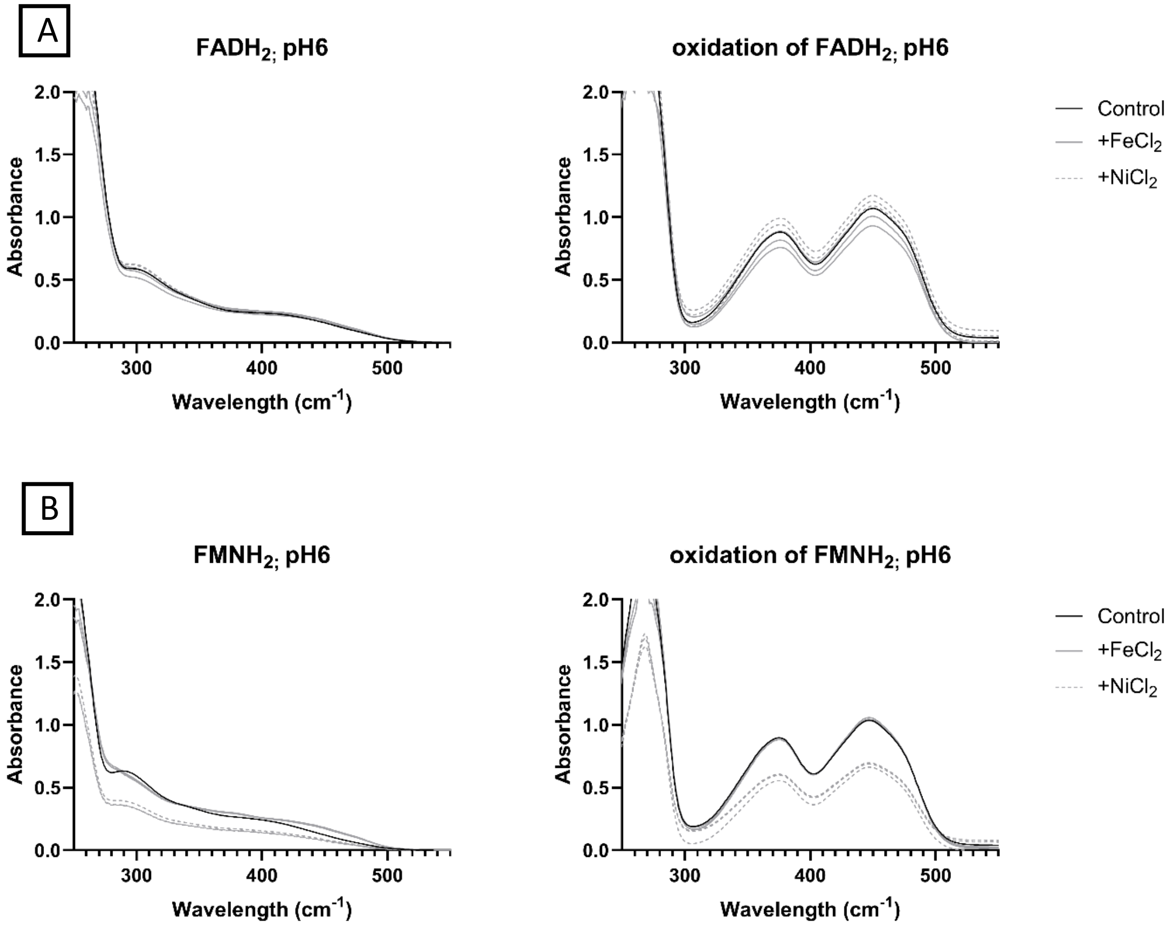


**Fig. S13.** UV-Vis spectra (200–600 nm) of dissolved samples ($\mathbf{d_{FAD;FMN}}$) that were originally 4 mM of $FADH_2$ (A) or $FMNH_2$ (B). Samples were prepared in 0.133M phosphate buffer at pH6, with equimolar amounts of sodium dithionite to the respective cofactor, then mixed with 4 mM $NiCl_2$ (dashed; triplicates), $FeCl_2$ (gray; triplicates), or no metal (black; single). The dissolved samples were analysed in air-tight cuvettes (left) and afterwards were opened and fully oxidized (right). A brown pellet was formed during sample preparation of $FMNH_2$ with $NiCl_2$.

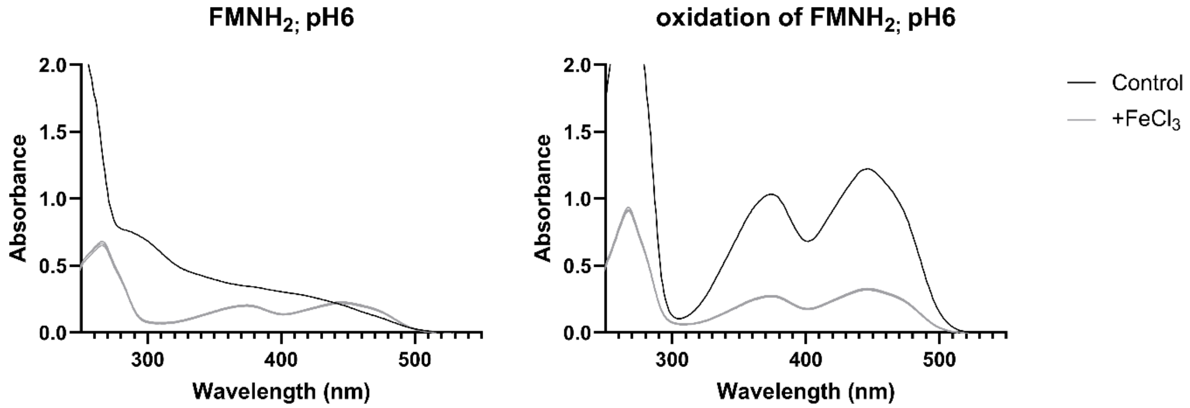


**Fig. S14.** UV-Vis spectra (200–600 nm) of dissolved samples ($\mathbf{d_{FMN}}$) that were originally 4 mM of $FMNH_2$. Samples were prepared in 0.133M phosphate buffer at pH6, with equimolar amounts of sodium dithionite to FMN, then mixed with 4 mM $FeCl_3$ (gray; triplicates), or no metal (black; single). The dissolved samples were analysed in air-tight cuvettes (left) and afterwards were opened and fully oxidized (right). A black pellet was formed during sample preparation of $FMNH_2$ with $FeCl_3$, most likely as a result of reduction of $Fe^{3+}$ species to $Fe^{2+}$, resulting in oxidation of $FMNH_2$, as seen in the left graph.

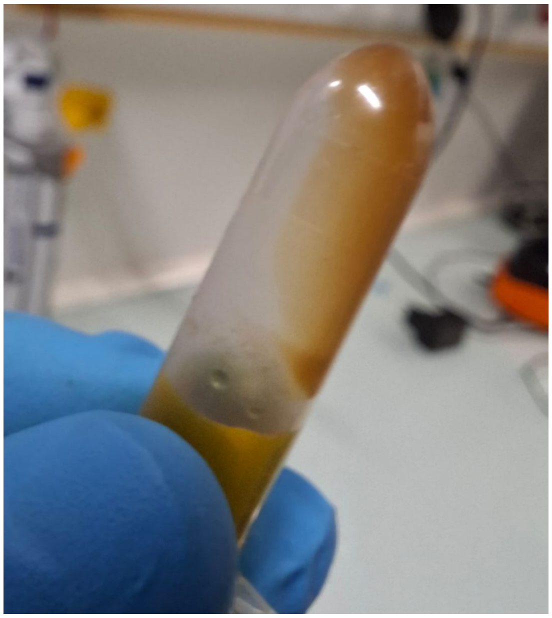

**Fig. S15.** A mixture of 4mM of $NiCl_2$ and $FMNH_2$ accumulated a red-brown pellet after centrifugation (4°C, 13 000 rpm, 20min).

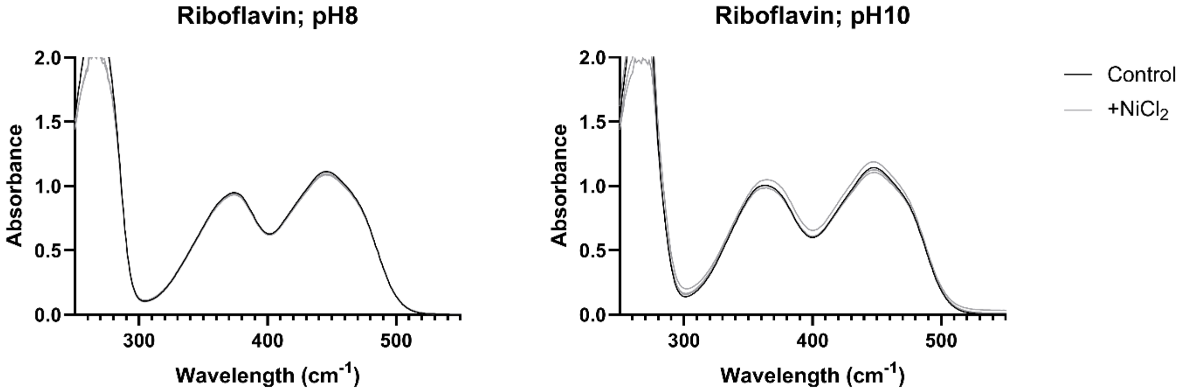


**Fig. S16.** UV-Vis spectra (200–600 nm) of 0.1 mM samples of riboflavin. Samples were prepared in 0.133M phosphate buffer at pH8 (left) or 0.1M carbonate buffer at pH10 (right), and then mixed with 0.1 mM of $NiCl_2$ (gray; triplicates), or no metal (black; single).

**Micro-X-ray fluoroscence (µXRF).** The sample preparation followed the same protocol as XRD. The µXRF measurements were performed using a M4 TORNADO (Bruker) with a Rh X-ray tube, and operated at 40 kV and 600 µA under vacuum. Elemental maps were acquired over an area of 1.0 × 0.6 mm with a step size of 20 µm and a beam/spot size of approximately 25 µm (see **Table S33.** Characterization of the pellet after a 2 h standard reaction under 5 bars of H2 or Ar (phosphate buffer, pH 6 and 8) from the µXRF analysis. The total acquisition time was 20 min per map. Data were processed using ESPRIT software. Energy calibration was performed prior to analysis.

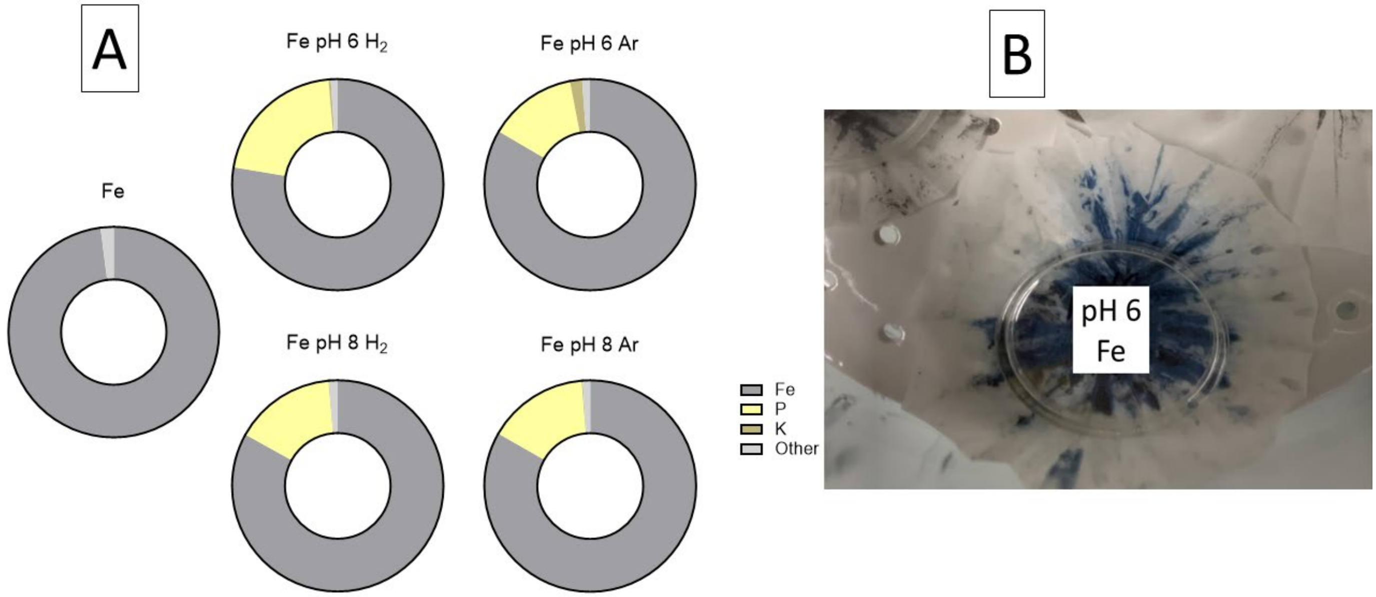


**Fig. S17.** (A) Ratios of Fe, P, K, and trace metals after 2 hour reaction in pH 6 or 8 phosphate buffer, under 5 bars of Ar or $H_2$ according to µXRF. (B) Picture of the $Fe^0$ pellet after 2 hour reaction in pH 6 phosphate buffer under 5 bars of Argon.

**Table S33.** Characterization of the pellet after a 2 h standard reaction under 5 bars of $H_2$ or Ar (phosphate buffer, pH 6 and 8) from the µXRF analysis.

| Sample | Element | Weight percent (mass %) | Weight percent (mass %) | Atomic percent (%) |
|---|---|---|---|---|
| **$Fe^0$ pH 6 Ar** | Al | 0.03 | 0.13 | 0.26 |
| | Cr | 0.01 | 0.06 | 0.06 |
| | Cu | 0.01 | 0.04 | 0.03 |
| | Fe | 19.57 | 90.06 | 84.51 |
| | K | 0.27 | 1.25 | 1.68 |
| | Mn | 0.05 | 0.25 | 0.24 |
| | Ni | 0.02 | 0.09 | 0.08 |
| | P | 1.62 | 7.45 | 12.61 |
| | Rh | 0 | 0 | 0 |
| | Zn | 0.14 | 0.66 | 0.53 |
| **$Fe^0$ pH 6 $H_2$** | Al | 0.03 | 0.15 | 0.28 |
| | Ca | 0 | 0.02 | 0.02 |
| | Cr | 0.01 | 0.05 | 0.05 |
| | Cu | 0.01 | 0.03 | 0.03 |
| | Fe | 15.6 | 85.66 | 77.41 |
| | K | 0.05 | 0.27 | 0.35 |
| | Mn | 0.04 | 0.2 | 0.18 |
| | Ni | 0 | 0.02 | 0.02 |
| | P | 2.37 | 13.02 | 21.22 |
| | Rh | 0 | 0 | 0 |
| | Zn | 0.11 | 0.58 | 0.45 |
| **$Fe^0$ pH 8 Ar** | Al | 0.02 | 0.12 | 0.23 |
| | Ca | 0 | 0.01 | 0.02 |
| | Cr | 0.01 | 0.06 | 0.06 |
| | Cu | 0.01 | 0.05 | 0.04 |
| | Fe | 18.42 | 89.49 | 83.22 |
| | K | 0.03 | 0.12 | 0.17 |
| | Mn | 0.05 | 0.24 | 0.23 |
| | Ni | 0 | 0.02 | 0.02 |
| | P | 1.91 | 9.26 | 15.53 |
| | Rh | 0 | 0 | 0 |
| | Zn | 0.13 | 0.63 | 0.5 |
| **$Fe^0$ pH 8 $H_2$** | Al | 0.03 | 0.15 | 0.3 |
| | Ca | 0 | 0.02 | 0.02 |
| | Cr | 0.01 | 0.06 | 0.06 |
| | Cu | 0.03 | 0.13 | 0.1 |
| | Fe | 18.77 | 89.27 | 82.96 |
| | K | 0.02 | 0.09 | 0.12 |
| | Mn | 0.05 | 0.25 | 0.23 |

| | | | | |
|---|---|---|---|---|
| | Ni | 0.01 | 0.03 | 0.02 |
| | P | 1.96 | 9.33 | 15.63 |
| | Rh | 0 | 0 | 0 |
| | Zn | 0.14 | 0.68 | 0.54 |
| **$Fe^0$** | Cr | 0.01 | 0.07 | 0.08 |
| | Cu | 0.01 | 0.08 | 0.07 |
| | Fe | 18.58 | 98.77 | 98.85 |
| | Mn | 0.07 | 0.38 | 0.39 |
| | P | 0 | 0.02 | 0.03 |
| | Rh | 0 | 0 | 0 |
| | Zn | 0.13 | 0.68 | 0.58 |
| | Cr | 0.01 | 0.07 | 0.08 |

**Pellet resuspension.** 4 mM FMN in pH 6 phosphate buffer was reduced with $Ni^0$ and $Fe^0$ at 40 °C for 2 h under an atmosphere of hydrogen or nitrogen. Following the reaction, the mixture was centrifuged at 4 °C and 13,000 rpm for 30 min. The supernatant was separated and measured with UV–Vis spectroscopy to ensure complete reduction.
The remaining pellet was gently washed with corresponding buffer without disturbing the solid phase. The wash solution was collected and analyzed by UV–Vis spectroscopy. The pellet was resuspended in buffer (1 mL, corresponding to the initial reaction volume). Resuspension was done by shaking and pipetting until complete dispersion was observed. The suspension was then centrifuged again at 4 °C and 13,000 rpm for 30 min.
An aliquot of the resulting supernatant was diluted 40-fold with buffer and analyzed by UV–Vis spectroscopy. In each step of measurement (supernatant, washing buffer, pellet) the sample was measured before and after reoxidation (see **Table S34 and Fig. S21.**).
The above mentioned protocol was also done on 0.1 mM riboflavin at pH6 phosphate buffer with $Fe^0$ under $N_2$, except these were not diluted after the reaction (see **Table S34.** and **Fig. S18.**)

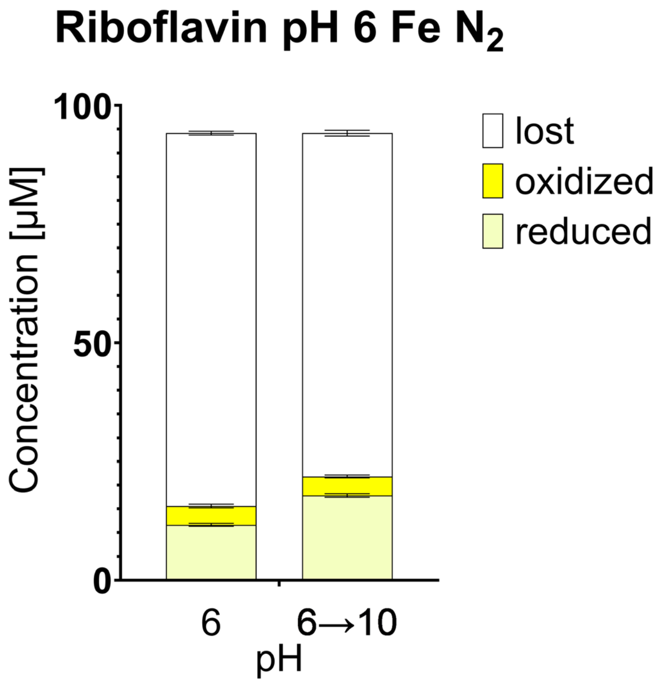


**Fig. S18.** Concentration of riboflavin in standard pH 6 samples with $Fe^0$, under 1 bar of $N_2$ for 2 hours. The pH of the samples was adjusted with 10 mM solution of KOH to pH 10 and the absorbance re-measured.

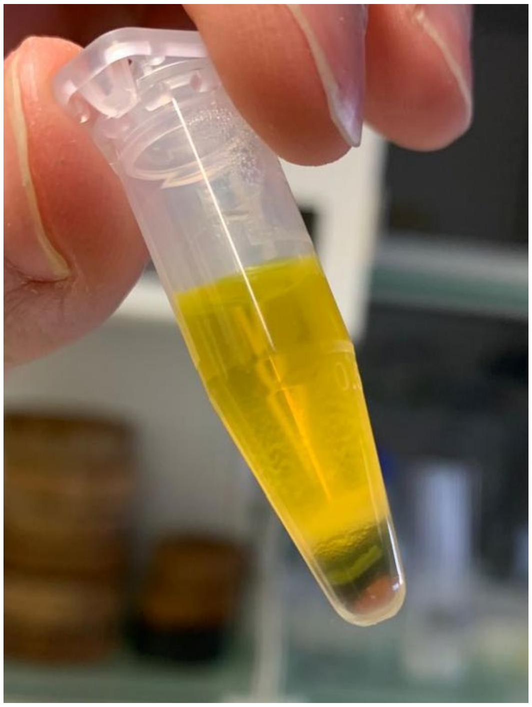

**Fig. S19.** Picture of 4 mM FMN with 18 mM $Fe^0$ after 2 hours at 40°C under $N_2$. The sample was transferred into a reaction tube centrifuged (4°C, 13000 rpm, 30 min) this separated four separate layers that are clearly visible at the bottom of the reaction tube: a white top layer, black, brown and red at the bottom. The red pellet disappeared after shaking the reaction tubetube and did not reprecipitate after another centrifugation.

**Table S34.** Post reaction absorbance values, pHs, and calculated concentration of the pellet and supernatant's samples reduced and oxidised proportions. The pH in the first column is that of the starting pH of the buffer, and in the pH on the second column is the pH of the sample after the 2h long reaction. Data regarding the corresponding supernatant is replicated here, regarding **Table S21**.

| Sample (metal +cofactor +pH +atmosphere +phase) | pH after reaction | Sample absorbance | Reoxidized sample Ab | Control absorbance | Reduced (µM) | Oxidised (µM) | Control (µM) | Change (µM) |
|---|---|---|---|---|---|---|---|---|
| **$Ni^0$ FMN pH 6 $N_2$ (supernatant)** | 5.9 | 0.17 | 0.65 | 1.20 | 1878.77 | 343.84 | 4138.73 | 1916.11 |
| | 5.9 | 0.18 | 0.67 | 1.20 | 1919.72 | 375.25 | 4138.73 | 1843.75 |
| | 5.9 | 0.21 | 0.60 | 1.20 | 1560.02 | 512.16 | 4138.73 | 2066.55 |
| **$Ni^0$ FMN pH 6 $H_2$ (supernatant)** | 5.89 | 0.12 | 1.14 | 1.16 | 4040.69 | -105.08 | 3992.25 | 56.64 |
| | 5.89 | 0.13 | 1.13 | 1.16 | 3961.69 | -55.60 | 3992.25 | 86.16 |
| | 5.89 | 0.15 | 1.07 | 1.16 | 3617.06 | 56.77 | 3992.25 | 318.42 |
| **$Fe^0$ FMN pH 6 $N_2$ (supernatant)** | 6.18 | 0.09 | 0.73 | 1.20 | 2495.17 | 2.87 | 4138.73 | 1640.69 |
| | 6.18 | 0.08 | 0.61 | 1.20 | 2093.65 | 12.73 | 4138.73 | 2032.35 |
| | 6.18 | 0.11 | 0.70 | 1.20 | 2324.04 | 94.16 | 4138.73 | 1720.52 |
| **$Fe^0$ FMN pH 6 $H_2$ (supernatant)** | 6.2 | 0.09 | 0.67 | 1.16 | 2318.64 | 1.95 | 3992.25 | 1671.66 |
| | 6.2 | 0.09 | 0.71 | 1.16 | 2435.18 | -0.81 | 3992.25 | 1557.89 |
| | 6.2 | 0.09 | 0.68 | 1.16 | 2350.80 | 2.65 | 3992.25 | 1638.80 |
| **$Ni^0$ FMN pH 6 $N_2$ (pellet)** | 5.9 | 1.08 | 1.79 | 1.20 | 680.44 | 90.59 | 4138.73 | 3367.69 |
| | 5.9 | 0.74 | 1.78 | 1.20 | 705.71 | 61.55 | 4138.73 | 3371.47 |
| | 5.9 | 0.67 | 1.72 | 1.20 | 684.87 | 55.69 | 4138.73 | 3398.16 |
| **$Ni^0$ FMN pH 6 $H_2$ (pellet)** | 5.89 | 0.02 | 0.15 | 1.16 | 10.51 | 2.11 | 3992.25 | 3979.63 |
| | 5.89 | 0.04 | 0.15 | 1.16 | 10.32 | 3.03 | 3992.25 | 3978.91 |
| | 5.89 | 0.04 | 0.14 | 1.16 | 8.55 | 3.08 | 3992.25 | 3980.62 |
| **$Fe^0$ FMN pH 6 $N_2$ (pellet)** | 6.18 | 0.76 | 1.15 | 1.20 | 432.88 | 63.99 | 4138.73 | 3641.85 |
| | 6.18 | 0.61 | 0.96 | 1.20 | 363.08 | 51.62 | 4138.73 | 3724.02 |
| | 6.18 | 0.72 | 1.08 | 1.20 | 404.39 | 61.07 | 4138.73 | 3673.27 |
| **$Fe^0$ FMN pH 6 $H_2$ (pellet)** | 6.2 | 0.74 | 1.14 | 1.16 | 429.80 | 61.99 | 3992.25 | 3500.46 |
| | 6.2 | 0.72 | 1.34 | 1.16 | 518.74 | 59.99 | 3992.25 | 3413.52 |
| | 6.2 | 0.84 | 1.32 | 1.16 | 498.76 | 70.56 | 3992.25 | 3422.93 |
| **$Fe^0$ Riboflavin pH 6 $N_2$ (supernatant)** | 6.2 | 0.06 | 0.20 | 1.21 | 11.99 | 3.58 | 94.18 | 78.61 |
| | 6.2 | 0.07 | 0.21 | 1.21 | 11.66 | 4.35 | 94.18 | 78.17 |
| | 6.2 | 0.06 | 0.19 | 1.21 | 11.37 | 3.83 | 94.18 | 78.98 |
| **$Fe^0$ Riboflavin pH 6 $N_2$ (pellet)** | 6.2 | 0.22 | 0.45 | 1.21 | 11.98 | 3.48 | 94.18 | 90.70 |
| | 6.2 | 0.08 | 0.43 | 1.21 | 11.66 | 4.25 | 94.18 | 89.93 |
| | 6.2 | 0.10 | 0.77 | 1.21 | 11.38 | 3.75 | 94.18 | 90.44 |
| **$Fe^0$ Riboflavin pH 6 $N_2$ (alkalified supernatant)** | 10 | 0.05 | 0.23 | 1.21 | 17.79 | 3.70 | 94.18 | 72.69 |
| | 10 | 0.06 | 0.24 | 1.21 | 18.26 | 4.29 | 94.18 | 71.64 |
| | 10 | 0.05 | 0.23 | 1.21 | 17.49 | 3.99 | 94.18 | 72.70 |

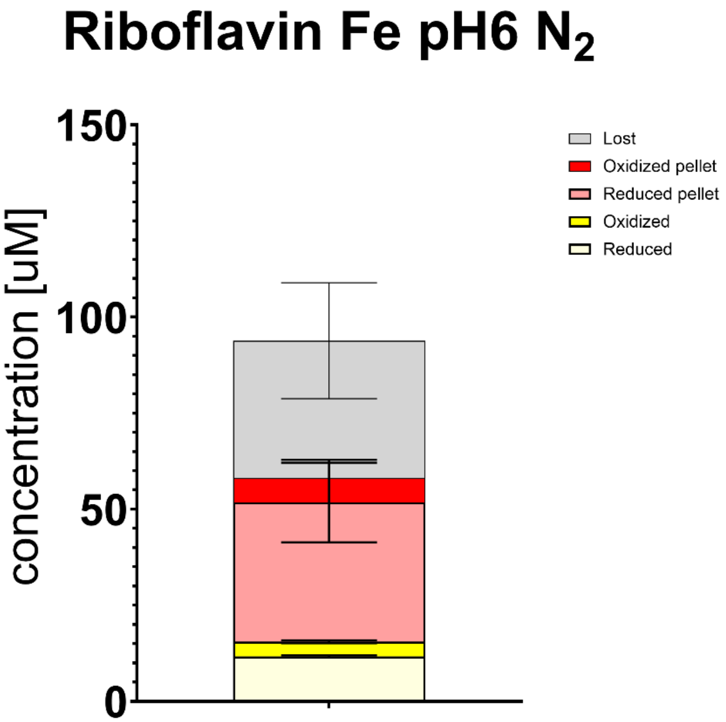


**Fig. S20.** Riboflavin ratios of reduced, oxidised, lost, reduced and oxidized pellet flavin concentrations after a two hour standard reaction at pH 6, $N_2$, and with $Fe^0$.

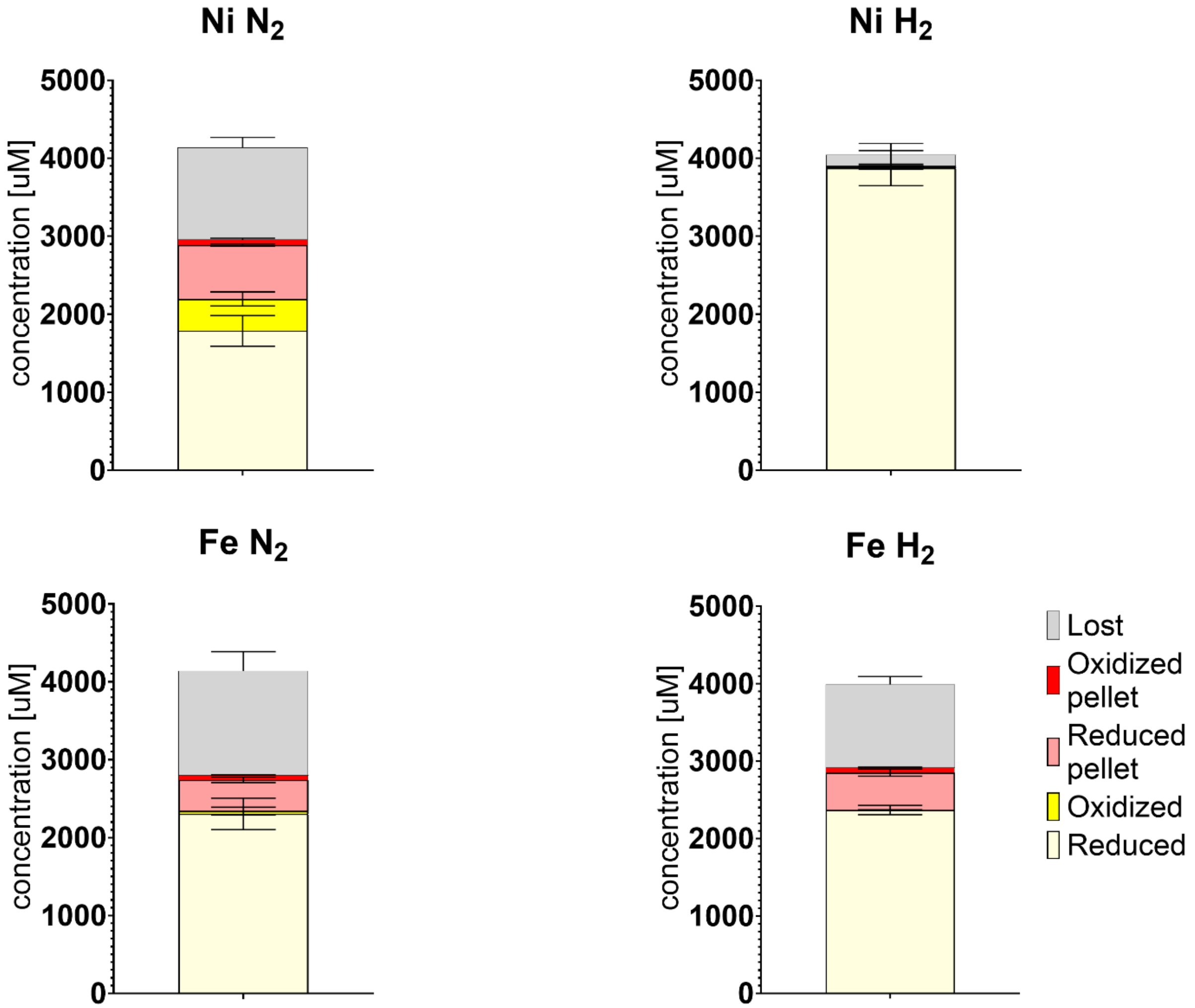


**Fig. S21.** FMN and $FMNH_2$ concentrations of the supernantant, and recovered from the pellet after a two hour standard reaction under all four different reaction conditions ($Fe^0$/$Ni^0$ and $H_2$/$N_2$).

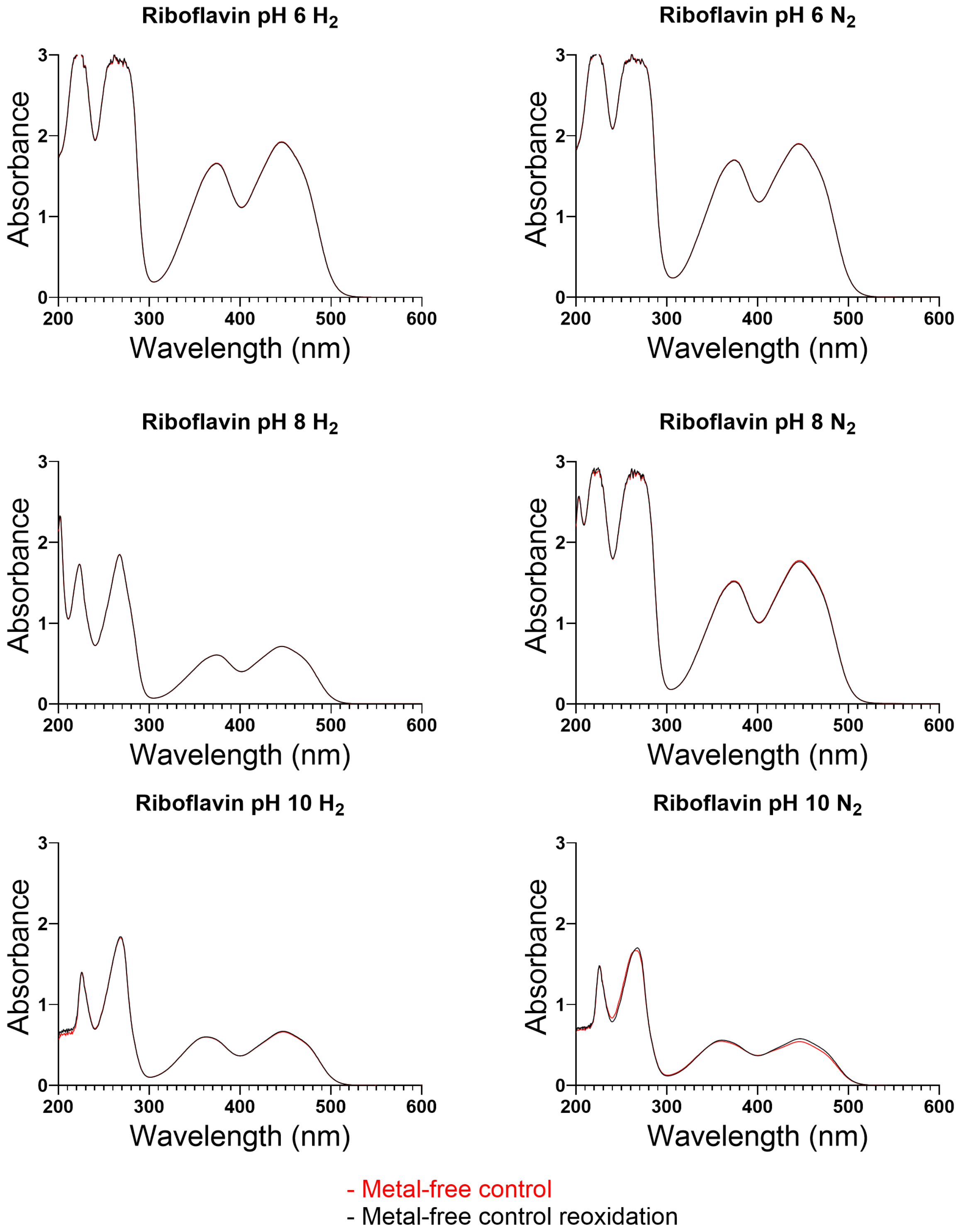


**Fig. S22.** Absorbance graph of metal free controls of riboflavin in their respective pHs after 2 hour reactions at 40°C. Red line is the control before, and black line is after exposure to oxygen.

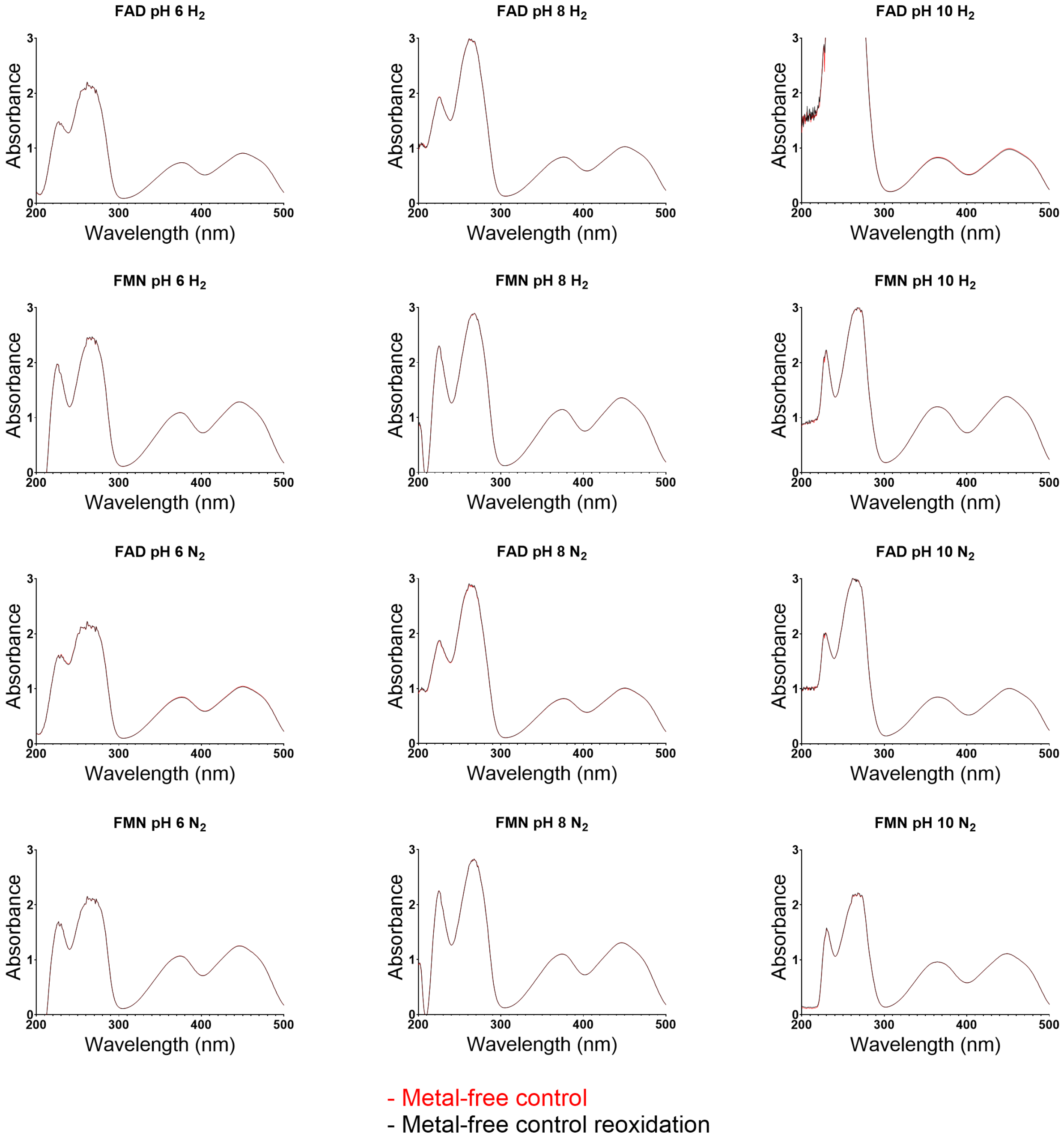


**Fig. S23.** Absorbance graph of metal free controls of FAD and FMN cofactors in their respective pHs after 2 hour reactions at 40°C. Red line is the control before, and black line is after exposure to oxygen.

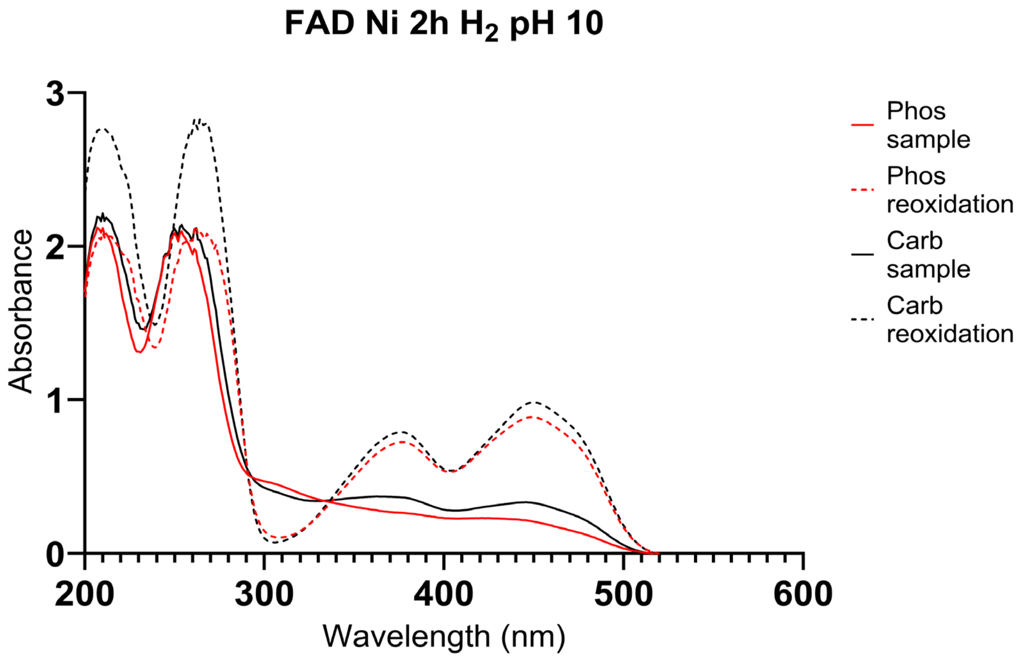


**Fig. S24.** Absorbance graph of pH 10 FAD in phosphate buffer (red line) and carbonate buffer (black line) with Ni, under $H_2$, after 2 hour reactions at 40°C. Dotted line is the sample before, and solid line is after exposure to oxygen
